# mTOR signalling, not clock gene rhythms, organises daily physiology

**DOI:** 10.64898/2026.08.28.747564

**Authors:** Aiwei Zeng, Andrei Mihut, Madhanagopal Anandapadamanaban, Alejandra Goity, Luíza Lane de Barros Dantas, Sew-Yeu Peak-Chew, Edward A. Hayter, Linda C. Andersson, Tom Smith, Giacomo Pasquini, Estere Seinkmane, Alessandra Stangherlin, Tomos E. Morgan, Nathan R. James, Charley Beresford, Jasmine Farnsworth, Joseph Menzies, Aymen al-Rawi, Liam J. Holt, Emmanuel Derivery, Rachel S. Edgar, Ralitsa R. Madsen, David A. Bechtold, Luis F. Larrondo, Antony N. Dodd, Jason Rihel, Gian Michele Ratto, Julie Williams, Peter Newham, Constanze Hilgendorf, Andrew D. Beale, Claudia Lodovichi, John S. O’Neill

**Affiliations:** MRC Laboratory of Molecular Biology, Cambridge, UK; Department of Infectious Disease, Imperial College London, London, UK; The Francis Crick Institute, London, UK; European Molecular Biology Laboratory, Heidelberg, Germany; Millennium Institute for Integrative Biology (iBio), Santiago, Chile; Facultad de Ciencias Biológicas, Pontificia Universidad Católica de Chile, Santiago, Chile; John Innes Centre, Norwich Research Park, Norwich, UK; Centre for Biological Timing, Faculty of Biology, Medicine and Health, University of Manchester, Manchester, UK; DMPK, Early Research and Development, Cardiovascular, Renal and Metabolism, BioPharmaceuticals R&D, AstraZeneca, Gothenburg, Sweden; Institute of Biophysics, CNR, Pisa, Italy; National Enterprise for Nanoscience and Nanotechnology (NEST), Scuola Normale Superiore, Pisa, Italy; University of Cologne, Faculty of Medicine and University Hospital Cologne, Cologne Excellence Cluster for Aging and Aging-Associated Diseases (CECAD), Institute for Mitochondrial Diseases and Ageing, Cologne, Germany; Institute for Systems Genetics, New York University Langone Health, New York, USA; MRC-Protein Phosphorylation and Ubiquitylation Unit, School of Life Sciences, University of Dundee, Dundee, UK; Cancer Research UK Scotland Institute, Glasgow, G61 1BD, United Kingdom; School of Cancer Sciences, University of Glasgow, Glasgow, G61 1QH, UK; Department of Cell and Developmental Biology, University College London, London, UK; Bioscience, Research and Early Development, Cardiovascular, Renal and Metabolism (CVRM), BioPharmaceuticals R&D, AstraZeneca, Gothenburg, Sweden; Clinical Pharmacology and Safety Sciences, AstraZeneca R&D, Cambridge, UK; Veneto Institute of Molecular Medicine, Padova Neuroscience Center, Padova, Italy

## Abstract

Daily rhythms in mammalian physiology are widely attributed to rhythmic transcription of “clock-controlled genes” driven by the circadian transcriptional/translational feedback loop (TTFL). Here we show that daily physiology can be uncoupled from the TTFL: inhibiting the mechanistic target of rapamycin (mTOR) abolishes most daily variation in cellular and organismal physiology, whilst clock protein rhythms persist. In cultured cells and mouse liver, functional rhythms are lost, including the proteome, RNA abundance, substrate utilisation and drug metabolism. In forebrain, most daily variation in protein abundance and phosphorylation is lost, yet locomotor activity and sleep-wake architecture are largely preserved in mice and zebrafish. PERIOD2 binds mTORC1 and limits its activity, providing a route by which clock proteins shape daily mTOR signalling. mTOR inhibition preferentially suppresses output over clock gene rhythms in *Arabidopsis* and *Neurospora*. We conclude that mTOR is a conserved conduit through which circadian and environmental timing impose daily organisation on physiology.

## Introduction

Most physiological processes follow a daily rhythm which arises from the interaction between external day/night cycles and endogenous timing (circadian) systems. Circadian rhythms persist under constant ambient/external conditions, *in vitro* and *in vivo*, and are observed at all levels of biological scale, from the molecular to the behavioural, optimising the use of bioenergetic resources by temporally compartmentalising metabolic processes according to the differing demands of day and night ^1–3^. Dysregulation of circadian physiology is intimately and causally linked with common age-related pathologies such as type II diabetes, neurodegeneration, cardiovascular disease and many forms of cancer ^4^. Understanding the factors that drive daily rhythms in healthy mammalian cells and tissues is essential to realising the potential of circadian medicine ^5^.

In mammals, the interaction between acute stimuli (e.g., light, feeding) and a hypothalamic master pacemaker determines the timing of systemic cues (e.g., body temperature, glucocorticoid and insulin signalling) that synchronise cell autonomous circadian rhythms throughout the body with each other and the external environment ^6–13^. These cell autonomous rhythms are thought to be driven by the activity of a core transcriptional/ translational feedback loop (TTFL) that is proposed to directly or indirectly drive circadian physiology by regulating the rhythmic transcription of "clock-controlled genes". In this circadian TTFL, transcription of *Period* and *Cryptochrome* genes is activated by a complex containing the BMAL1 transcription factor ^14^. The encoded PER and CRY proteins are translated, regulated post-translationally, and feed back to repress their own transcription, as well as that of other BMAL1-regulated genes, in ∼24h cycles. According to the canonical view, this and other auxiliary TTFLs drive rhythms in the transcription and translation of many proteins, imparting daily regulation to a wide array of physiological processes ^15,16^.

This hypothesis has been poorly supported by subsequent experimental evidence, however ^1^. The clock-controlled gene model makes a specific prediction: daily rhythms in physiology should be contingent upon rhythmic clock protein activity, and therefore inseparable from it except by disrupting the clock. To our knowledge, no mammalian gene has yet been shown to satisfy the full chain of inference the model requires: clock-protein-dependent rhythmic transcription, giving rise to rhythmic protein abundance, which is in turn rate-limiting for a rhythmic physiological output. Most rhythmically transcribed genes do not result in rhythmically abundant proteins, and *vice versa* ^17,18^, questioning the causal relationship between rhythmic transcription and daily physiology. Indeed, holding the cellular and nuclear abundance of the “core clock gene” PER constant over 24 hours has only modest effects on daily physiological rhythms ^19,20^. Rhythmic remodelling of the proteome is also inconsistent with the long half-life of most proteins (days) and the essential cellular requirement for protein homeostasis, which most cells expend much of their energy budget to maintain ^21–25^. Moreover, the number and extent of rhythmically abundant proteins is far more limited than rhythmic mRNA abundance would predict ^18,26–28^. For example, abundant complexes such as ribosomes have half-lives of many days, not hours ^26–28^, with no evidence that their abundance is rate-limiting for rhythmic translation in non-dividing cells ^29^.

If not transcriptional remodelling, what processes might instead give rise to daily physiology? Feeding is arguably the most potent input, outside of light entrainment of the suprachiasmatic nucleus (SCN), for entraining rhythms across the body, and is sufficient to synchronise daily rhythms *in vivo*, signalling through insulin/growth factor pathways and direct nutrient signalling (e.g., amino acid sensing) ^30–36^, of which the mechanistic target of rapamycin (mTOR) is a core downstream effector and a major input to the clock in cultured cells and tissues ^11^.

Essential and pan-eukaryotic, mTOR is a member of the atypical phosphatidylinositol 3-kinase-related kinase (PIKK) family of Ser/Thr kinases that participates in a wide variety of signalling pathways via its involvement in two complexes, mTORC1 and mTORC2. mTORC1 integrates extracellular stimuli (amino acid availability, energy charge, growth factor signalling) with intracellular signals of cellular state ^37^ to govern the catabolism/anabolism switch, stimulating energy intensive processes such as biogenesis of ribosomes and other complexes, with accompanying changes in macromolecular crowding and metabolic flux, whilst reconfiguring transcriptional programs and suppressing autophagy ^38^. mTORC2 has distinct roles including actin cytoskeletal organisation, membrane tension, and cell survival, but the copious crosstalk between mTORC1 and mTORC2 ensures that their activities are usually co-ordinately regulated ^39^.

mTOR activity is itself circadian regulated in cultured cells ^40,41^ and across multiple tissues *in vivo* ^41–45^. Several extracellular stimuli reset the cellular clockwork *via* mTOR or upstream insulin/insulin-like growth factor I signalling, such as class I PI3K and AKT ^11,46,47^. Numerous physiological processes downstream of mTOR are rhythmic, such as protein synthesis ^22,41,48,49^, metabolism ^50,51^, respiration ^52–54^, autophagy ^55^, crowding ^41^, and cytoskeletal dynamics ^56^. mTOR activity further drives rhythmic gene expression and metabolism in mouse liver ^57^, and mTORC1 reportedly interacts with known circadian regulators ^58^. Finally, hyperactivating mTORopathies already implicate mTOR as a critical regulator of daily physiology ^59,60^. Despite these links, whether mTOR activity itself drives daily physiology has not been directly tested.

Here we ask two questions. First, can daily physiology be separated from clock protein rhythms? Second, if the TTFL is not the proximal driver of daily physiology, what is? Using a reversible pharmacological approach to avoid the developmental and pleiotropic effects of genetic perturbation ^59,60^, we find that inhibition of mTOR abolishes most daily physiological variation in cultured cells, liver and forebrain while clock protein rhythms, locomotor activity and sleep–wake organisation persist. This tests a prediction we set out previously ^1^. We further show that mTOR is not a clock component but functions reciprocally as an input to the oscillator and as the principal conduit through which its timing, together with environmental cues, reaches physiology-a role conserved from plants and fungi to mammals. Our experiments establish that mTOR activity is necessary for most daily physiological rhythms, not that rhythmic mTOR activity alone is sufficient to generate them. What we test here is whether daily physiology depends on clock protein rhythms. It does not.

## Results

### Circadian cell physiology is uncoupled from clock protein rhythms

First, we tested the hypothesis that mTOR drives circadian cell biology *in vitro* using quiescent mouse lung PERIOD2::LUCIFERASE fibroblasts ^61^ (PER2::LUC), in which bioluminescence correlates with endogenous PER2 protein production ^62^. Mouse fibroblasts are a well-established model for circadian cell biology because they naturally quiesce through contact inhibition, with many cellular functions varying over the circadian cycle, such as protein synthesis, metabolism and migration ^22,53,56^.

mTORC1 activity shows cell-autonomous circadian regulation in cultured fibroblasts that reaches its maximum several hours after peak PER2::LUC (Fig 1A, Table S1), equivalent to the active phase *in vivo* ^41^. Treatment of cells with a saturating concentration of the selective mTORC1/2 active site inhibitor INK128 (1 µM) ^63,64^ had minimal effects on circadian phase of PER2::LUC rhythms (Fig S1A-B), consistent with previous reports ^45,65^. mTOR inhibition did, however, significantly decrease the baseline luminescence, consistent with the expected decrease in protein synthesis ^45,47^, and also reduced the amplitude of oscillation and lengthened the period by approximately 10% (Fig S1C-E)^45,47^. mTORC1 pathway engagement was verified by reduced S6K phosphorylation (Fig S1F). This demonstrates that mTOR activity influences, but is not strictly required for, circadian timekeeping in fibroblasts.

**Figure 1:**
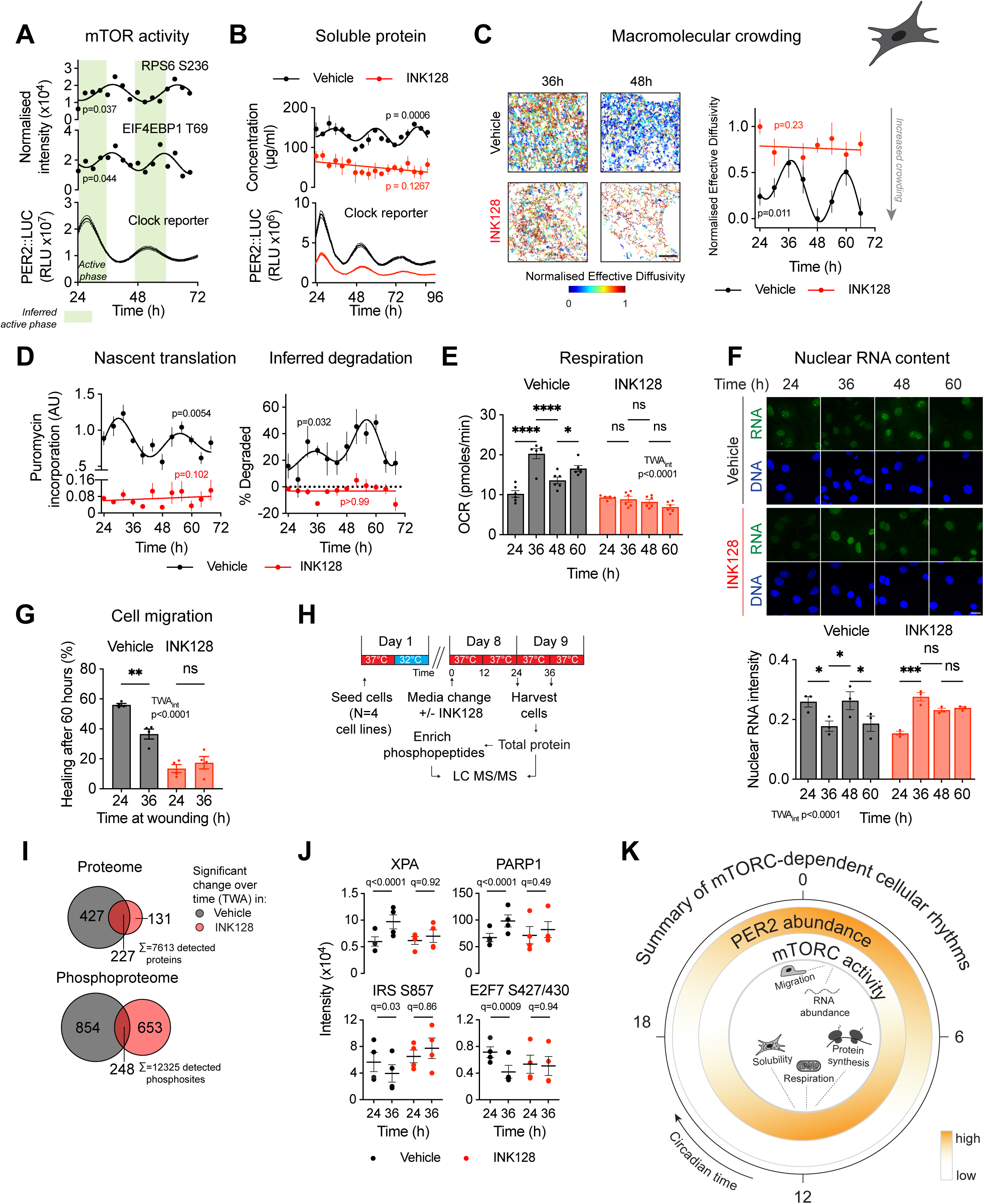
mTOR activity is required for circadian regulation of cell physiology A. Top: phosphorylation of mTOR target sites in PER2::LUC fibroblasts harvested every 3 hours across 3 days under constant conditions and analysed with mass spectrometry (p-value refers to the preferential fit to a straight line or damped cosine wave, null hypothesis = no rhythm). Bottom: PER2::LUC bioluminescence trace (n=6, mean ± SEM). Green shading indicates the presumed active phase equivalent in cells. B. Top: quantification of circadian variation in soluble protein abundance in PER2::LUC fibroblasts treated with INK128 (1 µM) and harvested every 4 hours for 3 days (n=3, mean ± SEM, p-value refers to the preferential fit to a straight line or damped cosine wave, null hypothesis = no rhythm. Black = vehicle, red = INK128). Bottom: bioluminescence trace of PER2::LUC (n=6, mean ± SEM). C. Circadian rhythms in effective diffusivity of fibroblasts stably expressing 40 nm cytosolic GEM-Sapphire (GEMs). Following INK128 treatment (100 nM), GEMs were imaged using spinning disk confocal microscopy every 6 hours for 2 days. Left: representative tracking images (scale bar 5 µm, colour scaled to the maximum and minimum effective diffusivity in the dataset). Right: quantification of effective diffusivity normalised to the dataset maximum and minimum (n>20 cells imaged per timepoint, p-value refers to preferential fit to a straight line or damped cosine wave, null hypothesis = no rhythm, mean ± SEM). D. Left: circadian rhythms in nascent protein translation in INK128-treated (1 µM) fibroblasts, pulsed with 10 µg/ml puromycin +/- 1 µM bortezomib (30 mins) every 6 hours for 2 days. Puromycin incorporation was assessed by Western blot and quantified as puromycin signal normalised to total protein. Right: % degradation inferred as the ratio of puromycin incorporation with versus without bortezomib (n=3, mean ± SEM, preferential fit to a straight line or damped cosine wave, null hypothesis = no rhythm). E. Daily variation in basal oxygen consumption rate in INK128-treated fibroblasts (1 µM) measured by Seahorse Mito Stress test every 12 hours across 2 days (n=6, mean ± SEM, two-way ANOVA (TWA) with Tukey’s MCT). F. Daily variation in nuclear RNA abundance in INK128-treated fibroblasts (1 µM). Cells were fixed and stained with SYTO RNASelect Green and DAPI every 12 hours for 2 days. Top: representative images (scale bar 25 µm). Bottom: quantification of nuclear RNA intensity (n=3 wells with >30 cells per field of view, TWA with Sidak’s MCT, mean ± SEM). G. Daily variation in residual wound area in fibroblasts treated with INK128, labelled with CellTracker Red, scratched 24 and 36 hours after INK128 treatment (10 nM), and allowed to heal for 60 hours. Residual wound area quantified as the percentage of original wound area repopulated with cells (n=4, TWA with Sidak’s MCT, mean ± SEM). H. Schematic of phosphoproteomics experiment: primary lung fibroblast cell lines from 4 PER2::LUC mice were harvested 24 and 36 hours after INK128 treatment (1 µM). Samples were enriched for phosphopeptides, each sample labelled with a different isobaric tandem mass tag and analysed by liquid chromatography tandem mass spectrometry (LC MS/MS). I. Overlap of proteins and phosphosites with significant time-dependent variation under vehicle or INK128 treatment (condition-specific two-way repeated-measures ANOVA with Benjamini-Krieger-Yekutieli (BKY) FDR correction, q≤0.05). J. Example proteins and phosphosites showing significant temporal variation in vehicle-treated cells (black, BKY q≤0.05) that is lost upon INK128 treatment (red, BKY q*≥*0.05). XPA = DNA repair protein complementing XP-A cells, IRS = insulin receptor substrate, PARP1 = poly[ADP-ribose] polymerase 1, E2F7 = E2F transcription factor 7. K. Summary of mTOR-dependent cellular rhythms across the circadian cycle. ns, not significant; \**p* < 0.05, \*\**p* < 0.01, \*\*\**p* < 0.001, \*\*\*\**p* < 0.0001

Macromolecular crowding (MMC) is largely regulated by mTORC1 ^41,66^ and is a fundamental aspect of cell physiology that affects diffusion rate, the stability of protein-protein interactions and many different enzyme activities ^67,68^. We first tested our prediction that mTOR drives circadian cell biology by investigating MMC rhythms in the presence and absence of mTOR inhibition using two independent methods: measurement of digitonin-soluble protein concentration ^69,70^ and the diffusion of 40 nm genetically encoded multimeric fluorescent nanoparticles (GEMs) ^66,71^. As predicted, mTOR inhibition completely abolished protein solubility rhythms, without affecting the phase of the parallel PER2::LUC clock reporter, whereas total cellular protein levels were not rhythmic in either condition (Fig 1B, S1G-H). Consistent with previous observations using membrane-permeable quantum dots ^41^, we observed a circadian rhythm in the diffusivity of GEMs in PER2::LUC fibroblasts (Fig 1C), which was abolished by mTOR inhibition. Taken together, this suggests that a daily rhythm in mTOR activity regulates daily rhythms in the solubility and sequestration of cytoplasmic macromolecules to internal membranes and/or membraneless compartments.

Protein homeostasis is maintained across the circadian cycle by coupled regulation of protein synthesis and degradation, minimising abundance fluctuations while facilitating proteome renewal and protein complex turnover ^3,22,40,41,48,72^. Since mTORC1 coordinates anabolic and catabolic balance ^37^, rhythmic mTORC1 activity should drive rhythms in proteome renewal. We tested this using a previously developed assay measuring puromycin incorporation ± the proteasome inhibitor bortezomib, enabling detection of nascent peptides that would otherwise be rapidly degraded ^22,73,74^ (Fig 1D, S1I). As previously observed ^22^, nascent protein production was circadian in control cells while net translation was not (Fig 1D, S1J). mTOR inhibition reduced overall translation rate whilst abolishing its rhythmic regulation; the puromycin ± bortezomib-derived index of degradative flux also lost rhythmicity (Fig 1D, S1I). Because INK128 also lowers the mean rate of nascent translation, we cannot formally exclude that a residual low-amplitude rhythm falls below our detection limit; we therefore interpret these data as loss of detectable circadian regulation of proteome renewal.

Given the substantial metabolic cost of protein synthesis, we next looked at mitochondrial metabolism, another process which shows both circadian rhythmicity and sensitivity to mTOR activity and MMC ^23,50,53,75^. Measuring oxygen consumption rate (OCR) every 12 hours for 2 days, we confirmed previously reported daily variation in basal OCR ^53^, that was elevated when nascent translation was highest and validated the prediction that daily variation would be abolished by INK128 (Fig 1E). OCR in INK128-treated cells was comparable to the daily nadir of vehicle-treated cells, consistent with our prediction that in quiescent cells, as in slow-growing yeast ^76^, rhythmic respiration is largely driven by the bioenergetic demands of rhythmic protein synthesis.

Circadian regulation of nuclear RNA transcription is central to the clock-controlled gene model, and AKT/mTOR signalling regulates RNA production at multiple levels, including Pol II-mediated transcription of protein-coding genes, and Pol I and Pol III-dependent transcription of ribosomal RNAs that constitute the majority of cellular RNA ^38,66,77^. Since mTOR inhibition does not affect the cellular capacity to maintain PER2 clock protein rhythms, we asked to what extent global circadian RNA metabolism depends on mTOR activity by quantifying total nuclear RNA abundance over time. Nuclear RNA levels indeed showed a circadian rhythm (Fig S1K,L) that was strongly disrupted under mTOR inhibition (Fig 1F), whereas the timing of PER2 rhythms was maintained (Fig S1A). This is consistent with our prediction that circadian RNA metabolism is primarily regulated by mTOR ^22^, rather than by direct TTFL activity.

Next, to test whether mTOR activity is required for temporal organisation of cellular response to challenge, we examined wound healing response in fibroblasts. As previously shown, in mesenchymal cells like fibroblasts, migratory capacity is circadian via rhythmic regulation of actin cytoskeletal dynamics ^22,56^. Actin polymerisation is sensitive to physiological changes in MMC ^78^, so we reasoned that rhythmic migration should not be observed in the absence of rhythmic MMC, when mTOR is inhibited. To preserve sufficient basal migratory capacity such that circadian disruption could be detected, we used sub-saturating concentrations of INK128 (Fig S1F). Under control conditions, we confirmed that fibroblasts wounded 24 hours after synchronisation healed faster than those wounded 36 hours post-synchronisation ^56^ (Fig 1G, S1M). Consistent with prediction, INK128 not only reduced overall healing rate but abolished the difference between timepoints (Fig 1G), strongly implicating mTOR activity in the circadian regulation of cell migration.

Globally, around 10% of detected fibroblast proteins and phosphopeptides typically vary over the circadian cycle ^21,22,56^. To determine what proportion of these circadian phosphorylations depend on mTOR activity, we performed quantitative 16-plex tandem mass tag (TMT) mass spectrometry on primary lung fibroblasts ± INK128 (Fig 1H, S1N, Table S1). Given that the circadian proteome and phosphoproteome of mouse fibroblasts are well characterised ^21,22,56^, we prioritised biological replicates over temporal resolution, harvesting cells at the circadian peak and trough of mTOR activity (24 and 36 hours after release into constant conditions, respectively) ^41^. After filtering for proteins and phosphosites present across all samples, we detected 7,613 proteins and 12,325 phosphosites. There was reduced overall abundance of 577 phosphosites under INK128 treatment, including well-characterised mTOR pathway components such as AKT1 S473 and RPS6 S244 (Fig S1O ^79,80^).

Consistent with previous work, 9% of proteins and 9% of phosphosites showed significant temporal variation in control cells (Fig 1I). Of these rhythmic proteins and phosphosites, significant difference between timepoints was lost in ∼65% and ∼77% respectively under mTOR inhibition (Fig 1I, S1P). This is comparable in scale with genetic TTFL disruption; for example, 75% of proteins lose rhythmicity in *Cryptochrome*-deficient fibroblasts ^21^. Interestingly, a subset of proteins and phosphosites gained temporal variation under INK128, suggesting that mTOR activity may normally suppress their rhythmicity (Fig 1I).

Gene ontology analysis of the time-significant mTOR-dependent proteins revealed an enrichment for terms relating to DNA binding and transcriptional regulation, with time-significant mTOR-dependent phosphosites additionally enriched for terms relating to protein binding and kinase activity (Fig S1Q). These included proteins such as DNA repair proteins XPA and Poly-ADP-ribosyltransferase 1 (PARP1), insulin pathway proteins IRS1, IRS2 and INSR, and transcription factor E2F7 (Fig 1J). Because TTFL rhythms continue to function under INK128 treatment, it seems plausible that most circadian changes in gene expression that have been attributed directly to TTFL activity are in fact mTOR-dependent, consistent with recent findings in mouse liver ^57^.

Taken together, our results in cultured fibroblasts reveal that circadian rhythms in mTOR activity occur cell-autonomously. mTOR is not strictly required for circadian timekeeping, yet is necessary for the circadian regulation of multiple fundamental cellular processes including proteome renewal, signalling, bulk RNA metabolism and transcription, as well as overt rhythms in cell function such as migration (Fig 1K).

### mTOR mediates signals to the cellular circadian clock

Since mTOR is a key effector of the insulin/IGF signalling pathway ^38^, we predicted that activation of mTOR alone would be sufficient to reset the cellular circadian clock. We applied MHY1485, an indirect mTOR activator ^81^ (Fig 2A, S2A). MHY1485 strongly reset circadian PER2::LUC rhythms (Fig 2A,B, S2A) without inducing PER2 or affecting the period of oscillation (Fig 2C) and with a modestly increased amplitude (Fig 2D). That clock resetting does not necessarily require PER2 induction suggests the involvement of other factors, e.g. PER1, or that the resetting mechanism involves post-translational modification of PER that normally accompanies, but does not require, changes in PER abundance.

**Figure 2:**
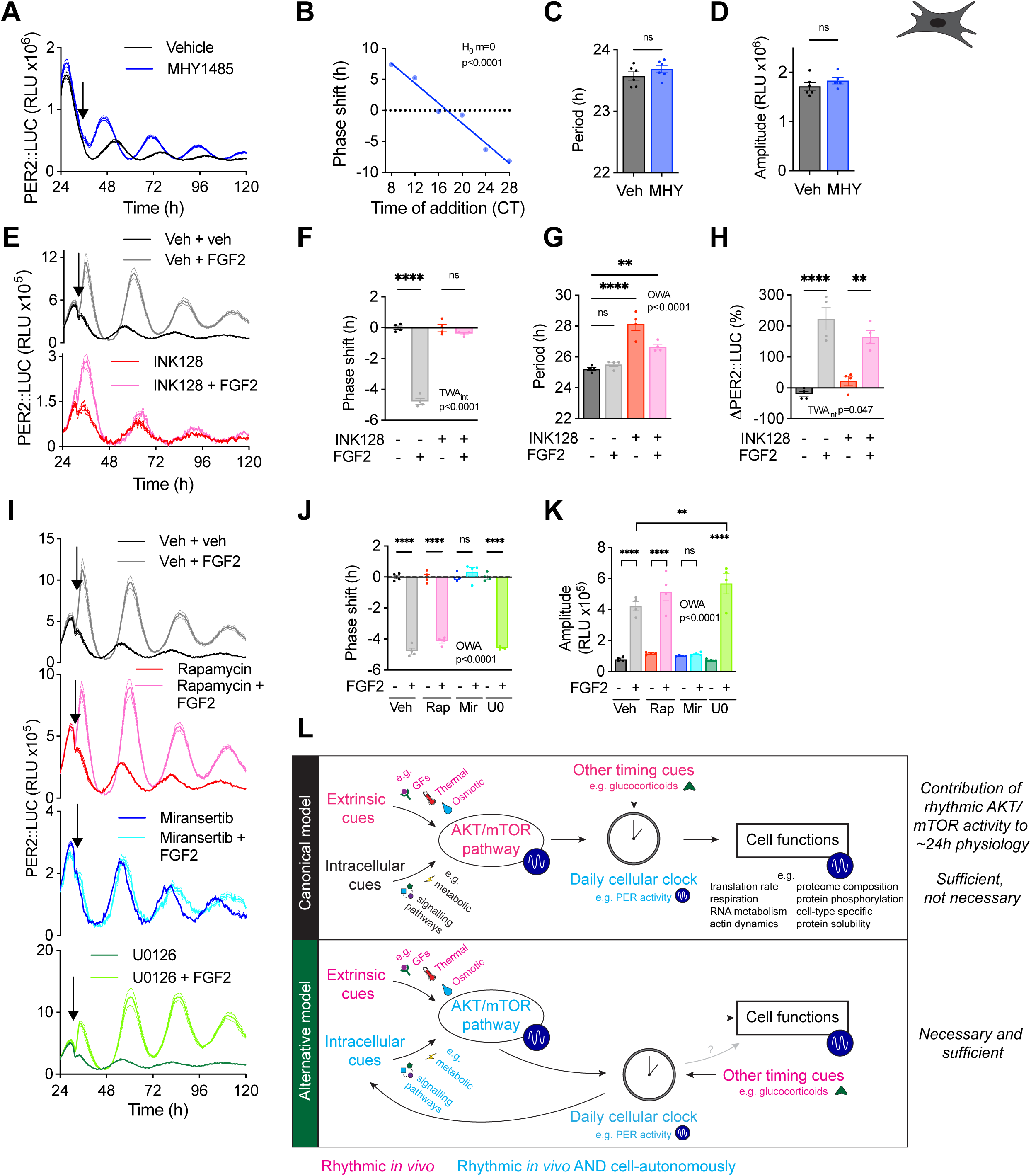
mTOR regulates inputs to the circadian timekeeping mechanism A. Representative bioluminescent trace of PER2::LUCIFERASE fibroblasts treated with 1 µM mTOR activator MHY1485 at 36 hours after release into constant conditions, as indicated by arrow, and maintained throughout (n=6, mean ± SEM, RLU = relative light units). B. Phase response curve of PER2::LUC fibroblasts treated with MHY1485 every 4 hours across 1 circadian cycle (mean ± SEM, null hypothesis: gradient = 0). Circadian time 0 (CT) is defined as the time of release into constant conditions. Phase shifts were calculated relative to vehicle treatment. C. Period of MHY1485 (MHY) treated cells compared to control vehicle (Veh) at T24 (two-tailed t-test, mean ± SEM). D. Amplitude of MHY1485 (MHY) treated cells compared to control vehicle (Veh) at T0 (two-tailed t-test, mean ± SEM). E. Pre-treatment of PER2::LUC fibroblasts with 1 µM INK128 (added at T0) blocks phase shift induced by addition of FGF2 36 hours after release into constant conditions, indicated by arrow (n=4, mean ± SEM). F. Phase shift for Fig 2E, calculated relative to untreated conditions (mean ± SEM, two-way ANOVA (TWA) with Sidak’s multiple comparisons test (MCT)) G. Period for Fig 2E (mean ± SEM, OWA with Sidak’s MCT). H. % change in PER2::LUC signal intensity (ΔPER2::LUC) for Fig 2E (mean ± SEM, TWA with Sidak’s MCT). I. Representative PER2::LUC trace showing that simultaneous mTORC1 inhibition alone (1 µM rapamycin) or 10 µM MEK1/2 inhibitor U0126 does not abolish the phase shift effect of FGF2, but AKT inhibition (1 µM miransertib) does (n=4, mean ± SEM). J. Phase shift for Fig 2I (mean ± SEM, OWA with Sidak’s MCT). K. Amplitude for Fig 2I (mean ± SEM, OWA with Sidak’s MCT). L. Canonical vs alternative model for the role of AKT/mTOR signalling in circadian physiology. In the canonical model, rhythmic AKT/mTOR activity, driven by extrinsic cues and intracellular pathways, feeds into the daily cellular clock, which in turn drives rhythmic cell functions. In this model, rhythmic AKT/mTOR activity is sufficient but not necessary for ∼24h physiology. In the alternative model, rhythmic AKT/mTOR activity acts both through and in parallel with the cellular clock to drive cell functions, and is necessary and sufficient for daily physiological rhythms. Pink: rhythmic *in vivo*, blue: rhythmic *in vivo* and cell autonomously.

Growth factors activate mTOR signalling physiologically ^37,82^. Application of fibroblast growth factor 2 (FGF2) to PER2::LUC fibroblasts acutely induced PER2, increased amplitude, and elicited strong circadian resetting, consistent with prior observations that employed insulin and TGFβ^11,47,83^ (Fig 2E-H). As expected, prior mTOR inhibition with INK128 abolished the FGF2-induced phase shift and amplitude increase but did not fully block the period increase or induction of PER2, suggesting that PER2 induction is not mTOR-dependent, while resetting is.

To identify which arm of the pathway mediates clock resetting, we tested various selective inhibitors (Fig 2I-K, S2A). Rapamycin, an allosteric partial mTORC1 inhibitor ^84^ had no effect on FGF2-induced phase shifts, PER2-induction or amplitude increase. In contrast, miransertib, a pan-AKT inhibitor acting upstream of mTOR ^85^, completely abolished both the FGF2-induced phase shift and amplitude increase, implying that upstream AKT-dependent signalling is required, though this does not exclude additional AKT-independent roles for mTORC2. FGF signalling also activates the RAS-MAPK pathway ^86,87^; however, addition of MEK1/2 inhibitor U0126 did not block FGF2-induced phase shifts but accentuated the amplitude response (Fig 2I-K). Taken together, these findings indicate that growth factor signals to the clock are primarily transduced by AKT-dependent signalling rather than MEK-ERK.

Acute increases in macromolecular crowding *via* osmolarity or temperature have also been reported to reset circadian rhythms *via* mTOR ^47,65,88,89^, therefore we applied a +100 mOsm/L hyperosmotic stimulus to PER2::LUC fibroblasts. Again, we found that inhibition of mTOR by INK128 attenuated hyperosmolarity-induced phase shifts (Fig S2B-D). In contrast, phase shifts elicited by other stimuli that are not known to signal *via* mTOR (e.g. glucocorticoids ^6^, cyclic AMP ^90^) ^91^were not blocked by mTOR inhibition (Fig S2E-G). Consistent with serum acting only in part through PI3K/AKT ^91^, INK128 partially disrupted serum-induced phase resetting and abolished amplitude increase after serum stimulation (Fig S2E-G).

mTOR signalling therefore mediates several, but not all, inputs to the cellular circadian clock, including extrinsic cues such as growth factor signalling and thermo-osmotic perturbations ^47^, as well as cell-intrinsic signals (Fig 2L) ^40,41^. In the canonical model for circadian regulation of cellular function, these mTOR-dependent signalling cues are communicated to the cellular clock which drives rhythmic outputs such as translation ^48^ and metabolism ^53^ via clock-controlled genes (Fig 2L, upper panel). According to this paradigm, mTOR may alter the timing of circadian cell physiology but is not necessary for it. However, our observations show mTOR regulates many biological processes, is required for circadian cell physiology, and also signals to the cellular clock (Fig 1, 2). This suggests a simpler alternative model where AKT/mTOR signalling reciprocally regulates the cellular clock and thereby drives most rhythmic cellular functions (Fig 2L, lower panel) ^43^. This alternative model could also explain why many daily physiological rhythms persist in mice where cellular clock function has been genetically compromised^18,20,92^.

### PER2 regulates mTORC1

Given that mTOR signalling to the cellular TTFL normally involves changes in PER protein activity ^11^, we next asked how the TTFL signals back to mTOR. PER2 has been proposed to regulate mTORC1 through direct interaction ^58^, raising the question of whether it acts as a circadian regulator of mTORC1 activity.

First, we examined whether endogenous PER2 interacts with native mTORC1 components using human PER2-HaloTag (PER2-HALO) knock-in U-2 OS cells ^47^. Consistent with previous over-expression experiments ^58^, endogenous PER2 co-immunoprecipitated with mTORC1 component RAPTOR (Fig 3A). Mass spectrometry of PER2-HALO immunoprecipitates revealed further interaction with mTORC1 components (Fig 3B, Fig S3A, Table S1) including mTOR and mLST8, as well as upstream regulators of mTORC1, such as components of the GATOR2 and TSC complexes ^93,94^. In contrast, neither the mTORC2-defining subunit RICTOR nor components of the Ragulator (LAMTOR) complex, which anchors mTORC1 to the lysosome for activation ^95^, were enriched, suggesting that PER2 associates selectively with mTORC1 and preferentially with a cytoplasmic, non-lysosomal pool of mTORC1.

**Figure 3:**
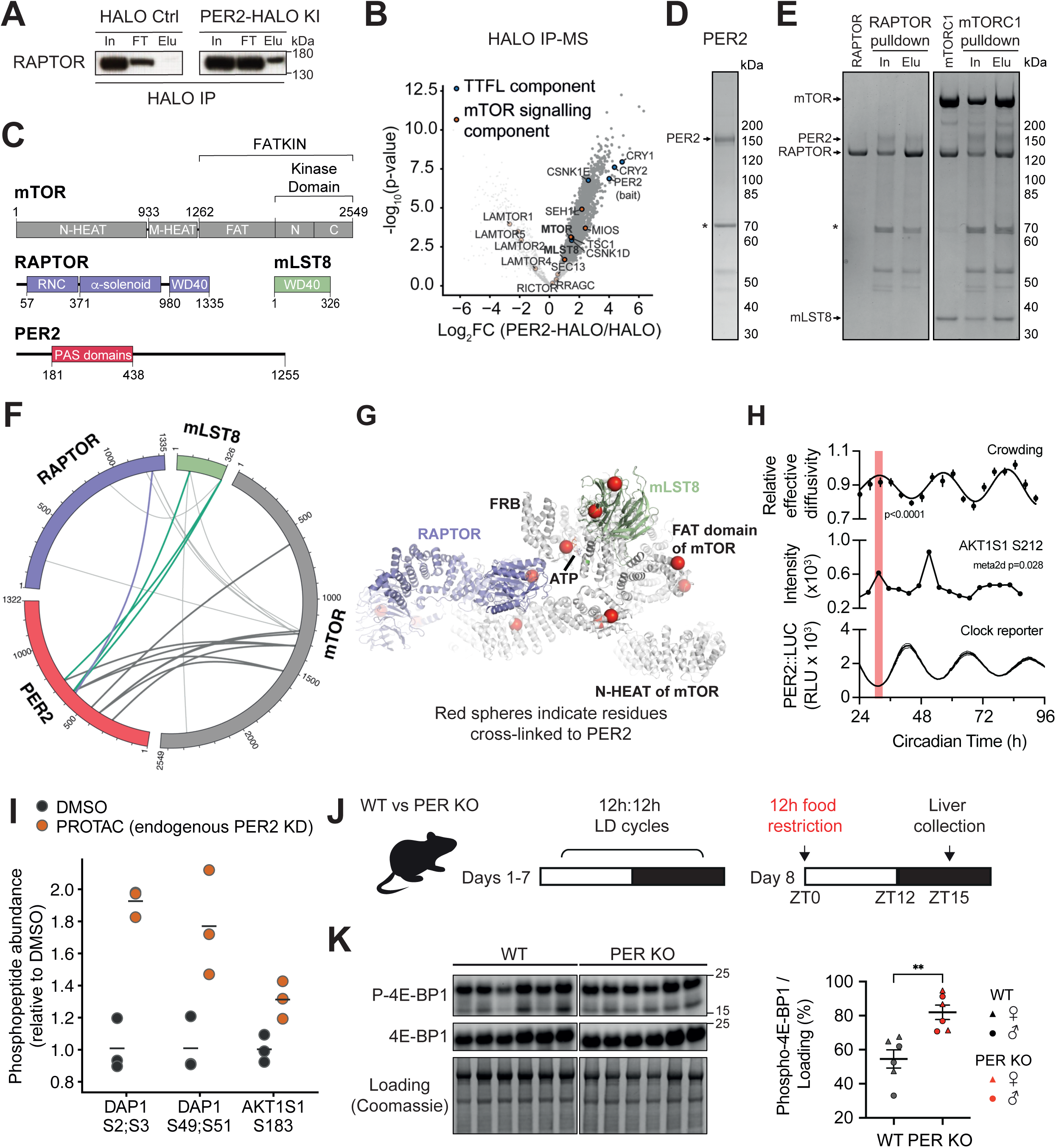
PER2 modulates the activity of mTORC1 A. HaloTag immunoprecipitation followed by immunoblotting for RAPTOR in U-2 OS cells overexpressing HaloTag or expressing endogenous PER2-HaloTag. IN, input; FT, flow-through; Elu, HaloTrap bead eluate. B. Volcano plot of fold change and p-value of proteins enriched in endogenous PER2-HaloTag immunoprecipitations relative to HaloTag controls, identified by LC–MS/MS. TTFL components and mTOR signalling components are highlighted. C. Schematic representation of mTORC1 subunits and PER2 constructs used for subsequent expression and purification. D. Purified full-length human PER2 protein. Asterisks denote co-purifying protein HSP70. E. In vitro Strep-tag pulldown assays assessing the interaction between purified human RAPTOR or mTORC1 with purified human, full-length PER2. Input and eluate fractions were analysed by Coomassie-stained SDS–PAGE. F. Circular plot of the observed cross-links between PER2 and mTORC1 subunits: mTOR, RAPTOR and mLST8. G. Summary of cross-links between PER2 and mTORC1 mapped on the structure of mTORC1 (PDB: 6BCX) shown in Fig 3F and Fig S3C. For clarity, the cross-linking residues are shown for only one protomer of the dimeric mTORC1 complex. H. Rhythmicity in U-2 OS cells of (top) relative effective diffusivity of U-2 OS cells (n=10 per timepoint) stably expressing 40 nm cytosolic GEM-Sapphire (GEMs); (middle) AKT1S1 S212 phosphorylation (n=1 per timepoint); and (bottom) PER2::LUC (n=6). For effective diffusivity, p-value refers to preferential fit to a straight line or damped cosine wave, null hypothesis = no rhythm. For AKT1S1 S212 phosphorylation, data were fit with MetaCycle. I. Fold change of select *in vitro* validated mTORC1 substrate sites identified by LC-MS/MS in U-2 OS cells expressing endogenous PER2-HaloTag following acute PER2 depletion by HaloPROTAC3 treatment, quantified relative to DMSO control. J. Schematic of experimental design. Wild-type and *Per1^-/-^, Per2^-/-^* adult mice (N=6 per genotype [3 male, 3 female]) were entrained to 12h:12h light:dark cycles (LD). On day 8 at lights on, food was restricted for 12 hours, then food was returned for 3 hours after lights off, and livers collected and blotted for mTORC1 activity. K. Assessment of hepatic mTORC1 activity by immunoblotting for phosphorylation of 4E-BP1, quantified relative to total protein loading (right). (Two-way ANOVA with genotype and sex as factors: genotype p = 0.0045, sex p = 0.302, genotype × sex p = 0.673. N = 6 per genotype [3 male,3 female], mean ± SEM).

To investigate the structural basis of the interaction *in vitro*, we independently purified full-length human PER2 and mTORC1 from Expi293 cells ^96^ (Fig 3C-E). In Strep-tag pull-down assays using Strep-tagged RAPTOR or mTORC1 ^96^ as bait, untagged PER2 was retained by both RAPTOR and mTORC1, but showed little or no retention with reconstituted mTORC2 (Fig 3E, Fig S3B). We then performed cross-linking mass spectrometry on purified PER2 and mTORC1 complex to gain further insight into their specific interaction sites. Most PER2 crosslinks formed with mTOR and mLST8, with few crosslinks to RAPTOR (Fig 3F, Fig S3C), indicating that PER2 contacts the mTOR-mLST8 core of the complex rather than the RAPTOR scaffold. Mapping cross-linked PER2 residues onto the mTORC1 cryo-EM structure (PDB ID: 6BCX) revealed contacts with the FAT (FRAP, ATM and TRRAP) domain (mTOR residue 1531) and the ATP-binding loop (mTOR residue 2165) (Fig 3G). Notably, DEPTOR, an endogenous allosteric inhibitor of mTORC1 activity, binds within the same region of the FAT domain (aa_mTOR_ 1525-1578) ^97,98^. While the cross-linking data do not establish a definitive binding mode, the proximity of the PER2 cross-links to this regulatory interface suggests that PER2 may act on the catalytic subunit itself, in addition to the TSC1-recruitment mechanism proposed previously ^58^.

We next asked whether this interaction has functional consequences for mTORC1 output. In U-2 OS cells under constant conditions, as with mouse fibroblasts, we observed a temporal relationship between mTORC1 activity (as measured both by phosphorylation of an mTORC1 substrate site, AKT1S1 (PRAS40) S212 ^99^, and by macromolecular crowding via GEMs ^66^) and circadian time such that mTORC1 activity is high when PER2 abundance is low (Fig 3H), in line with previous (overexpression) studies ^58^. To further examine this relationship, we acutely depleted endogenous PER2 protein using the HALO-PROTAC system ^47^ and quantified changes in the phosphoproteome (Table S1). In agreement with the above, we observed significantly increased phosphorylation of *in vitro* validated mTORC1 substrate sites ^99,100^ AKT1S1 (PRAS40) S183 and DAP1 S2;S3 and S49;S51 (Fig 3I, Fig S3D) upon 24h PER2 depletion. Phosphorylation of AKT1S1 S183 by mTORC1 relieves AKT1S1-mediated competition for the RAPTOR substrate-docking site ^99^ whereas S3 and S51 on the autophagy regulator DAP1 are directly phosphorylated by mTOR upon activation ^101^. Increased phosphorylation at these mechanistically distinct nodes after 24h PER2 depletion, such as substrate competition at the complex itself and downstream autophagy control, suggests that PER2 negatively regulates mTORC1 activity.

Daily regulation of mTORC1 and PER2 by extracellular cues is partially understood^11,13,41^, but the basis for cell-intrinsic rhythmicity in mTORC1 is not. Our observations suggest the hypothesis that a combination of cell-intrinsic and extracellular signals normally drive PER2 rhythms, and this is an important factor sculpting daily changes in mTORC1 activity. To test this, we subjected wild-type or *Per1/2*-deficient mice to a fasting-refeeding protocol (Fig 3J) designed to maximally stimulate hepatic AKT/mTOR signalling ^102^ at the onset of the active phase. Livers were collected 3 h after refeeding at the expected peak of insulin-driven mTORC1 activity. Consistent with our findings in cells, hepatic mTORC1 activity, assayed by 4E-BP1 phosphorylation, was elevated in the *Per1/2* knockout mice compared with wild-type controls (Fig 3K). Together, these data show that PER2 directly associates with mTORC1 and likely regulates its activity as PER levels vary across the circadian cycle in a fashion that may function (in part) by attenuating mTORC1 activation around the rest/activity transition. This provides a putative mechanistic basis by which the cellular circadian clockwork contributes to daily cycles of mTOR signalling and supports the hypothesis that mTOR is a major circadian output pathway.

### mTOR inhibition uncouples daily hepatic physiology from mouse behaviour

Next, we investigated the contribution of mTOR signalling to daily physiological rhythms *in vivo*. We took advantage of the blood-brain-barrier-permeability of INK128 ^63^ to transiently suppress mTOR activity whilst avoiding the development and/or deleterious effects of genetically perturbing mTOR signalling, e.g., *Mtor* knockout mice are embryonic lethal ^103^. Three days of 25 µM INK128 (∼1 mg/kg) provided *ad libitum* to wild-type mice in drinking water was sufficient to reduce phosphorylation of RPS6 and AKT by ∼75% and ∼60% respectively in the liver (Fig S4A), without adversely affecting body weight after two weeks of treatment (Fig S4B).

Given the measured hepatic half-life of INK128 (∼103 min, consistent with its reported serum half-life ^63^) (Fig S4C), and daily variation in fluid intake, the degree of mTOR inhibition likely fluctuates over the 24h cycle. However, estimated serum levels are not expected to drop low enough to appreciably relieve inhibition (Fig S4C), and the phosphoproteomic data confirm sustained pathway inhibition across the day (Fig S4D). INK128-treated mice showed no change in free-running period or total activity counts either under normal 12h:12h light:dark (LD) cycles or under constant darkness (Fig S4E-H). We did, however, note a subtle but significant increase in the consolidation of locomotor activity at the beginning of subjective night, in agreement with recent reports that mTOR activity outside the SCN can modulate the daily organisation of locomotor activity ^45,47^ (Fig S4H).

To gain initial insight into the role of mTOR in the daily organisation of murine physiology, we performed quantitative (phospho)proteomics in the liver. Vehicle or INK128 was provided *ad libitum* in drinking water for 3 days, then tissues were collected every 6h across a standard 12h:12h light:dark (LD) cycle (n=4 mice) (Fig 4A, Table S1). Due to extensive literature on the daily or circadian (phospho)proteome of mouse liver and other tissues ^92,104–108^, our experiment design and analyses focused on daily variation (>10% change in abundance) that is both functionally and statistically significant ^21^.

**Figure 4:**
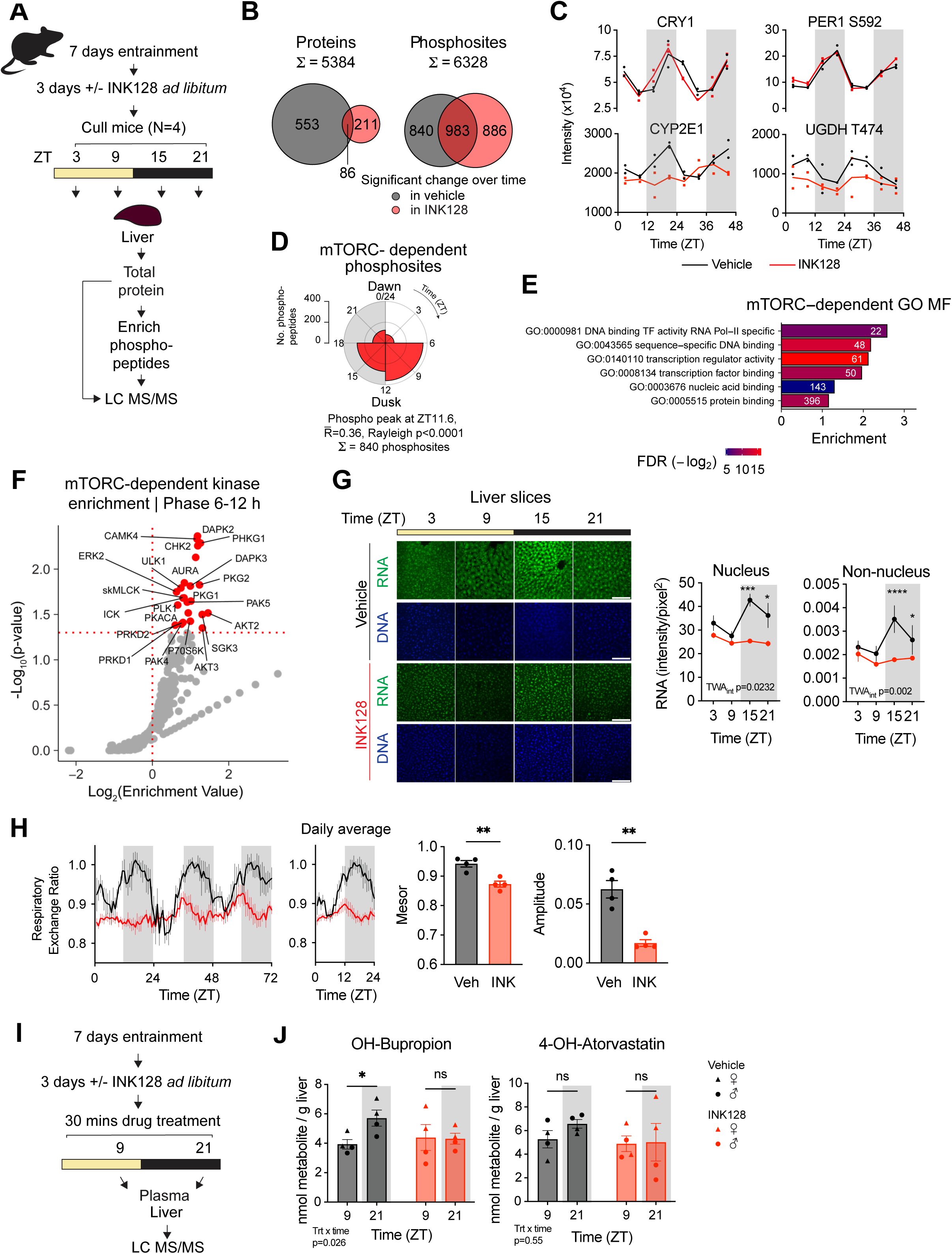
Partial mTOR inhibition severely attenuates daily physiology but not locomotor activity A. Schematic of quantitative (phospho)proteomics experiment *in vivo*: Mice were entrained for 7 days under 12h:12h light:dark cycles (LD), then treated with vehicle or 25 µM INK128 given *ad libitum* in drinking water. Livers were collected every 6 hours across 1 LD cycle (N=4 mice per timepoint and condition), and proteome and phosphoproteome analysed by LC MS/MS. ZT, zeitgeber time (ZT0 = lights on). B. Overlaps between proteins and phosphosites showing significant daily variation under vehicle or INK128 treatment (BioDare2 eJTK cycle Benjamini-Hochberg FDR≤0.05). C. Top: abundance of example core clock proteins and phosphosites. Bottom: example proteins and phosphosites (line = mean). Data plotted across 2 days for visualisation purposes but for rhythmicity analysis, timepoints were handled as 1 day. Shaded boxes represent time of lights off. D. Rose plot of peak phases of mTOR-dependent phosphosites (rhythmic under vehicle but not under INK128), binned into 6 h windows. Mean peak phase, mean resultant length (R̄) and Rayleigh test p-value are shown. E. Ranked GO analysis for molecular function (MF) performed using GOrilla with proteins showing time-of-day variation in phosphorylation in vehicle but not INK128 treatment, against a background of all proteins. Terms were simplified using REViGO. Number of genes is plotted for each term, coloured by FDR q-value and labelled with fold enrichment. F. Motif-based kinase enrichment analysis of phosphosites with time-of-day variation that peaked between ZT6-18 and were lost after INK128 treatment when compared to a background of all detected phosphosites (enriched kinases in red with p < 0.05). G. Daily variation in RNA abundance in liver slices from mice treated with vehicle or INK128 (25 µM) for 3 days, collected every 6 hours across 1 LD cycle. Tissues were fixed and stained with SYTO RNASelect Green. Left: representative images (scale bar 100 µm). Right: quantification of nuclear RNA intensity and cytoplasmic (non-nuclear) RNA intensity, normalised to pixel area (N=4, TWA with Sidak’s MCT, mean ± SEM). H. Respiratory exchange ratio for mice treated with vehicle or INK128 (25 µM) in LD (N=4, mean ± SEM). Mice were treated with INK128 for 3 days prior to experiment start. Left: grouped traces across 3 days of recording (mean± SEM). Middle left: average of 3 days’ recording (mean± SEM). Middle right: mesor of oscillation (mean ± SEM, t-test). Right: amplitude of oscillation (mean ± SEM, t-test). I. Schematic of experiment: mice were entrained for 7 days LD and treated with vehicle or INK128 (25 µM) for 3 days. At ZT9 or 21, mice (N=4 per group) were simultaneously intraperitoneally injected with bupropion and atorvastatin. 30 minutes later, blood plasma and liver were collected for LC-MS/MS analysis. J. Hepatic metabolite levels of OH-bupropion (metabolite of bupropion, left) and 4-OH-atorvastatin (metabolite of atorvastatin, right) normalised to parent drug abundance and input liver mass (N=4 [2 male, 2 female], mean ± SEM, multiple linear regression with treatment, time and treatment × time as factors and sex as a covariate, with Sidak’s MCT).

After excluding proteins/phosphosites that were not present across all samples, we quantified 5,384 proteins and 6,328 phosphosites (Fig 4B). Using eJTK Cycle ^109^ with Benjamini-Hochberg (BH)-corrected p<0.05, we detected significant time-of-day variation in the abundance of 639 (12%) proteins and 1,823 (29%) phosphosites in liver (Fig 4B), consistent with previous studies ^92,104–107^. Western blotting confirmed a daily rhythm in RPS6-S235/236 phosphorylation consistent with the mass spectrometry data (Fig S4I).

mTOR inhibition by INK128 had a profound effect on the daily composition of the hepatic proteome, closely mirroring our results in fibroblasts (Fig 1I, S1P): 86% of proteins showing significant time-of-day variation lost this variation under INK128 (Fig 4B, S4J, Table S1) indicating that mTOR signalling is required for daily variation in hepatic protein abundance. Interestingly, in the phosphoproteome, around half of phosphosites with time-of-day variation were lost upon mTOR inhibition, while an approximately equal number gained daily variation, suggesting that mTOR may normally suppress phosphorylation at roughly as many sites as it stimulates throughout the day. Critically, as with the effect of rapamycin on rhythmic transcription ^57^, time-of-day variation in the abundance and phosphorylation of clock proteins such as CRY1 and PER1 were unaffected by mTOR inhibition (Fig 4C). The profound remodelling of the daily hepatic (phospho)proteome under INK128 therefore cannot be attributed to TTFL disruption, demonstrating that many daily physiological rhythms in the liver can be uncoupled from clock gene activity.

Unlike protein abundance rhythms, daily phosphorylation peaked at the end of the rest phase and start of the active phase (ZT6-12) (Fig S4K,L), irrespective of experimental group, which is also when daily mTOR-dependent phosphorylation was highest (Fig 4D). GO analysis of these mTOR-dependent phosphosites revealed enrichment for transcriptional regulation (Fig 4E), and kinase enrichment analysis of this phase cluster confirmed differential activity of established mTORC1 effector kinases such as P70S6K and ULK1 (Fig 4F), consistent with mTOR driving daily variation in transcriptional and translational regulation during the start of the active phase. As in fibroblasts, the enrichment for RNA metabolism and processing among proteins losing time-of-day variation under INK128 suggested that daily variation in hepatic RNA metabolism requires mTOR activity. To test this *in vivo*, we perfusion-fixed mice every 6 hours across one LD cycle and stained liver sections for total RNA (Fig 4G). Consistent with our findings in fibroblasts (Fig 1F), nuclear and cytoplasmic RNA showed significant diurnal variation, which was clearly attenuated by INK128 treatment even though clock protein rhythms were not (Fig 4C).

Next, we examined the contribution of mTOR activity to daily organismal physiology. A functional clockwork is associated with proper metabolic function ^110^, and daily rhythms in metabolic gas exchange are a well-established readout of organismal circadian physiology. We assessed metabolic gas exchange using indirect calorimetry in mice pre-treated with INK128 for 3 days, followed by 4 days of continuous measurement under 12h:12h light:dark cycles. As expected ^52^, vehicle-treated mice showed clear daily rhythms in respiratory exchange ratio (RER), energy expenditure, oxygen consumption rate (VO_2_), and CO_2_ release rate (VCO_2_) (Fig 4H, S4M) that peaked during the dark phase. RER mesor (midline of the rhythm) was significantly reduced under INK128 treatment relative to control (Fig 4H), indicating a shift from carbohydrate metabolism towards protein and lipid metabolism, consistent with the role of mTOR in lipid metabolism ^111^. Critically, daily rhythms in RER were severely attenuated under mTOR inhibition, phenocopying previous observations of arrhythmic RER upon liver-specific disruption of the TTFL ^112^. Together, these data indicate that mTOR is required for the normal daily rhythm in substrate utilisation, which would otherwise track behavioural state ^113^, implicating mTOR activity as a major driver of metabolic rhythmicity.

Proteomics analysis showed mTOR to influence xenobiotic metabolism, including the phase I enzyme CYP2E1 and UDP-glucuronate synthesis regulator UGDH ^114^ (Fig 4C). Time-of-day variation in hepatic drug metabolism is of clear clinical relevance but is challenging to predict from protein or transcript abundance alone ^16,92,115,116^. To test whether daily variation in mTOR activity contributes to temporal differences in hepatic xenodetoxification, we selected two drugs metabolised by distinct hepatic cytochrome P450 enzymes: bupropion, which is metabolised to hydroxybupropion by CYP2B10 (CYP2B6 in humans) ^117^, and atorvastatin, which is metabolised to hydroxyatorvastatin by CYP3A11 (CYP3A4 in humans) ^118^. Mice pre-treated with vehicle or INK128 were injected simultaneously with bupropion and atorvastatin at ZT9 or ZT21, the timepoints at which our phosphoproteomic data predicted the greatest difference in liver function, and blood plasma and liver were harvested 30 minutes later for mass spectrometric quantification of drugs and metabolites (Fig 4I). Hydroxybupropion showed a significant time-of-day difference in abundance under vehicle conditions, which was abolished by INK128 treatment (Fig 4J), whereas hydroxyatorvastatin showed no time-of-day difference under either treatment. This proof-of-principle experiment indicates that, for drugs with diurnal pharmacokinetics, daily variation in mTOR activity may contribute to time-of-day variation in their hepatic metabolism *in vivo*.

Altogether, our observations demonstrate that short-term partial inhibition of mTOR activity attenuates daily physiological rhythms across the hepatic (phospho)proteome, RNA abundance, metabolic gas exchange, and drug metabolism. Critically, this occurred without disruption of central circadian timekeeping mechanisms or clock gene/protein activity, indicating that mTOR activity is required for most daily physiological rhythms in the liver.

### Daily forebrain physiology is uncoupled from clock protein rhythms

The contribution of mTOR to daily physiological rhythms in the brain is less well understood than in peripheral tissues such as liver. To gain new insights we extended our (phospho)proteomic analysis to forebrains harvested every 6 hours across the LD cycle from the same mice described above (Fig 4A, 5A, Table S1). INK128 adequately penetrated the blood-brain barrier: phosphorylation of S6K and AKT was reduced by ∼35% and ∼50% respectively (Fig S5A), and the number of phospho-S6-positive cells in cortical slices was reduced by ∼34% (Fig S5B). Phosphoproteomic analysis confirmed no time-of-day variation in mTOR pathway readouts under INK128 (Fig S5C). We detected 6,428 proteins and 7,703 phosphosites, of which 229 (3.5%) proteins and 2,706 (35%) phosphosites showed significant daily variation in control mice. The vast majority of these were lost upon INK128 treatment (Fig 5B, S5D-F), despite daily rhythms of locomotor activity being unaffected (Fig S4E). As a striking example, 27 of 41 detected phosphosites on the abundant microtubule-associated phosphoprotein tau (MAPT) showed daily variation that was abolished by mTOR inhibition (Fig S5G).

**Figure 5:**
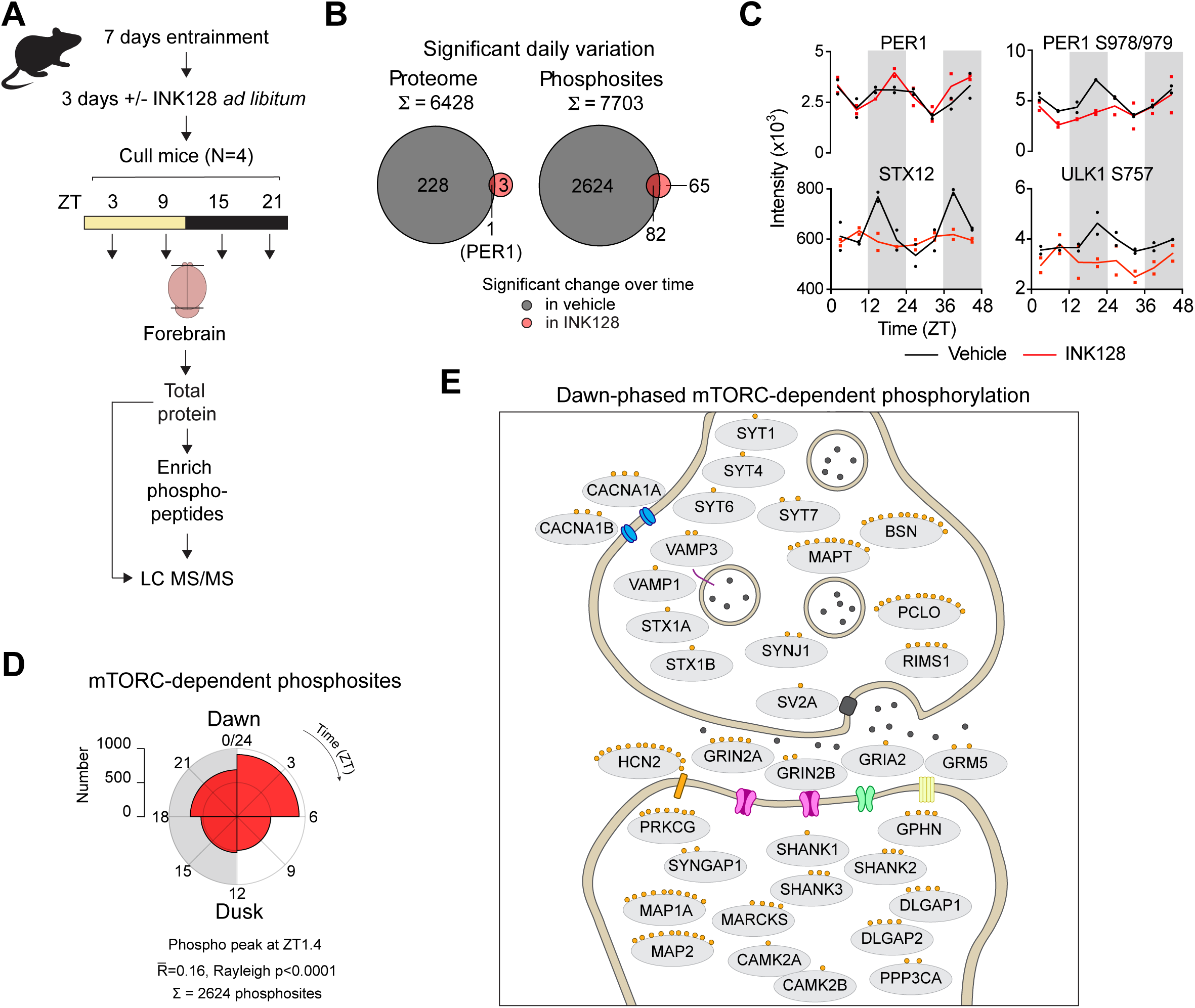
Partial mTOR inhibition severely attenuates daily physiology in the brain A. Mice were entrained under 12h:12h LD for 7 days, treated with vehicle or INK128 (25 µM) and culled every 6 hours across one LD cycle (N=4 mice per timepoint per condition). Forebrains were enriched for phosphopeptides and analysed by LC MS/MS. B. Venn diagram showing overlaps between proteins and phosphosites with significant daily variation under vehicle or INK128 treatment (BioDare2 eJTK cycle Benjamini-Hochberg FDR ≤0.05). C. Top: example PER1 protein abundance and S978/979 phosphorylation. Bottom: STX12 protein abundance and phosphorylation of mTOR target ULK1 S757. Data plotted across 2 days for visualisation purposes, but all analyses were performed treating timepoints as a single day (line= mean). D. Rose plot of peak phases of mTOR-dependent phosphosites (rhythmic under vehicle but not under INK128), binned into 6 h windows. Mean peak phase, mean resultant length (R̄) and Rayleigh test p-value are shown. E. Synaptic components carrying mTOR-dependent phosphosites around dawn (ZT18-6).

Of all detected proteins, PER1 was the only one showing time-of-day variation under both vehicle and INK128 treatment (other core clock proteins were not detected) (Fig 5B,C), demonstrating continued TTFL function throughout the forebrain. The small number (82/2706) of significantly varying mTOR-independent phosphosites included PER1 S978/979 and the circadian transcription factor DBP S86 (Fig S5H). Thus, as in liver, daily global variation in forebrain (phospho)proteome composition can be decoupled from both clock protein activity and daily locomotor rest/activity cycles.

In contrast with liver, where most mTOR-dependent phosphorylation peaked during the late day and early night (ZT6-18), mTOR-dependent phosphorylation in the forebrain peaked mostly during the late night and early day (Fig 5D, ZT18-6). GO analysis of this ZT18-6 phosphosite cluster, coinciding with the end of active/wake phase in nocturnal mice, revealed strong enrichment for components of the pre- and post-synaptic membrane (Fig 5E, Fig S5I). mTOR is a known driver of synaptic protein synthesis ^119,120^ and plasticity ^121–123^, and the circadian clock promotes arousal at the end of the active phase to counteract rising homeostatic sleep pressure ^124,125^. This is consistent with previous observations that sleep-wake cycles drive synaptic phosphorylation dynamics^106,107,126^ and suggests that mTOR outputs may contribute to clock-driven regulation of sleep-wake transitions at this time.

### mTOR inhibition affects sleep-wake architecture in mice and zebrafish

To test whether mTOR activity contributes to circadian regulation of sleep-wake transitions, we used electroencephalography (EEG) to analyse sleep-wake architecture in mice before and during INK128 treatment (Fig 6A). While the daily profiles of wakefulness, non-rapid-eye-movement (nREM, quiescent) and REM (active) sleep were largely preserved (Fig 6B), mice treated with INK128 were significantly more wakeful and had less nREM sleep around the light-dark transition (Fig 6C, S6A), consistent with locomotor activity (Fig S4H). Wakeful bouts were longer under INK128 treatment, while nREM and REM bout lengths were unchanged (Fig 6D, S6B). Power spectral density (PSD, EEG signal across frequencies) analysis showed minimal time-of-day difference in mice prior to INK128 treatment (Fig S6C-E), with no significant effect of mTOR inhibition on frontal or parietal cortex during any sleep-wake state (Fig S6C, D), and nREM delta power (1-4 Hz) was similarly unaffected (Fig 6E, S6F). Together, these findings suggest that mTOR-dependent processes shape sleep timing at a specific circadian window, without a detectable change in electrophysiological markers of sleep pressure.

**Figure 6:**
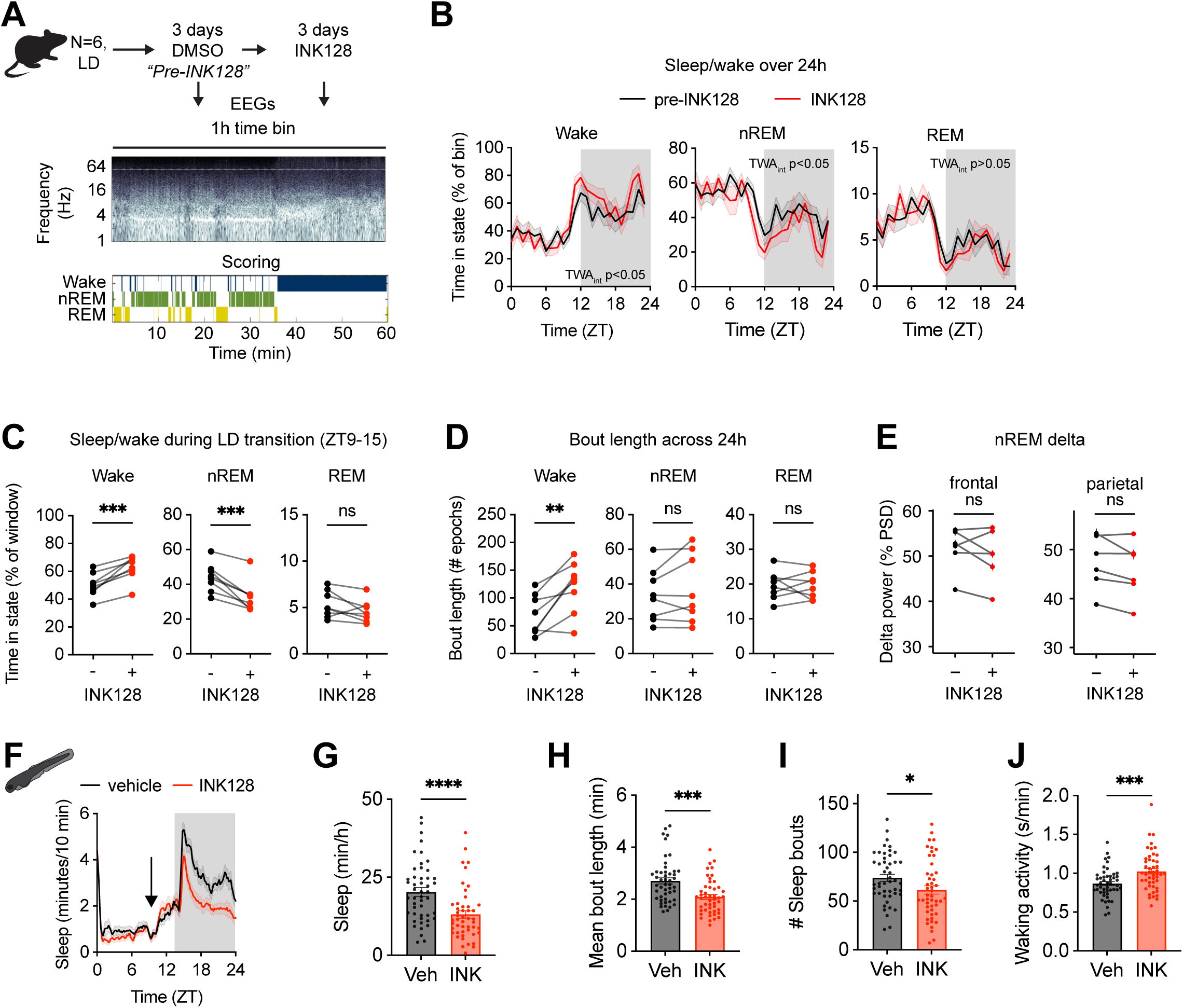
mTOR inhibition affects sleep-wake architecture in mice and zebrafish A. Experimental schematic. Mice implanted with EEG/EMG electrodes were singly housed under 12h:12h LD cycles. After surgical recovery, mice received vehicle in drinking water for 3 days, followed by INK128 (25 µM) for 3 days. EEG was recorded continuously throughout both conditions. Representative power spectrogram and corresponding sleep-wake scoring in 1h bins are shown. Quantification reflects treatment days 2-3. B. Daily profiles of wake, nREM and REM sleep expressed as a percentage of each 1h recording bin across 24h, before or after INK128 treatment (N=6, mean ± SEM, repeated measures TWA with Sidak’s MCT). C. Scored sleep-wake behaviour during the light-dark transition (ZT9-15) was compared pre- and during INK128 treatment (paired t-test). D. Bout lengths across 24h for wake, nREM and REM (paired t-test). E. nREM delta power (1–4 Hz) recorded from the frontal and parietal cortex (Wilcoxon test). F. Sleep in zebrafish larvae (defined as inactivity > 1 min duration) over 24h. Vehicle or 10 µM INK128 at ZT9-10 (black arrow). Lines represent a rolling average of mean total sleep per 10 minutes (n=48 per condition, mean ± SEM). G. Average nighttime sleep in larvae (Welch’s t-test, mean ± SEM). H. Mean sleep bout length (Welch’s t-test, mean ± SEM). I. Number of sleep bouts (Welch’s t-test, mean ± SEM). J. Waking activity (Welch’s t-test, mean ± SEM).

We could not achieve higher levels of mTOR inhibition in mouse brain without adverse effects, therefore, to explore the generality of our findings, we extended our approach to zebrafish larvae. This non-mammalian vertebrate is a well-established model for daily sleep/wake cycles ^127,128^ in which small molecules can be delivered passively and continuously by bath exposure ^129^. Moreover, like in diurnal humans, zebrafish sleep is consolidated at night, whereas sleep of the nocturnal mouse is much more fragmented.

We first applied a range of INK128 concentrations directly into their water late in the day and observed clear dose-dependent reductions in nighttime sleep and length of sleep bouts that mirrored our observations in mouse (Fig S6G-J). We then validated these findings in older larvae (7 dpf), by adding INK128 at the empirically determined lowest effective concentration (10 µM) in the middle of the afternoon (ZT9-10). Sleep throughout the following night was significantly attenuated by mTOR inhibition (Fig 6F, G), with significant reductions in both the number and length of sleep bouts and enhanced waking activity at night (Fig 6H-J).

Taken together, these data show that mTOR activity contributes to sleep duration and consolidation, in both diurnal and nocturnal vertebrates in a time-of-day dependent fashion. However, despite the profound loss of daily variation in the mouse forebrain (phospho)proteome composition under mTOR inhibition, daily sleep-wake and rest-activity cycles are mostly preserved, demonstrating that the majority of mTOR-dependent proteome and phosphoproteome rhythms in the brain are not required for the daily organisation of behaviour. Instead, we speculate that these rhythms may primarily fulfil a longer-term function to maintain neuronal protein homeostasis through temporal consolidation of proteome renewal and protein complex turnover ^22^.

### Conserved role of TOR signalling in circadian output regulation

TOR signalling is essential and conserved throughout eukaryotes ^130,131^. In the present study we have shown that mTOR is not a clock component, but functions as a major circadian output pathway regulating daily physiological rhythms. We therefore asked whether this circadian effector function is conserved across the eukaryotic lineage. To test this, we examined core timekeeping and output rhythms under TOR inhibition in two distantly related eukaryotes: *Arabidopsis thaliana* (Plantae) and *Neurospora crassa* (Fungi). Because the catalytic kinase domain of TOR is highly conserved across eukaryotes, selective ATP-competitive inhibitors such as INK128 and torin1 potently inhibit TOR activity across many species ^132,133^, whereas the absence of FKBP12 conservation in plants renders rapamycin and its derivatives ineffective ^134^.

We found that INK128 elicited similar effects on bioluminescent core clock reporters in plants (Fig 7A) and fungi (Fig 7B) as in mammalian cells (Fig S1A), specifically, CCR2::LUC (reporter of the cold-circadian rhythm-RNA binding 2 gene) in *A. thaliana*, and *pfrq_cbox_:luc* (reporter of the CLOCK-box of the *frequency* promoter) in *N. crassa*. INK128 treatment led to modest increases in period in fungi, but not plants (Fig S7A,B), whereas baseline luminescence was strongly suppressed (Fig S7C,D) consistent with the drop in global protein synthesis that is a hallmark of mTORC1 inhibition. To test the effects of TOR inhibition on output rhythms we examined rhythmicity of luciferase reporters of primary metabolic genes, glutamine-dependent asparagine synthetase 1 (DIN6) in *A. thaliana* and D12/D15 fatty acid desaturase (NCU09497) in *N. crassa* under the same conditions (Fig 7C,D). Consistent with mouse cells and tissues, output rhythms were more strongly repressed by TOR inhibition than core clock rhythms (Fig 7E). We suggest that, like casein kinase 1, the circadian function of TOR signalling may be conserved from the last eukaryotic common ancestor as a central conduit of timing information (Fig 7F) ^135^.

**Figure 7:**
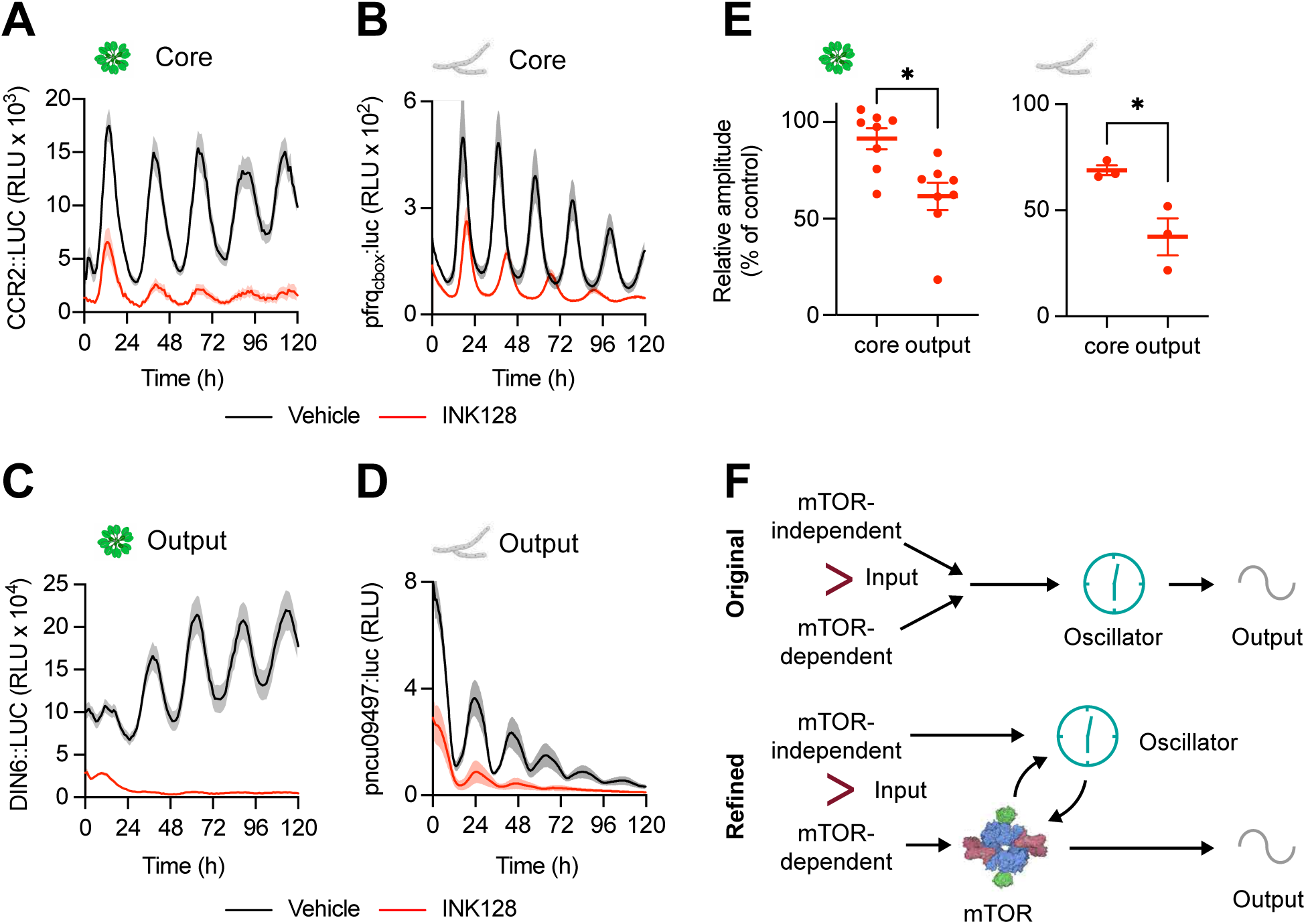
mTOR signalling is a conserved conduit of timing information A. *Arabidopsis thaliana* plants expressing a reporter of core clock activity, CCR2::LUC ^143^ , the presence of INK128 (red) or vehicle control (black) (N=6-8, mean ± SEM) B. *Neurospora crassa* expressing the core clock reporter, *pfrq_cbox_:luc* ^144^, the presence of INK128 (red) or vehicle control (black) (N=3, mean ± SEM). C. *Arabidopsis thaliana* plants expressing a bioluminescent reporter representing clock output, DIN6::LUC ^145,146^, the presence of INK128 (red) or vehicle control (black) (N=6-8, mean ± SEM). D. *Neurospora crassa* expressing a bioluminescent reporter of clock output, P*ncu09497::luc* ^147^ in the presence of INK128 (red) or vehicle control (black) (N=3, mean ± SEM). E. Relative amplitude of core and output rhythms in the presence of INK128, expressed as a percentage of control. Relative amplitude was calculated as the amplitude of a sine wave fit (24-120h) divided by the mean luminescence over the same period and compared between core and output by t-test. F. Cartoon representations of the flow of timing information in circadian systems. Under the original model, timing information flows vectorially from input (daily timing cues) through the oscillator to outputs, such as proteome renewal in cells and locomotor activity in animals. Under our refined, mTOR-centric model, mTOR activity sits between the oscillator and rhythmic outputs, dictating the timing of most, but not all, output rhythms across Kingdoms.

## Discussion

The dominant model of circadian physiology makes a prediction that can be tested directly: if daily rhythms in physiology are driven by clock-protein-dependent transcription of clock-controlled genes, those rhythms should not be separable from clock protein rhythms. Here they were. Acute mTOR inhibition abolished the majority of daily variation in the proteome, phosphoproteome, RNA content and metabolic output of cultured cells, liver and forebrain, while clock protein rhythms, SCN-dependent behavioural timing and sleep–wake organisation persisted. Daily physiology is therefore not contingent upon clock protein rhythms in the manner the model requires, and an alternative account of what imposes daily organisation on physiology is needed.

That this assumption has persisted is less a reflection of the evidence for it than of the absence, until now, of a mechanistically plausible alternative. For example, rather than acutely synchronising cells, most daily systemic cues gradually entrain cellular clocks over many days^11,47,136^. We therefore propose a simple alternative model: that most daily rhythms arise post-translationally as direct and/or indirect consequences of daily rhythms of rhythmic AKT/mTOR signalling, which itself arises through an interaction between cell-intrinsic and extrinsic timing cues. Extrinsic timing cues include feeding-fasting cycles via insulin signalling that directly drive mTOR rhythms, while our data reveal a previously unrecognised intrinsic component arising from direct association of PER2 with mTORC1, such that daily accumulation of PER2 during the rest phase attenuates mTORC1 activity.

As a hub sensing and integrating metabolic, growth and environmental signals, mTORC1/2 ultimately regulates almost every cellular process including transcription, translation, proteostasis and enzyme activity ^38^. This places it ideally to couple temporal information to cellular function across tissues and species. We show that mTOR signalling is required for rhythmic regulation of diverse physiological processes both cell-autonomously and in tissues *in vivo*. Pharmacological inhibition of the mTOR pathway abolished the majority of cell-autonomous circadian rhythms in protein abundance and phosphorylation, without disrupting circadian timekeeping itself, consistent with rapamycin’s reported effects on rhythmic gene expression in the liver ^57^. Indeed, even the partial mTOR inhibition achieved *in vivo,* carried out under physiological conditions where daily timing cues are still present, had similar effects in mouse forebrain and liver to those observed in isolated cells, and had a greater effect on the rhythmic proteome and phosphoproteome than reported for most clock gene mutants^18,21,92^.

The similarity of this function across distantly related eukaryotes, with TOR inhibition suppressing output rhythms more strongly than core oscillator rhythms in both *Arabidopsis* and *Neurospora*, further supports the view that mTOR acts as a deeply conserved conduit between the circadian oscillator and its physiological outputs. Global mTOR inhibition or activation could plausibly act independently of the circadian oscillator; however, the direct physical interaction between PER2 and mTORC1, the antiphase activity relationship, and elevated hepatic mTORC1 activity in *Per1/2-*deficient mice point to a specific regulatory link between cellular clock proteins and mTOR signalling.

Daily systemic cues have long been thought to act synergistically with cell-intrinsic circadian timekeeping to facilitate healthy physiology. Whereas, outside the SCN and so long as there is a daily feeding rhythm, rhythms in mTOR activity and many other physiological rhythms do not require a functional TTFL ^11,20,57,137^. In other words, systemic regulation by daily cycles of feed-fast (via insulin pathway), light-dark (via glucocorticoid signalling), and rest-activity (via temperature) appear dominant over cell-intrinsic circadian mechanisms. We speculate that the primary function of the circadian clock in peripheral tissues is homeostatic - to buffer against mistimed extrinsic signals and oppose exaggerated responses, rather than to drive "anticipatory rhythms” ^110^. Extending this logic, rather than being "essential to the daily organisation of biological processes" ^57^, we propose the testable hypothesis that many transcriptional rhythms are a consequence, rather than a cause, of circadian homeostatic regulation - a possibility that could be addressed by direct comparison of transcriptional and post-translational rhythms under mTOR inhibition.

By contrast, locomotor and sleep-wake organisation were only modestly affected by mTOR inhibition, consistent with the evidence that hypothalamic SCN-dependent behavioural timing is remarkably robust to inhibition of mTOR and other major signalling pathways ^11,47,138–140^. Given this modest effect on sleep-wake state, sleep-driven mechanisms are unlikely to explain most of the rhythmic changes we observe, suggesting a circadian rather than sleep-dependent origin. However, mTOR inhibition did produce a specific increase in wakefulness in both mice and zebrafish, at specific times of day, suggesting that mTOR outputs contribute to circadian regulation of sleep-wake transitions, without being required for overall daily organisation of behaviour. Indeed, in patients with Smith-Kingsmore syndrome caused by hyperactivating *MTOR* mutations, low-dose rapamycin treatment improved sleep-wake disturbances ^60^, suggesting that mTOR activity must remain within a defined range to support normal sleep-wake timing ^60,107^.

Together, these findings suggest that mTOR inhibition dissociates circadian timekeeping from most rhythmic daily physiological outputs. This does not imply that all rhythms depend on mTOR, as some persisted during inhibition and robust behavioural timing was retained, indicating contributions from additional pathways. Nonetheless, our data support a model in which mTOR signalling acts as a conduit through which intrinsic timekeeping and environmental signals impose daily organisation on physiology, a function that appears conserved across eukaryotes.

### Limitations of the study

We chose a pharmacological approach to investigate the effect of reducing the activity of the mTOR pathway because complete deletion of mTOR is lethal. While INK128 shows high potency and selectivity for mTOR over other kinases ^63^, as with any pharmacological approach, off-target effects remain possible, particularly at higher concentrations and with prolonged exposure. In addition, the use of INK128 does not distinguish between the relative contributions of mTORC1 and mTORC2, and we have not investigated specific effectors downstream of either pathway. Moreover, as noted in the introduction, our experiments establish that mTOR activity is necessary for rhythmic physiological outputs, but do not establish that rhythmic mTOR activity itself is required to sustain these downstream rhythms. We note that INK128 suppresses global protein synthesis and phosphorylation; this may reduce absolute signal levels to the point where residual rhythmicity, if present, falls below detection limits, but we have endeavoured to rule out this possibility. Some observed effects may therefore reflect secondary consequences of sustained pathway inhibition or suppression of protein synthesis, rather than direct suppression of mTOR signalling itself. We did not test inhibition of other broadly acting anabolic/signalling hubs (e.g. AMPK, GSK3).

Our *in vivo* experiments were performed in whole tissues where complete inhibition was not achieved: mTOR activity was reduced to 25-50% of control. This represents the maximum tolerated level of inhibition before adverse effects were observed (e.g., weight loss) but also reduces the magnitude of observed effects compared with cultured cells. In addition, as dosing was systemic, we cannot exclude off-target contributions, although we did confirm pathway engagement by S6K and AKT phosphorylation. Furthermore, our sampling interval for proteomics studies represents a compromise that inevitably limits the identification of rhythms with lower amplitude or higher variance ^17,141^. Furthermore, this sampling frequency may be too low to resolve subtle shifts in phase or waveform. Changes in rhythmicity could arise from altered phase coherence between cells as well as loss of rhythms within individual cells but, with the exception of the microscopy studies (Fig 1F, 4G), our measurements were taken at the population level (i.e., well of cells, whole tissue) and therefore cannot differentiate between these two possibilities. Therefore, although these data demonstrate that mTOR activity is necessary for many rhythmic outputs, further optimization of techniques and manipulations will be important to refine the causal architecture we have proposed at the cellular level. Future work in this direction would be desirable across many different cell/tissue types, to establish the generality of mTOR signalling as an important input to, and a major output from, the cellular circadian clock.

Our data support PER2 as a negative regulator of mTORC1 but do not fully establish the mode-of-action. PER2 physically associates with mTORC1 components and acute PER2 depletion increases phosphorylation of mTORC1 substrates, yet we have not demonstrated direct inhibition of mTOR kinase activity by PER2 in a reconstituted *in vitro* system and cannot exclude indirect contributions. Moreover, although PER2 abundance and mTORC1 activity are antiphasic in cultured cells, PER2 was acutely depleted at a single time point, so, although plausible and consistent with the literature, we have not directly demonstrated that PER2-mediated inhibition of mTORC1 varies with circadian phase. Further work, including in vitro kinase assays with purified components and PER2 depletion at multiple circadian times, will be important to test whether PER2 inhibits mTORC1 directly and in a time-dependent manner.

## RESOURCE AVAILABILITY

### Lead contact

Further information and requests for resources and reagents should be directed to and will be fulfilled by the lead contact, John O’Neill.

### Materials availability

Plasmids generated in this study are available upon request.

### Data and code availability

• Proteomics data have been deposited to the ProteomeXchange Consortium (http://proteomecentral.proteomexchange.org) via the PRIDE partner repository with dataset identifier PXD085200, and are publicly available as of the date of publication. All accession numbers are listed in the key resources table. Microscopy data reported in this paper will be shared by the lead contact upon request.

• Original code for analysing proteomic data has been deposited on Zenodo, DOI: 10.5281/zenodo.23105933 and DOI: https://doi.org/10.5281/zenodo.23046941

• Any additional information required to reanalyse the data reported in this paper is available from the lead contact upon request.

## Acknowledgements

JSO is supported by the Medical Research Council (MC_UP_1201/4) as part of United Kingdom Research and Innovation, and AstraZeneca BlueSky 2.0. AZ and RSE are supported by UKRI Future Leaders Fellowship MR/Y017552/1. AND and LLBD are funded by UKRI-BBSRC (Institute Strategic Programme BRiC BB/X01102X/1) and the European Union (ERC, SyG MicroClock, 101166968). AG and LFL are funded by ANID-Millennium Science Initiative Program-Millennium Institute for Integrative Biology (iBio ICN17_022), ANID/FONDECYT 1251234. We thank O’Neill lab members past and present for useful discussions and valuable feedback; Dr. Olga Perišić for assistance with the Expi293 cell culture and plasmid expression and for useful discussions about the PER protein purification; Dr. Iain Hay for providing the purified human mTORC2 complex; Dr. Roger Williams for insightful discussions about mTOR biology; Dr. Mick Hastings for providing the PDKO mice; MRC LMB animal facility staff for assistance with experiments, breeding, and animal care; MRC LMB Light Microscopy facility; Antonio di Scoccio for mouse sleep analysis; Dr. David Wong for developing the initial proteomics analysis pipelines.

## Author contributions

Conceptualization-JSO

Investigation-AZ, AM, MA, AG, LLBD, SPC, GP, ADB, EAH, LCA, TS, TEM, CB, JF, AS, NRJ, JM, ES, JR

Formal analysis-AZ, AM, AaR, SPC, TS, TEM, JSO, EAH, RRM, JR, LCA, ADB, GMR, CL

Writing – original draft-AZ, AM, ADB, JSO

Writing – reviewing and editing-AZ, JSO, AM, ADB, SPC, TS, MA, AaR, LLBD, NRJ, RSE, ED, DAB, LFL, AND, JR, GMR, JW, PN, CL, CH

Resources-JSO, LJH, CL

Funding acquisition-JSO, RSE

Supervision-JSO, CL, RSE

## Declaration of interests

JW, CH and LCA are employees of and hold shares in AstraZeneca. PN holds stock in AstraZeneca.

## Declaration of generative AI and AI-assisted technologies in the manuscript preparation process

During the preparation of this work, the authors used Claude (Anthropic) to improve the readability of the text and to assist with debugging and refining proteomics analysis code. The authors reviewed and edited the output as needed and take full responsibility for the content of the published article.

---

Figure S1: mTOR inhibition disrupts circadian regulation of multiple cellular physiological outputs, related to Figure 1

A. Representative bioluminescence trace of PER2::LUC fibroblasts treated with 1 µM INK128 36 hours after release into constant conditions (T36) (n=6, mean ± SEM)

B. Phase response curve of PER2::LUC fibroblasts treated with INK128 every 4 hours across 1 circadian cycle (n=6, mean ± SEM, null hypothesis: gradient = 0). Phase shifts were calculated relative to vehicle treatment and plotted normalised to period as circadian time (CT).

C. Period of INK128-treated cells compared to control vehicle at T24 (two-tailed t-test, mean ± SEM).

D. Amplitude of INK128-treated cells compared to control vehicle at T24 (two-tailed t-test, mean ± SEM).

E. Baseline PER2::LUC bioluminescence at T24 of INK128-treated cells compared to control vehicle (Veh) (two-tailed t-test, mean ± SEM)

F. Western blot of S6K phosphorylation at T389 in response to vehicle or INK128 (0.01, 0.1, or 1 µM, 2h).

G. Relating to Fig 1B. Quantification of total protein abundance in fibroblasts treated with INK128 (1 µM, starting at T0) and harvested every 4 hours for 3 days (n=3, mean ± SEM, p-value refers to preferential fit to a straight line or damped cosine wave, null hypothesis = no rhythm).

H. Relating to Fig 1B. Phase of PER2::LUC bioluminescent reporter (n=3, mean ± SEM, unpaired t-test).

I. Relating to Fig 1D. Right: puromycin (Puro) is incorporated into nascently translated peptides with a proportion degraded by the proteasome. Left: bortezomib (BTZ) inhibits the proteasome, preventing this degradation such that all nascently translated peptides can be detected. Degradation is inferred by calculating the proportion of Puro vs Puro with BTZ (100 x (1 - Puro/ (Puro with BTZ))). Bottom: Representative western blot of puromycin assay without or with bortezomib, with enhanced contrast below, and total protein stained with Ponceau S.

J. Relating to Fig 1D: quantification of puromycin incorporation without bortezomib (n=6, mean ± SEM, preferential fit to a straight line or damped cosine wave, null hypothesis = no rhythm).

K. Relating to Fig 1F. Quantification of circadian rhythms in nuclear RNA abundance in fibroblasts fixed and stained with SYTO RNASelect Green and DAPI every 4 hours for 2 days (n=3 wells with >30 cells per field of view, TWA with Sidak’s MCT, mean ± SEM), with PER2::LUC bioluminescence below (n=3, mean ± SEM).

L. Relating to Fig S1K. Representative images (scale bar 25 µm).

M. Relating to Fig 1G. Representative images of CellTracker Red-labelled fibroblasts immediately after scratch wounding (T0) and following 60 hours healing, for wounds made either 24 or 36 hours after treatment with vehicle or INK128 (scale bar 25 µm).

N. Relating to Fig 1H: PER2::LUC bioluminescence traces for each biological replicate cell line treated with vehicle or INK128 (n=4, mean ± SEM).

O. Relating to Fig 1H. Phosphosite abundances of AKT1 S473 and RPS6 S244 (paired t-test).

P. Relating to Fig 1H. Distributions of fold change over time in proteins and phosphosites showing significant time-of-day variation only under vehicle, with corresponding fold changes under INK128 treatment (n= number of proteins/phosphosites, one-sided Wilcoxon rank sum test).

Q. Relating to Fig 1H. Ranked gene ontology (GO) enrichment analysis for molecular function performed using GOrilla with (phospho)proteins that varied significantly over time under vehicle but not INK128 treatment, compared to a background of all detected proteins. Redundant terms were simplified using REViGO. Bars represent number of genes associated with each term, coloured by FDR q-value and labelled with fold enrichment.

Figure S2: The effect of AKT-mTOR pathway activation and inhibition on PER2::LUC rhythms in fibroblasts, related to Figure 2

A. Left: Western blot of S6K phosphorylation at T89 in response to vehicle or rapamycin (0.1 or 1 µM, 2h), controls for S6K phosphorylation were 16h serum starved versus 2h 50% serum shock. Middle: blot of AKT phosphorylation at S473 in response to 2-hour treatment with 1 µM INK128 (INK), 1 µM MHY1485 (MH), 1 µM rapamycin (rap), 1 µM miransertib (mir), and 16h serum starve vs 2h 50% serum shock. Right: western blot of p44/42 MAPK phosphorylation at T202/Y204 in response to vehicle or U0126 (0.1, 1, or 10 µM, 2h).

B. Representative trace of fibroblasts pre-treated with INK128 followed by a +100 mOsm hyperosmotic shock (100 mM sorbitol, n=4, mean ± SEM, arrow indicates time of treatment).

C. Left: circadian phase for Fig S2B plotted on a circular axis corresponding to a 24-hour day (mean ± SEM, Watson-Williams test between indicated groups). Right: phase shift relative to control (mean ± SEM, TWA with Sidak’s MCT).

D. Period for Fig S2B (mean ± SEM, OWA with Sidak’s MCT).

E. Representative trace of cells treated with vehicle or INK128 and treatment with dexamethasone (100 nM), forskolin (10 µM), or 10% serum (n=6, mean ± SEM, arrow indicates time of treatment). Dashed line at the peak of vehicle or INK128 control plotted for illustration purposes.

F. Top: circadian phase for Fig S2E plotted on a circular axis corresponding to a 24-hour day (mean ± SEM, Watson-Williams test between indicated groups). Bottom: phase shift relative to control (mean ± SEM, TWA with Sidak’s MCT).

G. Amplitude for Fig S2E expressed as a percentage of pre-treatment control (mean ± SEM, TWA with Sidak’s MCT).

Figure S3: PER2 modulates the activity of mTORC1, related to Figure 3

A. Relating to Fig 3B. Selected TTFL components, mTOR complex components, and mTOR regulatory proteins enriched in endogenous PER2-HaloTag immunoprecipitations relative to HaloTag controls, identified by LC–MS/MS.

B. Relating to Figure 3D. In vitro Strep-tag pulldown assays assessing the interaction between purified human mTORC2 with purified human, full-length PER2. Input and eluate fractions were analysed by Coomassie-stained SDS–PAGE. The "PER2 + beads" lane contains PER2 incubated with Strep-Tactin resin in the absence of bait, processed in parallel with equal PER2 input and an equal number of washes, and controls for non-specific binding of PER2 to the resin.

C. Relating to Fig 3F. Inter-protein crosslinked peptides identified by XL–MS between PER2 and mTORC1 subunits.

D. Relating to Figure 3I. Detailed quantification of all detected AKT1S1 phospho-peptides following acute PER2 depletion in U-2 OS cells expressing endogenous PER2-HaloTag induced by HaloPROTAC3 treatment relative to DMSO control. 29 out of the 136 *in vitro* validated mTORC1 substrate sites were detected in this experiment. Please note that, as a group, detected mTORC1 substrate sites were not more likely to be differentially phosphorylated than other sites (10.3% vs 5.6% Fisher’s exact test p = 0.22) presumably because steady state phosphorylation levels at other sites either adapt to 24h PER2 depletion (e.g., by reduced dephosphorylation) or because not all mTORC1-regulated sites are equally sensitive to PER2 depletion.

Figure S4: Partial mTOR inhibition *in vivo* uncouples daily physiology from behaviour, related to Figure 4

A. Western blots and quantification of liver from mice treated with vehicle, 8 or 25 µM INK128 provided *ad libitum* in drinking water for 3 days, culled at ZT0, and probed for phosphorylated and total RPS6 and p-AKT (N=4, OWA with Dunnett’s MCT, mean ± SEM).

B. Change in mouse weights before and after 14 days INK128 treatment (OWA with Dunnett’s MCT, mean ± SEM, N=4).

C. Western blot of phosphorylated 4E-BP1 in livers of mice intraperitoneally injected with 1 mg/kg or 3 mg/kg INK128 and livers harvested after the indicated timepoints (OWA with Dunnett’s MCT, mean ± SEM, N=4). The functional half-life we observed for INK128 (103 mins) after injection of an acute bolus was consistent with the measured half-life 1.5-2 hours. Based on literature observations ^63^, INK128 provided ad libitum at 25 µM in drinking water is expected to achieve steady state serum concentrations peaking around 330 ng/ml at the end of the active phase, falling to no less than ∼30 ng/ml (∼100 nM) 12 hours later, assuming that water consumption occurs only in the active phase. Whilst the extent of mTOR inhibition may therefore vary over 24h, the intracellular concentration of INK128 is not expected to ever be completely saturating or to drop below that required for inhibition of mTOR activity.

D. 4E-BP1 phosphorylation at T45 from liver phosphoproteomics analysis (mean ± SEM, eJTK BH-corrected p-values shown).

E. Representative actograms showing locomotor activity for mice across 4 phases: experimental mice (right) received vehicle under LD, INK128 (25 µM) under constant darkness (DD), vehicle under DD, then returned to INK128 under DD (N=4). Control cohort (left) received vehicle throughout, the other received INK128 during treatment phases (right, red dashed box). Grey shading indicates lights off, time shown in circadian time (CT), aligned to onset of motor activity.

F. Period in DD for Fig S4E (mean ± SEM, unpaired t-test).

G. Total summed counts in DD for Fig S4E (mean ± SEM, unpaired t-test)

H. Average activity profiles of mice across 7 days LD or DD in Fig S4E (mean ± SEM, TWA with Sidak’s MCT, TWA interaction and treatment (TWA_trt_) indicated on figure).

I. Relating to Fig 4A. Western blot confirmation of liver lysates probed for total RPS6 and RPS6 phosphorylation at S235/236, with quantification of RPS6 phosphorylation by western blot compared with values measured by mass spec (TWA with Sidak’s MCT, mean ± SEM).

J. Relating to Fig 4A. Distributions of fold change over time in proteins and phosphosites showing significant time-of-day variation only under vehicle, with corresponding fold changes under INK128 treatment (n= number of proteins/phosphosites, one-sided Wilcoxon rank sum test).

K. Relating to Fig 4D. Rose plots of liver phase distribution of cycling proteins/phosphosites under vehicle or INK128 treatment, coloured by the mean relative amplitude of each phase.

L. Relating to Fig 4A. Heat maps of z-score normalised protein/phosphosite abundances. Each row represents a single protein or phosphosite, each column a biological replicate grouped by peak timepoint. Z-scores were calculated per protein/phosphosite across all replicates and timepoints. Proteins/phosphosites grouped by significant variation in either vehicle only, INK128 only, or in both conditions. Gain= gain of rhythm, Un= unaltered.

M. Relating to Fig 4H. Energy expenditure, oxygen consumption and carbon dioxide production in mice treated with vehicle or INK128 (25 µM) (N=4, mean ± SEM). Middle left: average of 3 days (mean ± SEM). Middle right: mesor for each measure (t-test, N=4, mean ± SEM). Right: amplitude for each measure (t-test, N=4, mean ± SEM).

Figure S5: Partial mTOR inhibition severely attenuates daily variation in brain (phospho)proteome, related to Figure 5

A. Western blots and quantification of forebrain lysates from mice treated with vehicle, 8, or 25 µM INK128 *ad libitum* in drinking water for 3 days, culled at ZT0, and probed for p-S6K and p-AKT (N=4, OWA with Dunnett’s MCT, mean ± SEM).

B. Representative confocal image and quantification of phospho-S6 immunostaining in visual cortex from mice culled at ZT6 (N=4, unpaired t-test, mean ± SEM, scale bar: 100 µm).

C. Relating to Fig 5A. Phosphorylation of mTORC1 substrate USP20 at S132/S134 ^142^ from forebrain phosphoproteomics analysis (mean ± SEM, eJTK BH-corrected p-values shown).

D. Relating to Fig 5A. Heat maps of row-wise z-score normalised proteins and phosphosites showing significant time-of-day variation, ordered by phase. G= gained time-of-day variation under INK128, U= unaltered.

E. Relating to Fig 5D. Rose plots showing phase distribution of significantly varying proteins (top) and phosphosites (bottom) coloured by mean relative amplitude.

F. Relating to Fig 5A. Distributions of fold change over time in proteins and phosphosites showing significant time-of-day variation only under vehicle, with corresponding fold changes under INK128 treatment (n= number of proteins/phosphosites, one-sided Wilcoxon rank sum test).

G. Relating to Fig 5A. Proportion of MAPT phosphosites showing significant time-of day variation in vehicle only (grey), both conditions (red), or neither condition (blue).

H. Relating to Fig 5A. Example brain phosphosites (mean). Data plotted across 2 days for visualisation but for all analyses, timepoints were treated as 1 day (line = mean).

I. Relating to Fig 5A. Gene ontology (GO) enrichment analysis for molecular function performed using GOrilla with phosphorylated proteins showing significant time-of-day variation under vehicle but not INK128 treatment, compared to a background of all detected proteins. Terms were simplified using REViGO. Bars represent number of genes, coloured by FDR q-value, and labelled with fold enrichment.

Figure S6: Sleep-wake organisation in mice and zebrafish under mTOR inhibition, related to Figure 6

A. Sleep/wake homeostasis per 6-hour time windows for mice (N=6, mean ± SEM, repeated measures TWA with Sidak’s MCT).

B. Wake bout length per 6-hour time windows (N=6, mean ± SEM, RM TWA with Sidak’s MCT).

C. Mean PSD across 24 hours in frontal and parietal cortex (mean ± SEM).

D. Mean PSDs per 6-hour time windows (mean ± SEM).

E. Power density in the delta domain (1-4 Hz) during nREM in the indicated time windows, recorded from the frontal cortex (N=6, Wilcoxon test, mean ± SEM)

F. Power density in the delta domain (1–4 Hz) during wake, nREM, and REM, and nREM in the presence of INK128, recorded from the frontal and parietal cortex (N=6, mean ± SEM, RM OWA + Sidak’s MCT).

G. INK128 dose-sleep response in zebrafish (N=24-48, Welch’s t-test, mean ± SEM).

H. INK128 dose-bout length response (Welch’s t-test, mean ± SEM).

I. INK128 dose-number of bouts response (Welch’s t-test, mean ± SEM).

J. INK128 dose-waking activity response (Welch’s t-test, mean ± SEM).

Figure S7: Parameters of core oscillations under mTOR inhibition in plants and fungi, related to Figure 7

A. Period of *CCR2:luc* rhythms in INK128-treated plants compared to control vehicle (Veh) (t-test, mean ± SEM).

B. Period of *pfrq_cbox_:luc* rhythms in INK128-treated fungi compared to control vehicle (Veh) (t-test, mean ± SEM).

C. Baseline luminescence of *CCR2:luc* rhythms in INK128-treated plants compared to control vehicle (Veh) (t-test, mean ± SEM).

D. Baseline luminescence of *pfrq_cbox_:luc* rhythms in INK128-treated fungi compared to control vehicle (Veh) (t-test, mean ± SEM).

## EXPERIMENTAL MODEL AND SUBJECT DETAILS

### Mice

#### Mice used for cell culture, proteomics, and behavioural experiments

C57BL/6 PER2::LUCIFERASE (PER2::LUC) mice^61^ were originally supplied by Joe Takahashi (University of Texas Southwestern) and bred at the MRC Laboratory of Molecular Biology in a specific pathogen-free barrier facility. Wild-type C57BL/6J mice originated from Jackson Laboratory. *Per1^-/-^/Per2^-/-^*mice were kind gifts from Dr. Michael Hastings ^148^. For husbandry and non-experimental housing, mice were group housed with environmental enrichment under 12h:12h light:dark cycles with lights on at 7am. All animal work was licensed by the Home Office under the Animals (Scientific Procedures) Act 1986, with Local Ethical Review by the Medical Research Council and the University of Cambridge, UK. Animal numbers were minimised in line with the 3Rs (Replacement, Reduction, Refinement) principles, with sample sizes determined by power calculation to minimise number of animals required. The animals used in this study were of both sexes (males only for behaviour and proteomics; males and females for metabolite analysis and experiment involving *Per1^-/-^/Per2^-/-^* mice) and within an age range of 2-4 months. This study is reported in accordance with ARRIVE 2.0 guidelines.

#### Mice used for sleep tracking experiments

Mice were housed in filtered cages in a temperature-controlled room with a 12:12h dark/light cycle with lights on at 6am. All procedures conformed to the EU Directive 2010/63/EU for animal experiments and the ARRIVE guidelines. Experimental protocols were approved by the Italian Ministry of Health. Experiments were performed on female and male wild-type mice (C57BL/6J, The Jackson Laboratory, Stock No: 000664), at the age of about 8 weeks.

### Zebrafish

Adult TLxAB zebrafish (*Danio rerio*) were housed in the University College London Fish Facility under the Animals (Scientific Procedures) Act 1986 project licence PA8D4D0E5 awarded to JR. Larvae were collected by natural spawning.

### Arabidopsis

Seeds of *Arabidopsis thaliana* were surface sterilised by exposure to 70% (v/v) ethanol for 1 min (VWR Chemicals, UK), followed by 10 min incubation at room temperature in 20% (v/v) sodium hypochlorite (Merck Life Science Ltd, UK). Seeds were subsequently washed two times using sterile deionised water. Seeds were resuspended in 0.1 % (w/v) water agar solution, and sown onto half-strength Murashige & Skoog plant growth media (Duchefa, Netherlands), pH = 5.7 with 0.8% w/v agar (Bactoagar, BD), in 6-well CytoOne Plate (Non-Treated; Starlab, Milton Keynes, UK). On the agar surface of each well, we positioned 4 non-phthalate PVC tubing sections (Thermo Scientific Nalgene Metric Non-Phthalate PVC Tubing), measuring 1 cm high. Between 14 – 18 seeds were sown into the ring formed by each tubing section, to produce a cluster of seedlings. Plated seeds were stratified at 4°C for 2 – 3 days in darkness and transferred to plant growth cabinets (MLR-352, PHCbi) at a continuous temperature of 19°C and cycles of 12h light / 12h darkness using photon irradiance of 80 – 100 μmol m^-2^ s^-1^, delivered from a mixture of cool and warm LED CorePro LEDtube EM/Mains (Philips, UK). Seedlings were cultivated under these conditions for 12 – 14 days. *CCR2:luc*^149,150^; *DIN6:luc* ^145,146^ lines were used for experiments.

### Neurospora

*Neurospora crassa* strains were maintained at 25°C under constant light (LL) on slants containing 1× Vogel’s minimal medium supplemented with 2% (w/v) sucrose and 1.5% (w/v) agar. Core clock oscillations were monitored using strain x654-14a (*his-3::frq c-box-luc; a*), whereas desaturase promoter activity was monitored using the *Pncu09497::luc* reporter strain (*his-3::Pncu09497-luc; a*) ^147^

### Cell lines

Mouse fibroblasts were obtained from lung tissue of adult PER2::LUC mice and immortalised by serial passage, as previously described^11^. Mice were euthanised by cervical dislocation and confirmed by exsanguination. Lung tissue was taken, stored in PBS on ice, then cut into ∼1 mm sections using sterile scalpels, and incubated at 37°C with 10 ml digestion medium for 30 minutes (DMEM/F12 (Gibco) supplemented with Pen/Strep, Mycozap Plus PR (Lonza), 0.14 U/ml Liberase (Merck)). The tissue fragments were triturated and 40 ml initial culture medium added (DMEM/F12 supplemented with Pen/Strep, Mycozap Plus PR, 15% Hyclone FetalClone III (Cytiva), centrifuged at 700 x g for 5 min, supernatant discarded, resuspended in 20 ml initial culture medium, transferred to a 10 cm tissue culture dish, and incubated at 37°C, 5% CO_2_, 3% O_2_. After 7 days, media was refreshed. After a further 7 days, cells were split and replated in selection medium (MEM (Sigma) supplemented with Pen/Strep, non-essential amino acids (Gibco), sodium pyruvate (Gibco), 10% HyClone FetalClone III). After a further 2 weeks, cells were transferred to standard culture medium (high-glucose (27.8 mM), GlutaMax-containing DMEM (Gibco) supplemented with 10% HyClone FetalClone III and Pen/Strep). Immortalisation was achieved by serial passage of cells at 20% O_2_. Cell lines were authenticated by observation of morphology and by continued expression of the bioluminescent reporter. Fibroblasts were not used after 30 passages. Cells were maintained in standard cell culture medium. Cell lines tested negative for mycoplasma contamination.

PER2-Halotag U-2 OS cells were generated by CRISPR-Cas9 mediated genome editing as previously described^47^. Cells were maintained in standard culture medium (high-glucose (27.8 mM), GlutaMax-containing DMEM (Gibco) supplemented with 10% HyClone FetalClone II and Pen/Strep) unless otherwise stated.

Expi293F cells were grown in Expi293 media (ThermoFisher) in a Multitron Pro shaker set at 37 °C, 8% CO_2_ and 125 RPM.

## METHOD DETAILS

### Cell culture experimental structure

Immortalised PER2::LUCIFERASE mouse lung cells were seeded at a density of around 30,000 cells/cm^2^ and PER2-Halotag U-2 OS cells at 62,500 cells/cm^2^ and grown to confluence in temperature cycles (12h:12h 32°C:37°C) in standard cell culture medium for a minimum of 4 days. At the transition to 32°C (mouse) or 37°C (human), cells were given a medium change and moved into constant 37°C at experimental time 0 (T0) for minimum 24 hours before any experimental perturbations.

For bioluminescent recordings, cells were given 1 mM (mouse fibroblast) or 0.3 mM (human U-2 OS) D-luciferin (Biosynth), sealed with a gas-permeable plate seal (4titude), and bioluminescence recorded in an ALLIGATOR (Cairn Research) maintained at 37°C at 30-minute intervals. Small molecules were given at the indicated times in figure legends. For all experiments, a parallel bioluminescence plate was recorded using identical experimental conditions.

### Cell treatments

Small molecules were, unless otherwise stated, used at the following final concentrations in cell culture: MHY1485-1 µM, INK128-1 µM, miransertib-1 µM, rapamycin-1 µM, murine FGF2-20 ng/ml, U0126 – 10 µM, forskolin – 10 µM, dexamethasone – 100 nM, bortezomib – 1 µM. FGF2 was dissolved in PBS with 0.1% BSA; all others were dissolved in DMSO. Unless stated otherwise, INK128 was added to cell culture medium at time=0 (T0, at the transition from 37°C to 32°C) and left in the medium throughout the experiment. Hyperosmotic medium was achieved through adding a 10x bolus of 1M sorbitol (to a final concentration of 100 mM) dissolved in serum-free cell culture medium. All perturbations were carried out on isothermal pads to minimise temperature fluctuations.

To induce endogenous PER2 protein depletion, PER2-Halotag knock-in U-2 OS cells were treated with 1 µM HaloPROTAC3 (Promega) for a minimum of 24h ^47^ before harvest or experimental perturbations.

### Protein extraction and quantification

For cellular soluble protein extraction, cells were washed twice with ice-cold PBS and incubated with digitonin lysis buffer (50 mM tris pH 7.4, 0.01% digitonin (Invitrogen), 5 mM EDTA, 150 mM NaCl, protease and phosphatase inhibitors (Roche)) on ice for 15 mins. Supernatant was collected without scraping, as previously described^41^ , to ensure minimal disruption to internal membranes and structures. For total protein extraction, cells were extracted with either RIPA buffer or UTS buffer. For lysis with RIPA buffer, cells were washed twice with ice-cold PBS and incubated with RIPA buffer (150 mM NaCl, 1% NP40, 0.5% Na-deoxycholate, 50 mM Tris pH 7.4, 5 mM EGTA, protease and phosphatase inhibitors) on ice for 15 mins. Cells were scraped, lysates collected and sonicated using a Bioruptor (Diagenode) for 2x 30s on/30s off. Lysates were centrifuged at 20,000 x g for 5 minutes and supernatant collected. For lysis with UTS buffer, cells were washed twice with room-temperature PBS, incubated with UTS buffer (7 M urea, 1% sodium deoxycholate, 20 mM tris pH 8, 5 mM TCEP, 5 mM DTT) at room temperature for 20 minutes, and scraped, sonicated, and centrifuged as above.

For mouse tissues, mice were culled by cervical dislocation at the indicated timepoints, confirmed by exsanguination, and organs (liver and brain) were dissected and flash frozen in liquid nitrogen. On dry ice, organs were chopped into ∼5 mm pieces, weighed, and 100 mg transferred to 2 ml tubes containing CK14 ceramic beads (Bertin Instruments) with 1 ml UTS buffer and homogenised using a Precellys 24 Tissue Homogeniser (Bertin Instruments) for 3 x 15 s at 5,000 RPM with 30 s breaks. Lysates were centrifuged at 20,000 x g for 1 min at 4°C, sonicated, centrifuged at 20,000 x g for 30 minutes at 4°C, and supernatants collected.

Protein concentration was determined using the bicinchoninic assay (BCA assay, Pierce) or 660 nm assay (Pierce), according to manufacturer’s instructions. Bovine serum albumin (BSA) standards were diluted in the same lysis buffer as experimental samples. Quantification was carried out in U-bottom 96-well plates (Costar) using a Tecan Spark 10M microplate reader in triplicate technical repeats. Standard curves were fitted using GraphPad Prism, and concentrations interpolated.

### Gel electrophoresis and Western blotting

Proteins were prepared for polyacrylamide gel electrophoresis (SDS-PAGE) by diluting samples in NuPage LDS sample buffer (ThermoFisher) with 5 mM DTT and heating to 70°C for 10 minutes. Samples were run on NuPage Novex 4-12% Bis-Tris protein gels (ThermoFisher) in MES buffer (Formedium).

Proteins were transferred to nitrocellulose membranes using the iBlot 2 system (ThermoFisher). Membranes were stained for total protein using Ponceau S and imaged using a ChemiDoc MP (Bio-Rad) to assess protein loading and transfer. Membranes were blocked in 5% milk (Marvel) or BSA in Tris-Buffered Saline with 0.1% Tween-20 (TBST) for 1 hour at room temperature, then incubated with primary antibody diluted in blocking buffer at 4°C overnight. After washes in TBST and incubation in appropriate HRP-conjugated secondary antibody for 1 hour at room temperature, chemiluminescence was detected with a ChemiDoc using Immobilon ECL reagent (Millipore). Densitometric analysis was performed using Fiji/ImageJ.

### Puromycin incorporation

Cells were seeded in 12-well culture plates and grown to confluence in temperature cycles. At T0, cells were given a medium change to standard cell culture medium +/-1 μM INK128 and moved to constant 37°C. At the indicated timepoints, puromycin dihydrochloride (Gibco) +/- bortezomib was added directly to cells in culture medium as a 10x bolus to a final concentration of 10 μg/ml puromycin +/- 1 μM bortezomib. Cells were labelled for 30 minutes at 37°C before lysis in RIPA buffer. Puromycin was detected by Western blotting, total protein was detected using Ponceau S, and puromycin incorporation was calculated as puromycin intensity / total protein intensity.

### Extracellular flux analysis

Oxygen consumption rate was measured using the Seahorse XFp Mito Stress Kit (Agilent) in a Seahorse XFp analyser (Agilent). Cells were seeded in Seahorse XFp cell culture miniplates and grown to confluence in temperature cycles. At T0, cells were given a medium change to standard cell culture medium with 100 nM dexamethasone +/- 1 μM INK128 and moved to constant 37°C. At least 1 hour before each assay was performed, Seahorse XFp sensor cartridges were incubated with Seahorse XF calibrant and placed in a non-CO_2_ 37°C incubator for 45-60 mins, prior to loading the sensor drug ports with the provided drugs (final concentrations oligomycin – 1.5 μM, FCCP – 2 μM, rotenone/antimycin – 0.5 μM). 1 hour before each timepoint, media was changed to bicarbonate-free medium (DMEM powder (Sigma), 10 mM glucose, 1mM pyruvate, 1% GlutaMax, osmolarity adjusted to 350 mOsm/kg with NaCl, pH adjusted to 7.6), and cells placed in a non-CO_2_ incubator at 37°C. 30 mins before each timepoint, sensor cartridges were calibrated in the Seahorse XFp analyser. At each timepoint, cells were added to the Seahorse XFp analyser and the test started.

### Live cell imaging and analysis of macromolecular crowding using genetically encoded multimeric nanoparticles

Immortalised PER2::LUCIFERASE mouse lung fibroblasts were transduced with a lentivirus expressing 40 nm genetically encoded multimeric (GEM) nanoparticles fused to fluorescent protein T-Sapphire in the cytosol (GEMs ^66,71^) at a low level via a mouse phosphoglycerate kinase 1 (PGK1) promoter and selected with blasticidin for 2 weeks to generate a polyclonal stable cell line. U-2 OS cells were similarly transduced with a lentiviral construct expressing the same open reading frame under the human elongation factor 1 alpha (EF1α) promoter and selected with blasticidin to generate a polyclonal stable cell line. Cells were seeded at high confluency in 8-well μ-slide chambers (Ibidi) pre-coated with fibronectin (Invitrogen). Cells were grown until confluent and entrained for 5 days under entraining temperature cycles (32°C : 37°C for mouse cells, or 37°C : 32°C for human cells). At T0 (at the transition from 32°C to 37°C for mouse cells, or 37°C to 32°C for human cells), the medium was changed to imaging medium (phenol-free DMEM (Gibco), 1x GlutaMax, 4.5 g/L glucose, 1x sodium pyruvate, 10% HyClone FetalClone III, 1% Pen/Strep, NucBlue Live ReadyProbes Reagent (Invitrogen)) +/- 100 nM INK128 and moved into constant 37°C and 5% CO_2_. 24h after this medium change, at the indicated timepoints, GEMs were imaged using a Nikon X1 Spinning Disk microscope with a sCMOS camera with a 100x/1.4 NA oil objective, with a time step of 30 ms for 30 s (approx. 1000 frames), excited by a 488 nm laser.

Particle tracking was performed using TrackMate^151,152^ in Fiji/ImageJ Script Editor, using an adapted Jython code. The following parameters were used: DoG detector (radius = 0.5, threshold = 0.285, spot minimum intensity > 95, spot quality filter > 0.132, with median filtering and subpixel localisation) and Kalman Tracker (linking max distance = 0.5, search radius = 0.5, max frame gap = 2, no. spots in track > 9).

For each track, the Mean Square Displacement (MSD) was calculated using the MSD Analyzer in MATLAB 2023b^153^. MSD curves were fit to a subdiffusion model captured by the function:

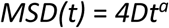

where *t* is the delay time, *a* is the power exponent of the anomalous diffusion (*a* < 1 indicates subdiffusive behaviour), *D* is the effective diffusion rate. MSD curves were filtered to retain only those with good fit (*R*^2^ > 0.9) and with *a* > 0.5. The effective diffusion coefficient (D_eff_) was calculated by fitting the first 50% of the MSD curve to the function:

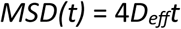

D_eff_ was averaged across the filtered tracks to give the mean across a given field of view. The mean D_eff_ per field of view was averaged for each timepoint of the circadian timecourse.

### Cell migration assay

8-well μ-slide chambers (Ibidi) were coated with fibronectin (Invitrogen). Immortalised PER2::LUCIFERASE mouse lung fibroblasts were seeded at high confluency in fibronectin-coated slide chambers and entrained for 4-7 days in temperature cycles. At the transition to 32°C (T0), cells were labelled with 1 µM CellTracker Red (ThermoFisher) for 30 minutes, followed by a medium change to “air” medium (bicarbonate-free DMEM powder (Gibco), 0.35 mg/ml NaHCO_3_, 5 mg/ml glucose, 1% Glutamax, 20 mM MOPS, 100 units/ml penicillin, 100 μg/ml streptomycin, 10% HyClone FetalClone III serum, 1 mM luciferin, adjusted to pH 7.6 and osmolality to 350 mOsm/kg using 5 M NaCl), with 10 nM INK128 or DMSO (n=4), sealed with vacuum grease, and moved to constant 37°C. At the same time, a parallel plate received the same medium change along with 1 mM luciferin and moved to an ALLIGATOR to record bioluminescence. At T24 or T36, cells were wounded by scratching with a 200 µl pipette tip and transferred to a Nikon iSIM Swept Field microscope with a 20x/0.75NA air objective and sCMOS camera held at 37°C and atmospheric CO_2_. The entire monolayer was captured every 2 hours.

The movement of cells into the wound area was monitored by measuring pixel intensity in Fiji/ImageJ and calculated by:

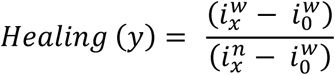

Where *i* is the mean intensity of the wound (w) or non-wound (*n*) at time *x* or *0*.

### Mouse behavioural experiments

Wild-type C57BL/6 mice were used throughout, with ages around 8 weeks old. Mice were individually housed in a temperature and humidity-controlled cabinet (Phenome Technologies), with locomotor activity monitored by wheel-running and infra-red. Mice were entrained with 12:12 hour light:dark (LD) cycles (400 lux) for at least 1 week prior to any experimental procedures and allowed *ad libitum* access to water and food throughout. Throughout experiments, mice received weekly water bottle and food changes, with 3 visual checks per week, and remote PC checks multiple times per day. Mice were weighed before, during, and after experiments. After completion of behavioural experiments, mice were culled by cervical dislocation and confirmed by exsanguination.

For the locomotor activity experiment, 12 male mice (1 control group, 2 experimental groups, N=4) were entrained under LD cycles for 1 week with 0.05% DMSO in drinking water supplemented with 10% blackcurrant and apple squash (Robinsons) to train them to the taste of the squash. After 1 week, the experimental groups were given 25 µM INK128 in diluted squash, while the control group were given 0.05% DMSO in 10% squash. After 1 week, the light schedule was changed to constant darkness (DD). After 1 week, mice were returned to DMSO diluted in diluted squash to test drug reversibility. Mice were returned to the drug under LD cycles for 3 days, and subsequently culled with brain and livers harvested and flash frozen. Tissues were processed and lysed in RIPA buffer and Western blotting performed to confirm drug efficacy. Mice were weighed every 6-8 days and observed daily for adverse effects.

### Fasting-refeeding protocol

6 WT and 6 *Per* double knock out mice (three males and three females per genotype) were placed under 12:12 hour LD cycles for 7 days. Upon transition to lights on, on day 8, food was removed from all mice. Upon transition to lights off on day 8, food was returned to the mice. Three hours later, animals were culled via cervical dislocation, confirmed by exsanguination, livers harvested and flash frozen in liquid nitrogen. Livers were chopped into ∼5 mm pieces, weighed, and 100 mg transferred to 2 ml tubes containing CK14 ceramic beads (Bertin Instruments) with 1 ml RIPA buffer: 150mM NaCl, 5 mM EDTA pH 8.0, 50 mM Tris pH 8.0, 1% NP-40, 0.5% Na-Deoxycholate, 0.1% SDS) and homogenised using a Precellys 24 Tissue Homogeniser (Bertin Instruments) for 3 x 15 s at 5,000 RPM with 30 s breaks. Lysates were centrifuged at 20,000 x g for 1 min at 4°C, sonicated, centrifuged at 20,000 x g for 30 minutes at 4°C, and supernatants collected for subsequent western blotting analysis. Phospho-4E-BP1 signal was normalised to total protein loading and analysed by two-way ANOVA with genotype and sex as factors. The study was not powered to detect sex-specific differences; the genotype × sex interaction is reported for transparency.

### Mass spectrometry (phospho)proteomics

#### Cell protein extraction

For timecourse and INK128 fibroblast and U-2 OS phosphoproteomics, cells were entrained in 12:12 hour 32:37°C or 37:32°C temperature cycles, respectively for 7 days. For the timecourse data, at T0, cells received a media change and moved to constant 37°C. 24 hours later, fibroblasts were harvested every 3 hours for 2 days in digitonin lysis buffer as above, and 200 µg protein was submitted for analysis. U-2 OS cells were sampled every 4 hours for 3 days in UTS lysis buffer and 200 µg protein was submitted for analysis.

For the INK128 phosphoproteomics experiment, at T0, cells received a media change to media +/- 1 µM INK128 and moved to constant 37°C. 24 and 36 hours later, cells were harvested in UTS buffer as above, and 50 µg protein was submitted for analysis.

### Liver and brain protein extraction

Due to cabinet space restrictions, the experiment was repeated twice consecutively and results pooled together. In each replicate experiment, 16 male mice (8 control, 8 experimental mice) were entrained under LD cycles for 7 days with 0.05% DMSO in diluted blackcurrant squash, followed by 3 days of either vehicle or 25 µM INK128 in drinking water (approximately 1 mg/kg/day). Locomotor activity was recorded throughout. Mice were culled at the indicated timepoints (2 mice per timepoint) after the transition from dark to light (zeitgeber time (ZT) = 0) starting on the 4^th^ day after INK128/vehicle treatment, with organs removed and flash frozen. Tissues were processed and Western blotted as above to check for drug efficacy. Livers were lysed in UTS buffer as above and 300 µg protein was submitted for analysis. For each experiment, a pool was made containing lysate from every replicate.

### Protein digestion

Protein samples in digitonin lysis buffer were reduced with 5 mM DTT at 56°C for 30 minutes, followed by alkylation with 10 mM iodoacetamide for 30 minutes in the dark at room temperature. Samples were digested with Lys-C (Promega) for 4 hours at 25°C (100:1 w/w) and with trypsin (Promega) at 37°C overnight (70:1 w/w). Digestion was quenched by the addition of formic acid (FA) to a final concentration of 0.5%. Precipitates were removed by centrifugation at 18000 x g for 10 minutes. Supernatants were desalted using homemade C18 stage tips containing 3M Empore extraction disks (Sigma) and 5 mg of Poros R3 resin (Thermo Scientific). The stage tips were equilibrated with 80% acetonitrile (MeCN)/0.5% FA followed by 0.5% FA. Bound peptides were eluted with 30-80% acetonitrile in 0.5% FA and lyophilized.

Protein samples from cells, mouse liver and brain in UTS lysis buffer, after dilution to 4.5M urea, were reduced with 4 mM DTT at 37 °C for 40 min and alkylated with 12 mM iodoacetamide (IAA) in the dark at room temperature for 30 min. Samples were diluted to 3M urea and digested with Lys-C (Promega) for 4 hours at 25 °C. Next, the samples were further diluted to 1.3 M urea and were digested with trypsin (Promega) over night, at 30°C. After digestion, Na-deoxycholate was removed either by acid precipitation or phase separation using ethyl acetate. For acid precipitation, the samples were acidified with FA to 0.5% so that the detergent precipitated and then separated by centrifugation from the supernatant. For phase separation, equal volume of ethyl acetate was added and acidified with formic acid (FA) to a final concentration of 0.5%. The mixtures were agitated for 2 min and centrifuged at 15,800 g for 2 mins, to completely separate the aqueous and the organic phases. After the removal of the organic phase, the aqueous phase was desalted using home-made C18 stage tips (3M Empore) filled with 5 mg Poros oligo R3 (Thermo Scientific) resin, as above.

### Tandem mass tag (TMT) labelling

From cells: lyophilized peptides were resuspended in 100 μl 200 mM HEPES. 32 μl of each TMTpro 16plex reagent (Thermo Scientific) reconstituted in 200 μl MeCN was added. Peptides from each time point were labelled with a TMT tag for 60 min at room temperature. The reaction was quenched by incubation with 5 μl 5% hydroxylamine for 30 min. The labelled peptides from all time points per experiment were combined into a single sample and partially dried to remove MeCN in a SpeedVac (Thermo Scientific). Samples were desalted as before, and the eluted peptides were lyophilized.

From mouse tissues: dried peptide mixtures (300 µg) from each condition were re-suspended in 128 μl 200 mM HEPES. 8 μl from each condition (total 32 samples) were mixed and divided to make 2 pooled samples for each set of TMT multiplex. 60 μl (1.5 mg) TMTpro 18plex reagent (Thermo Fisher Scientific), reconstituted according to manufacturer’s instructions, was added to each sample and incubated at room temperature for an hour. The labeling reaction was then terminated by incubation with 12 μl 5% hydroxylamine for 30 min. The labelled peptides were combined into two sets of TMT multiplex and were desalted using the same stage tips method as above.

### Basic pH reverse-phase fractionation

About 200 µg of the labelled peptides was separated on an off-line, high pressure liquid chromatography (HPLC). The experiment was carried out using XBridge BEH130 C18, 5 µm, 2.1 x 150 mm (Waters) column with XBridge BEH C18 5 µm Van Guard cartridge, connected to an Ultimate 3000 Nano/Capillary LC System (Dionex). Peptides were separated with a gradient of 1-90% B (A: 5% MeCN/10 mM ammonium bicarbonate, pH8; B: MeCN/10mM ammonium bicarbonate, pH 8, [9:1]) in 60 min at a flow rate of 250 µl/min. A total of 54 fractions were collected, combined into 18 fractions, and lyophilized. Dried peptides were resuspended in 1% MeCN/0.5% FA and desalted using C18 stage tips and ready for mass spectrometry analysis.

### Phosphopeptide enrichment

Phosphopeptides were enriched using TiO_2_ titansphere-chromatography (GL Science Inc. Japan). The rest of the lyophilized peptides were resuspended in a solution of 2 M lactic acid in 50% MeCN (loading buffer) and incubated at room temperature for 30 mins with TiO_2_ beads (1:5, peptides: TiO_2_ beads, w/w), that were prewashed with the loading buffer. Next, the TiO_2_ beads were centrifuged for 2 min, and the supernatant was transferred to fresh TiO_2_ beads for a second round of enrichment. After incubation, TiO_2_ beads with enriched phosphopeptides were loaded onto C8 stage tips and washed sequentially twice with loading buffer and once with 50% MeCN, 0.1% TFA. The bound phosphopeptides were eluted once with 80 μl 0.4 M ammonia solution, once with 30% MeCN, 0.4 M ammonia solution and 50% MeCN, 0.1% TFA. Samples were then acidified and partially dried down using a SpeedVac, desalted with C18 Stage tips and lyophilized. Dried phosphopeptide was fractionated using off-line high pH reverse-phase peptides fractionation same as above. The collected fractions were combined into 14 fractions and lyophilized. Dried peptides were resuspended in 30 µl 20% MeCN/0.1 % FA, MeCN was removed by vacuum centrifugation, and ready for mass spectrometry analysis.

### Mass spectrometry

LC-MS/MS analysis of TMT labelled samples were performed either on Q Exactive Plus or Orbitrap Eclipse mass spectrometers (Thermo Fisher Scientific).

For Q Exactive Plus data acquisition, the same method was used for both proteomics and phosphoproteomics. Fractionated peptides were analysed by LC-MS/MS using a fully automated Ultimate 3000 RSLC nano System (Thermo Fisher Scientific) fitted with a 100 μm x 2 cm PepMap100 C18 nano trap column and a 75 μm × 25 cm, nanoEase M/Z HSS C18 T3 column (Waters). Peptides were separated using a binary gradient consisting of buffer A (2% MeCN, 0.1% formic acid) and buffer B (80% MeCN, 0.1% formic acid) at a flow rate of 300 nl/min. The outlet of the nano column was directly interfaced via a nanospray ion source to a Q Exactive Plus mass spectrometer (Thermo Fisher Scientific). The mass spectrometer was operated in standard data-dependent mode, performing a MS full-scan in the m/z range of 380-1600, with a resolution of 70000. This was followed by MS2 acquisitions of the 15 most intense ions with a resolution of 35000 and CE of 33%. MS target values of 3e6 and MS2 target values of 1e5 were used. The isolation window of precursor ion was set at 0.7 Da and sequenced peptides were excluded for 40 seconds.

For LC-MS/MS carried out on Orbitrap Eclipse, the Ultimate 3000 RSLC nano System was fitted with a PepMap Neo C18 5 μm 0.3 x 5 mm nano trap column (Thermo Fisher Scientific) and an Aurora Ultimate TS 75 μm x 25 cm x 1.7 μm C18 column (IonOpticks). Peptides were separated using buffer A (0.1% FA) and buffer B (80% MeCN, 0.1% FA) at flow rate of 300 nl/min and column temperature of 40 °C. Eluted peptides were introduced directly via a nanoFlex ion source into an Orbitrap Eclipse mass spectrometer (Thermo Fisher Scientific). The mass spectrometer was operated in real-time database search (RTS) with synchronous-precursor selection (SPS) -MS3 analysis for reporter ion quantification. MS1 spectra were acquired using the following settings: resolution of 120K; mass range of 400-1400 m/z; AGC target of 4e5; MaxIT of 50 ms and dynamic exclusion was set at 60 s. MS2 analysis were carried out with HCD activation, ion trap detection, AGC of 1e4; MaxIT of 50ms; collision energy (CE) of 33% and isolation window of 0.7 m/z. RTS of MS2 spectrum was set up to search UniProt *Mus musculus* proteome (2021), with fixed modifications cysteine carbamidomethylation and TMTpro 16plex at N-terminal and Lys residue. Met oxidation was set as variable modification. Missed cleavage of 1 and maximum variable modifications of 1. In MS3 scans, the selected precursors were fragmented by HCD and analysed using the orbitrap with these settings: isolation window of 0.7 m/z; CE of 55, orbitrap resolution of 120K; scan range of 110-450 m/z; MaxIT of 250 ms and AGC of 1.5e5.

For phosphoproteomics, multistage activation (MSA)-RTS-SPS-MS3 method was carried out on Orbitrap Eclipse. MS1 scans were using the same parameters as above except MaxIT was 75ms. MS2 analyses were carried out with CID MSA activation on neutral loss mass 97.9673; detection in the Orbitrap; orbitrap resolution of 60K; AGC of 1e5; MaxIT of 118ms; CE of 35% and isolation window of 0.7 m/z. RTS of MS2 spectrum was set up as proteomics, with inclusion of phospho STY as variable modification and maximum variables of 3. In MS3 scans, the selected precursors were fragmented by HCD and analysed using the orbitrap with these settings: Isolation window of 0.7 m/z; CE of 55, orbitrap resolution of 120K; scan range of 110-450 m/z; MaxIT of 400 ms and AGC of 1.5 e5.

### Raw MS data processing

The acquired LC-MS/MS raw files were processed using MaxQuant^154^ with the integrated Andromeda search engine (v.1.6.17.0 or 2.4.2.0). MS/MS spectra were quantified with reporter ion MS2 or MS3 from TMTpro 18-plex experiments and searched against *Mus musculus* reviewed UniProt Fasta database (downloaded in Nov 2019 or Nov 2020). Carbamidomethylation of cysteines was set as fixed modification, while methionine oxidation, N-terminal acetylation (protein), and STY phosphorylation (for phosphoproteomics) were set as variable modifications. Protein quantification requirements were set at 1 unique and razor peptide. Other parameters in MaxQuant were kept as default values. The MaxQuant output file was then processed with Perseus software (v. 1.6.15.0 or 2.0.10.0). After uploading the matrix, the data was filtered, to remove identifications from reverse database, identifications with modified peptide only, and common contaminants. The localization probability of the phospho(STY).txt file was also filtered to ≥0.75. Multiply phosphorylated peptides were retained where phosphosite localization probability was ≥0.75, and reported phosphosite numbers include confidently localized sites derived from both singly and multiply phosphorylated peptides, but residue-specific interpretations were restricted to confidently localized sites. Both sets of data with the reporter intensities of ‘0’ were converted to NaN and exported as text files for further data analysis.

### Recombinant PER2 and mTORC1 protein expression and purification

The mTORC1 complex was expressed (via co-expression of mTOR, RAPTOR and mLST8) by transient transfection of Expi293F cells. Cells were transfected at a density of 2.5 x10^6^ cells/mL with a total of 1.1 mg DNA and 3 mg Polyethyleneimine "MAX" (Polysciences) per 1 L of cells. After 52 h, cells were harvested by centrifugation and cell pellets were frozen in liquid N_2_. mTORC1 was purified from 2L Expi293F cells as described previously ^98^ by affinity purification on StrepTrap HP resin (Cytiva), followed by Strep-tag cleavage by TEV protease (purified in house by the Williams lab) overnight on the column. The cleaved protein was further purified by anion-exchange chromatography (AEX) on a 5 mL HiTrap Q column (Cytiva), concentrated with Amicon Ultra-4 100 kDa concentrators (Merck Millipore), flash frozen in liquid N_2_ and stored at -80 °C.

Full length, human PER2 tagged with 2x Strep-tag was newly synthesized as a plasmid in a pRP mammalian expression plasmid (VectorBuilder) and was expressed by transient transfection of Expi293F cells grown in Expi293 media (Gibco) in a Multitron Pro shaker set at 37 °C, 8% CO2 and 125 RPM. After 48 h, cells were harvested by centrifugation and cell pellets frozen in liquid N_2._ Full length, human PER2 tagged with 2x Strep-tag was purified from 1 L Expi293F cells by affinity purification on a StrepTrap HP resin (Cytiva). Cell pellets were lysed in a lysis buffer consisting of 50 mM Tris-HCl pH 7.5, 300 mM NaCl, 2 mM MgCl_2_, 0.5 mM TCEP, 10% glycerol. Following sonication, the clarified lysate was loaded onto a tandem StrepTrap HP 5ml column. The column was washed with 20 column volumes (CV) of lysis buffer followed by 20 CV wash buffer (50 mM Tris-HCl pH 7.5, 200 mM NaCl, 2 mM MgCl_2_, 0.5 mM TCEP, 10% glycerol) and protein was eluted with wash buffer supplemented with 10 mM desthiobiotin (Sigma).

Fractions containing PER2 were pooled and cleaved overnight by adding TEV protease. The TEV-cleaved sample was subjected to Ni-NTA chromatography using a HisTrap HP column (Cytiva) to capture the His_6_-TEV protease. The flow-through was collected and concentrated. The cleaved PER2 protein was further purified by anion-exchange chromatography (AEX) on a 5 mL HiTrap Q column (Cytiva). The protein was further purified by a size-exclusion chromatography using Superdex 75 16/60 column (Cytiva), and fractions containing PER2 were concentrated with an Amicon Ultra-4 10 kDa concentrator (Merck Millipore), flash frozen in liquid N_2_ and stored at -80 °C.

### In vitro Strep-tag pull-down assays

Strep-tagged mTORC1 complex, or Strep-tagged RAPTOR at a final concentration of 0.5µM was incubated for 30 min on ice with StrepTactin Sepharose High Performance beads (GE Healthcare). The pulldown reaction buffer was 50 mM HEPES pH 7.5, 100 mM NaCl, 1 mM TCEP, 5 mM MgCl_2_, 10% glycerol. The protein-bound beads were washed twice with reaction buffer, after which 4 µM PER2 was added. Samples of input reactions were taken, before incubating on ice for 90 min. Thereafter, the beads were washed 10 times in the same reaction buffer, and the samples were taken and heated to 70 °C for 5 min after addition of NuPAGE LDS sample buffer (ThermoFisher) to a final 1x concentration. The control experiment was performed with PER2 that had no Strep-tag but was incubated with StrepTactin Sepharose High Performance beads and washed in the same way as the above reactions. The samples were analysed by Coomassie-stained SDS-PAGE gel. Reactions were performed in three independent experiments.

### Cross-linking Mass Spectrometry

Protein cross-linking reactions were performed for 60 min at room temperature using purified PER2 and mTORC1 complexes at final concentrations of approximately 20 µM and 1.5 µM, respectively. The proteins were prepared in 50 mM HEPES, pH 7.5, 100 mM NaCl, 1 mM TCEP, 5 mM MgCl₂, and 10% glycerol, and cross-linking was initiated by adding EDC to a final concentration of 10 mM. Crosslinked protein was quenched with the addition of Tris pH 7.4 to a final concentration of 50 mM. The quenched solution was reduced with 5 mM DTT and alkylated with 20 mM iodoacetamide. SP3 protocol as described previously ^155,156^ was used to clean-up and buffer exchange the reduced and alkylated protein. Proteins were washed with ethanol using magnetic beads for protein capture and binding. The proteins were resuspended in 100 mM NH_4_HCO_3_ and were digested with trypsin (Promega) at an enzyme-to-substrate ratio of 1:25, and protease max 0.1% (Promega). Digestion was carried out overnight at 37 °C. Clean-up of peptide digests was carried out with HyperSep SpinTip P-20 (ThermoScientific) C18 columns, using 60% acetonitrile as the elution solvent. Peptides were then evaporated to dryness *via* Speed Vac Plus (Savant).

Dried peptides were resuspended in 30% acetonitrile and were fractionated *via* size exclusion chromatography using a Superdex 30 Increase 3.2/300 column (GE Healthcare) at a flow rate of 20 μL/min using 30% (v/v) ACN 0.1 % (v/v) TFA as a mobile phase. Fractions were taken every 5 minutes, and the 2nd to 7th fractions containing cross linked peptides were collected. Dried peptides were suspended in 3% (v/v) acetonitrile and 0.1 % (v/v) formic acid and analysed by nano-scale capillary LC-MS/MS using an Ultimate U3000 HPLC (ThermoScientific, USA) to deliver a flow of 300 nl/min. Peptides were trapped on a C18 Acclaim PepMap100 5 μm, 0.3 μm x 5 mm cartridge (ThermoScientific, USA) before separation on PepMap RSLC C18, 2 μm, 100 A, 75 μm x 50 cm EasySpray column (ThermoScientific, USA). Peptides were eluted on optimised gradients of 90 minutes and interfaced *via* an EasySpray ionisation source to a tribrid quadrupole Orbitrap mass spectrometer (Orbitrap Eclipse, ThermoScientific, USA) equipped with FAIMS. MS data were acquired in data dependent mode with a Top-25 method, high resolution full mass scans were carried out (R = 120,000, *m/z* 400 – 1550) followed by higher energy collision dissociation (HCD) with stepped collision energy range 21, 30, 34 % normalised collision energy. The tandem mass spectra were recorded (R=60,000, isolation window *m/z* 1, dynamic exclusion 50 s). Mass spectrometry measurements were cycled for 3 s durations between FAIMS CV -45, and -60 V.

#### Crosslinking data analysis

Xcalibur raw files were converted to MGF files using ProteoWizard^157^ and cross links were analysed by XiSearch^158^. Search conditions used 3 maximum missed cleavages with a minimum peptide length of 5. Variable modifications used were carbamidomethylation of cysteine (57.02146 Da) and Methionine oxidation (15.99491 Da). False discovery rate was set to 5%. Crosslinks were illustrated using xiView.

### Cell RNA stain microscopy

#1.5 glass coverslips (VWR) were coated with 1% bovine fibronectin (Sigma) diluted in PBS for 1 hour at 37°C. Cells were seeded onto coverslips and entrained using temperature cycles. At T0, cells received a final medium change +/- 1 µM INK128 and were moved into constant 37°C. At the indicated times, cells were fixed in ice-cold methanol at -20°C for 10 minutes. After all timepoints, cells were stained together with 1 µM SYTO RNASelect Green Cell Stain (Invitrogen) for 20 minutes at room temperature, followed by washes with PBS and mounting onto glass slides using ProLong Gold with DAPI (Invitrogen).

Cells were imaged using a Zeiss 780 inverted confocal microscope with a 63x/1.4NA oil objective with 2 µm z-stacks and 4x4 binning. Maximum intensity z-projections were analysed using CellProfiler v4.0.7^159^ . Individual cells were identified through Otsu thresholding of the nuclear channel (DAPI), followed by the watershed algorithm to identify cell edges in the RNA channel (SYTO), where the area of lowest inverted intensity is used to detect the boundaries between cells. Nuclei in the SYTO channel were identified through masking of the DAPI channel.

### Liver RNA stain microscopy

#### Mouse cardiac perfusion

Due to cabinet space restrictions, the experiment was repeated twice consecutively and results pooled together. In each replicate experiment, 16 male and female mice (8 control, 8 experimental mice) were entrained under 12h:12h LD cycles for 7 days with 0.05% DMSO in diluted blackcurrant squash, followed by 3 days of either vehicle or 25 µM INK128 (approximately 1 mg/kg/day). Locomotor activity was recorded throughout. At the indicated timepoints, mice were administered a lethal dose of 100 mg/ml sodium pentobarbital diluted 1:1 in PBS by intraperitoneal injection at 10 ml/kg. Once mice were checked for blinking and pedal reflexes, hearts were exposed, and left ventricle flushed 2x with 10 ml PBS with a 21G butterfly needle. Mice were then flushed 2x with 20 ml cold 4% PFA in 0.1 M phosphate buffer (19.22 g/L Na_2_HPO_4_.2H_2_O, 3.952 g/L NaH_2_PO_4_.2H_2_O). Tissues were removed and stored in PBS on ice.

#### Liver cryosectioning and staining

Livers from perfused/fixed mice were fixed in 4% PFA overnight at 4°C and incubated in 30% sucrose in PBS overnight at 4°C. Livers were cut into lobes, embedded in gelatin (7.5% gelatin, 10% sucrose in PBS), plunge frozen in 2-methylbutane (Sigma) at -50°C and cryosectioned at a thickness of 20 μm on a cryostat (Leica). Sections were stained with 1 µM SYTO RNASelect Green for 1 hour at room temperature, followed by washes with PBS and mounted onto SuperFrost Plus slides (VWR) with #1 rectangular coverslips (Menzel) using VECTASHIELD Hardset Antifade Mounting Medium with DAPI (2B Scientific).

### Imaging of liver slices

Liver slices were imaged using a Nikon X1 Spinning Disk microscope with a sCMOS camera with a 20x/0.75NA air objective. Nuclei masks were segmented using the Fiji/ImageJ plugin for StarDist ^160,161^, using the Versatile (fluorescent nuclei) training model. Measurement of nuclei intensity was carried out in Fiji/ImageJ. Background fluorescence was calculated by excluding the nuclei ROIs from the fluorescence across the entire image.

### Immunostaining of mouse brain slices

#### Preparation of brain slices

Mice were deeply anaesthetized with urethane (0.5 ml/hg, 20% solution in saline, i.p. (Sigma)) and transcardially perfused with 0.9% saline, followed by 4% paraformaldehyde in 0.1 M PBS, pH 7.4. Brains were removed and postfixed in 4 % paraformaldehyde overnight at 4°C. Brains were sectioned at 60 µm thickness using a vibratome (Leica) and washed in PBS. Sections were immersed in blocking solution (1% Triton X-100, 3% BSA in PBS), for 1 hour at room temperature, then incubated with primary antibodies diluted at 1:1,000 overnight at 4°C. Sections were incubated with Alexa Fluor 488-conjugated secondary antibodies diluted at 1:100 for 2 hours at room temperature. After PBS washes, sections were mounted onto slides using VECTASHIELD (Vector Laboratories).

#### Imaging of brain slices

Sections immunostained for pS6 were imaged using a confocal microscope (Leica SP5) equipped with a 20× oil immersion objective (Leica N PLAN EPI, NA 0.40 OIL POL XLR). Fluorescence was collected using a 510–570 nm emission filter. Images were acquired at a resolution of 512 × 512 pixels, and adjacent fields were collected as tiled images to encompass the entire thickness of the visual cortex. Each final image represented the average intensity projection of six consecutive optical sections acquired along the z-axis.

#### Mouse physiological monitoring

Mice were pretreated with 25 µM INK128 in drinking water (approximately 1 mg/kg/day) or vehicle for 3 days, then were transferred to individually housed indirect calorimetry cages (CLAMS, Columbus Instruments) and monitored for 4 days. The first day was excluded from analysis to allow for acclimatisation. O_2_ consumption and CO_2_ production were recorded every 10 minutes. Respiratory exchange ratio was calculated as VCO_2_/VO_2_, and energy expenditure calculated as 3.815 × VO_2_ + 1.232 × VCO_2_.

### Mouse drug metabolism experiment

#### Tissue collection

16 male and female mice were entrained under 12h:12h LD cycles for 7 days with 0.05% DMSO in diluted blackcurrant squash, followed by 3 days of either vehicle or 25 µM INK128 in drinking water (approximately 1 mg/kg/day). Locomotor activity was recorded throughout. At ZT9 and 21, 4 mice per condition were intraperitoneally injected with 4 mg/kg bupropion hydrochloride and 2 mg/kg atorvastatin in PBS (4 μl/g volume) and returned to the cage. 30 minutes later, under isoflurane, 500 μl blood was drawn by cardiac puncture in 1 ml MiniCollect tubes with K3EDTA (Greiner Bio-One). Blood plasma was collected by centrifugation at 2000 x g for 10 minutes and frozen on dry ice. Livers were dissected and flash frozen in liquid nitrogen, then chopped on dry ice, transferred into Precellys CK14 homogenising tubes, and around 100 mg homogenized in 4 volumes of Milli-Q water using a Precellys homogenizer on wet ice for three cycles, each consisting of 2 × 20 s at 5500 rpm.

#### Mass spectrometry for metabolites

Liver homogenates and plasma samples were prepared for LC-MS/MS analysis using the same extraction procedure. Briefly, 50 µL of sample was transferred to a 500 µL low-binding plate and mixed with 180 µL acetonitrile containing 0.2% formic acid and 10 nM DIDB as internal standard. Samples were vortex-mixed and centrifuged at 4000 × g for 20 min at 4°C. An aliquot of 75 µL of the resulting supernatant was transferred to a fresh low-binding plate and diluted with 75 µL Milli-Q water. Plates were sealed and centrifuged again at 4000 × g for 10 min at 4°C prior to analysis.

LC-MS/MS analysis was performed using an Acquity UPLC system coupled to a Xevo TQ-XS tandem quadrupole mass spectrometer (Waters, Milford, MA, USA) operated with electrospray ionization. Chromatographic separation was achieved on a Waters Acquity UPLC HSS T3 column (50 × 2.1 mm, 1.8 µm) maintained at 40°C. The injection volume was 1 µL. The mobile phases consisted of 0.2% formic acid in Milli-Q water (A) and 0.2% formic acid in acetonitrile (B), delivered at a flow rate of 1.0 mL/min. The gradient elution program was: 0– 0.3 min, 0.2% B; 0.3–1.3 min, increased to 95% B; 1.3–1.8 min, held at 95% B; 1.8–1.81 min, returned to 0.2% B; and 1.81–2.3 min, re-equilibrated at 0.2% B.

Instrument control and data acquisition were performed using MassLynx v4.2, and peak integration was conducted using TargetLynx v4.2 (Waters). Detection was performed in multiple reaction monitoring (MRM) mode using the following transitions: atorvastatin, 559.30 → 440.15; hydroxyatorvastatin, 575.27→439.98; bupropion, 239.80→130.94; and hydroxybupropion, 255.99→130.94. Concentrations were quantified against calibration standards prepared in plasma and liver homogenate, respectively.

### Investigation of sleep-wake architecture in mice

#### Surgical procedure for EEG electrodes implantation

Animals were anesthetized with a mixture of Ketamine (Nimatek, 100mg/mL; Dechra, Norwich, UK) and Rompun (Xylazine 2%, 0.06 mL/kg; Bio98, Milan, Italy) and placed on a heating blanket in a custom stereotaxic frame. Two small holes (∼1.2 mm diameter) were made in the skull to allow the insertion of two custom-made iron-plated, round-tipped miniature screws that served as EEG electrodes. Electrodes were placed over the right parietal lobe (3 mm lateral to midline, 2 mm posterior to bregma) and the right frontal lobe (1.5 mm lateral to midline, 1.5 mm anterior to bregma). A third screw, used as a reference electrode, was fixed on the skull above the cerebellum (2 mm posterior to lambda, on the midline). Two small stainless-steel wires (Advent) were inserted into the neck muscles to record the electromyogram (EMG). Electrodes were soldered to stainless steel wires and secured to the skull with dental cement (Paladur, AgnTho’s, Lidingö, Sweden). Mice were immediately treated with post-operative analgesia (Tramadol 10 mg/kg, Formevet, Milan, Italy). Upon waking up from the anaesthesia, animals were placed in individual cages for seven days to allow complete recovery before initiation of EEG recordings. Surgical procedures were performed between 10.00 am and 12.00 pm.

### Data acquisition

The EEG signal was recorded using the Micromed Brain Quick LTM Holter EEG system (Micromed, Mogliano Veneto, Italy). Data were acquired at 256 Hz and band-pass filtered (0.5 Hz to 100 Hz). Mice were allowed to move freely within a Plexiglas arena (35 × 35 × 40 cm) placed inside a Faraday cage. Before the start of each recording session, mice were habituated to the apparatus for approximately 30 min. EEG recordings were performed for 3 consecutive days before INK128 administration (baseline) and for 3 consecutive days during INK128 treatment, in a quiet, temperature-controlled room with a 12h light/dark cycle (lights on from 6:00 AM to 6:00 PM). Animals were inspected daily to monitor their well-being, verify the integrity of the EEG implant, and ensure proper signal acquisition. After each recording session, the bedding was removed, the arena was cleaned with 70% ethanol, and fresh bedding was added before the next recording.

### Zebrafish behavioural tracking

Zebrafish embryos were collected after lights on from spawning adults (strain TLxAB) from the University College London Fish Facility and raised in a 28°C incubator with a 14hr:10hr light:dark cycle. Following swim bladder inflation at 4dpf, larvae were individually placed in a 96-well square well plate with ∼650 µL of fish water (0.3 g/L Instant Ocean and 1 mg/L methylene blue [pH 7.0]) and tracked with infrared light on 14hr:10hr LD cycles (550 lux) in a Zebrabox (ViewPoint Behavior Technology) using the quantized mode of the ZebraLab software (ViewPoint Behavior Technology), as in ^162^.

For the INK128 dose response curve, a 100 mM stock solution of INK128 dissolved in DMSO was serially diluted to 500x for each dose, such that 1.3 µL was added to each 650 µL well to achieve the final test concentrations of 0.1, 0.5, 1, 2, 10, and 20 µM (0.2% DMSO). Drugs were added by pipette directly to each well between ZT9-10 at 5 dpf (for the dose response curve) or 7 dpf (for the retesting of 10 µM) during the behavioural tracking. At the end of the tracking experiment, prior to data analysis, larvae were visually inspected for any abnormalities, and damaged or sick larvae were excluded (only 1 larva, from the 0.1 µM condition, was excluded in this manner).

### *Arabidopsis thaliana* bioluminescence imaging

Prior to bioluminescence imaging and sampling, seedlings were treated at dawn with either 10 µM INK128 (Cell Signaling Technology Europe) or equivalent volume of vehicle, combined with 5 mM D-luciferin potassium salt. Four plates were opened under sterile conditions, and seedlings were frozen in liquid N_2_. One plate was placed under a Photek HRPCS intensified CCD camera (Photek, Hastings, UK) for six days, with 800 s bioluminescence integrations every hour. During imaging, plants experienced 12 h of light and 12 h darkness for the first 24 h, followed by 5 days under free-running conditions of constant light at 19°C. Irradiance during imaging was around 50 μmol m^-2^ s^-1^, delivered from combined red / blue LED panels (660 nm and 470 nm). Bioluminescence data were extracted using the Photek Image32 software (Photek, Hastings, UK). For analysis, only the five days under free-running conditions were considered.

### *Neurospora crassa* bioluminescence imaging

In vivo bioluminescence assays were performed in 96-well plates containing LNN-CCD medium (0.03% glucose, 0.05% arginine, 50 ng/ml biotin, 1.5% agar, and 25 μM luciferin) supplemented with 0.01 M quinic acid (QA). Cultures were entrained for three days under 12 h light:12 h dark (LD 12:12) cycles before being transferred to constant darkness (DD) at 25°C for bioluminescence monitoring. INK128-treated cultures were supplemented with the drug to a final concentration of 10 μM, whereas mock-treated controls received an equivalent volume of DMSO. Data acquisition and analysis were performed as previously described ^163^.

## QUANTIFICATION AND STATISTICAL ANALYSIS

### Circadian data analysis

Data from bioluminescence assays were measured in Fiji/ImageJ v2.3^164^, quantified by integrated density, and analysed in GraphPad Prism v9. Data analysis began from T24, to exclude the transient impact of serum or drug additions on clock gene expression. Data was detrended using a 24-hour moving average in Excel or R v4.2.1. A circadian damped cosine wave was fitted by least-squares to calculate period, damping rate and amplitude:

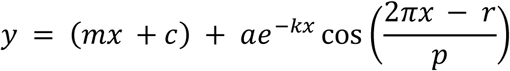

Where *m* is the baseline, *c* is the displacement from the *y*-axis, *k* is the damping rate, *a* is the amplitude, *r* is the phase, and *p* is the period. Rhythmicity was calculated by comparing the fit of this equation to the null hypothesis of a straight line using the Extra sum-of-squares F-test.

For circadian phase calculations, area under the curve analysis was performed to identify the time of the first circadian peak. Circadian phase was calculated in Excel as:

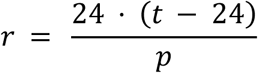

Where *r* is the phase, *t* is the time of the first peak, and *p* is the period. Phase shifts were calculated as mean untreated control phase subtracted from experimental phase then transformed (*y* = 0 – *y*) to assign phase advances (positive phase shifts) and phase delays (negative phase shifts). Mean phase shifts and time of treatment administration were used to generate a phase response curve for each treatment. Relative amplitude comparisons for core vs output rhythms were performed by fitting a standard sine wave to data in a time window of T36-48 to T72-84 and dividing sine-fitted amplitude by mean bioluminescence during the time window.

### Analysis of timecourse proteomics data

All bioinformatics analysis of timecourse (phospho)proteomics data was done using R v4.5.2.

#### Sample normalisation

Only proteins/phosphosites present in all samples and pools (if applicable) were included for analysis. Sample loading normalisation was carried out by normalising intensities to the mean of the sum of all intensities at each timepoint. TMTpro 18-plex experiments were additionally normalised by internal reference scaling^165^, where the arithmetic mean abundance for each protein in both pools were calculated, then the mean of the means was calculated and used to normalise protein values for each sample.

#### Rhythmicity analysis

For fibroblast timecourse phosphoproteomics, rhythmicity analysis was carried out using RAIN ^166^ with a cut-off of Benjamini-Hochberg corrected FDR < 0.05.

For two timepoint proteomic and phosphoproteomic fibroblast data, temporal variation was assessed separately in vehicle and INK128-treated fibroblasts using two-way repeated measures ANOVA with time as a factor. Multiple testing correction was applied using the two-stage step-up method of Benjamini, Krieger, and Yekutieli, with a false discovery rate threshold ≤ 0.05. Proteins and phosphosites were considered mTOR-dependent if they showed significant temporal variation in vehicle-treated cells but not in INK128-treated cells.

For liver and brain data, rhythmicity analysis was carried out using eJTK Cycle^167^ performed with BioDare2^109^ , with no detrending and “eJTK Classic” preset parameters, and a cut-off of Benjamini-Hochberg-(BH) corrected FDR ≤ 0.05 was used. For amplitude and phase calculation, the MFourFit algorithm was used: data was concatenated, with period constrained to 23.5-24.5 hours, and for phase analysis, data was log_2_ transformed.

#### Gene ontology analysis

Ranked gene ontology molecular function analysis was performed using the GOrilla tool ^168^ using a background of all detected proteins. GO terms were simplified using REViGO^169^.

#### Kinase enrichment analysis

Data was segregated into four tables according to their phase, where sites peaking at 0 to 6, 6 to 12, 12 to 18 and 18 to 24 hours were concatenated. To calculate kinase enrichment for phosphorylated residues in each of the 4 datasets, the column ‘sequence window’ containing the sequence of the 15 amino acids both up-and downstream of the phosphorylated residue was imported into the Kinase Library ^170,171^. Sequences of rhythmic sites within each table were inserted as foreground and all phosphorylated residues quantitated in the experiment were used as background. Enrichment value for each kinase were log_2_ transformed and plotted along with the -log_10_ transformed p-value of that enrichment.

### Proteomics data analysis for differential protein and phosphopeptide abundance in PER2-HALO cells

#### Differential protein and phosphopeptide abundance analysis

Proteomics data analysis for differential protein and phosphopeptide abundance was performed in R (v4.5.2) using the QFeatures ^172^ (v1.20.0) and biomasslmb ^173^ (v0.0.4) packages. All analysis code is provided as R Markdown notebooks available from https://github.com/lmb-mass-spec-compbio/PER2_HALO_PROTAC v 0.1 and archived on Zenodo.

#### PER2-HALO immunoprecipitation analysis

Analysis of PER2–HALO immunoprecipitation data was conducted using peptide-level output (modificationSpecificPeptides.txt) from MaxQuant. Peptides matching contaminants and peptides derived from proteins with Q-values ≥ 0.01 were removed. Remaining peptides were log2-transformed and median-normalised using histone-derived peptides only. The limpa package ^174^ (v1.0.6) was used to fit a detection probability curve and to estimate protein-level abundances with associated uncertainties. These abundance estimates were subsequently analysed using limma^175^ (v3.64.3) to identify differences between PER2–HALO and HALO samples. P-values were adjusted using the Benjamini–Hochberg false discovery rate (FDR) procedure ^176^, and an FDR threshold of 0.05 was used to define statistical significance.

#### PER2-HALO PROTAC analysis

Analysis of PER2–PROTAC data was performed using peptide spectrum match (PSM)-level output (evidence.txt) from MaxQuant. PSMs matching contaminants and PSMs from proteins with Q-values > 0.01 were excluded. Phosphopeptides were further filtered to retain only PSMs in which all phosphosite (STY) localization probabilities exceeded 0.5. PSMs with evidence of high co-isolation interference were also removed: MaxQuant’s Parent Ion Fraction (PIF) — the proportion of MS1 ion current within the isolation window attributable to the target peptide — was used to exclude PSMs with PIF < 0.5, while PSMs for which PIF could not be calculated (no MS1 isotope pattern assembled) were retained. PSM intensities were median-normalised. PSMs with more than 3 of 18 missing values were removed, and remaining missing values were imputed using QFeatures::impute with method = “MinProb”. Protein- and phosphopeptide-level abundances were obtained by summing PSM-level intensities and then log2-transformed. To assess changes in phosphorylation independent of changes in total protein abundance, protein and phosphopeptide abundances were modelled jointly using limma^175^ (v3.64.3) with an interaction model of the form condition + type + condition:type, where type denotes protein or phosphopeptide. The interaction term captures changes in phosphorylation relative to changes in protein abundance. Because protein and phosphopeptide measurements were derived from the same biological samples, sample identity was included as a blocking factor in limma::lmFit, and limma::duplicateCorrelation was used to estimate intra-block correlation. P-values were adjusted using the Benjamini–Hochberg FDR procedure^176^, and an FDR threshold of 0.1 was used to define statistical significance.

### Mouse behavioural analysis

#### Locomotor activity

Blinded assessment of phase shift amd period were determined using ClockLab software (Actimetrics). Period and summed counts were calculated from the time window when mice were in constant darkness, excluding the first day of recording. Activity profile data was calculated binning average activity counts across the recording window in LD or DD (excluding the first day of recording) into 1-minute intervals, then expressed as % activity by normalising binned activity to the sum of all counts.

#### Sleep staging and EEG analysis in mice

The three behavioural states (i.e., wakefulness, nREM and REM sleep) were scored in epochs of 4 seconds, by visual inspection of the EEG and EMG signals using the Sirenia Sleep software (Pinnacle Technology). Epochs were retained for sleep scoring even when only a single EEG channel contained a clean, artefact-free electrophysiological signal in conjunction with the EMG. However, such epochs were excluded from subsequent spectral analyses, which were performed only on epochs in which both the frontal and parietal EEG channels exhibited clean, artefact-free signals.

The EEG recordings, together with the corresponding sleep scores, were imported into MATLAB (MathWorks Inc.) for further processing. The percentage of each behavioural state was calculated by summing all epochs assigned to that state and expressing the total as a percentage of the 24-hour recording period (24 h = 100%).

Sleep "bouts" were defined as periods comprising at least three consecutive epochs of the same behavioural state. Power spectral densities (PSDs) of the EEG signals were calculated using Welch’s method with a 0.5-second Hann window and 0.25-second overlap between consecutive windows, as implemented in the MATLAB pwelch function. Two subjects were excluded from the spectral analysis at this stage due to insufficient artefact-free EEG data.

Unless differently specified, statistical analyses were conducted using a Wilcoxon signed rank test (‘signrank’ function on MATLAB).

### Analysis of zebrafish behavioural data

Behavioural data for each larva was analysed using custom-written Matlab scripts (available on Github: https://github.com/JRihel/Sleep-Analysis). A zebrafish sleep bout was defined as any continuous period of inactivity lasting 1 minute or longer, the duration of which was defined as the ‘sleep bout length’ ^177^.

### Statistics

Data presented as mean ± SEM (indicated in figure legends), and statistical significance of mean differences determined using t-test, one-way ANOVA (OWA) and two-way ANOVA (TWA) as indicated in the figure legend, with the multiple comparisons test (MCT) used also reported in the figure legend. TWA_int_ indicates the ANOVA of interaction. The sample size (n) is also reported in the figure legend for each experiment, with n defined as the number of identically treated replicates. N refers to the number of biological replicate mice or zebrafish used for each experiment. Comparison of line fits for PRC was performed using the extra sum-of-squares F test comparing the null hypothesis of a first-order polynomial to a third-order polynomial. For analysis of EEG data, statistical analyses were conducted using a Wilcoxon signed rank test (‘signrank’ function on MATLAB).

Statistical analyses were performed using R v4.6.0, Graphpad Prism v11, or MATLAB R2025. P values are reported using the following symbolic representation: ns = p > 0.05, * = p ≤ 0.05, ** = p ≤ 0.01, *** = p ≤ 0.001, **** = p ≤ 0.0001.

## KEY RESOURCES TABLE

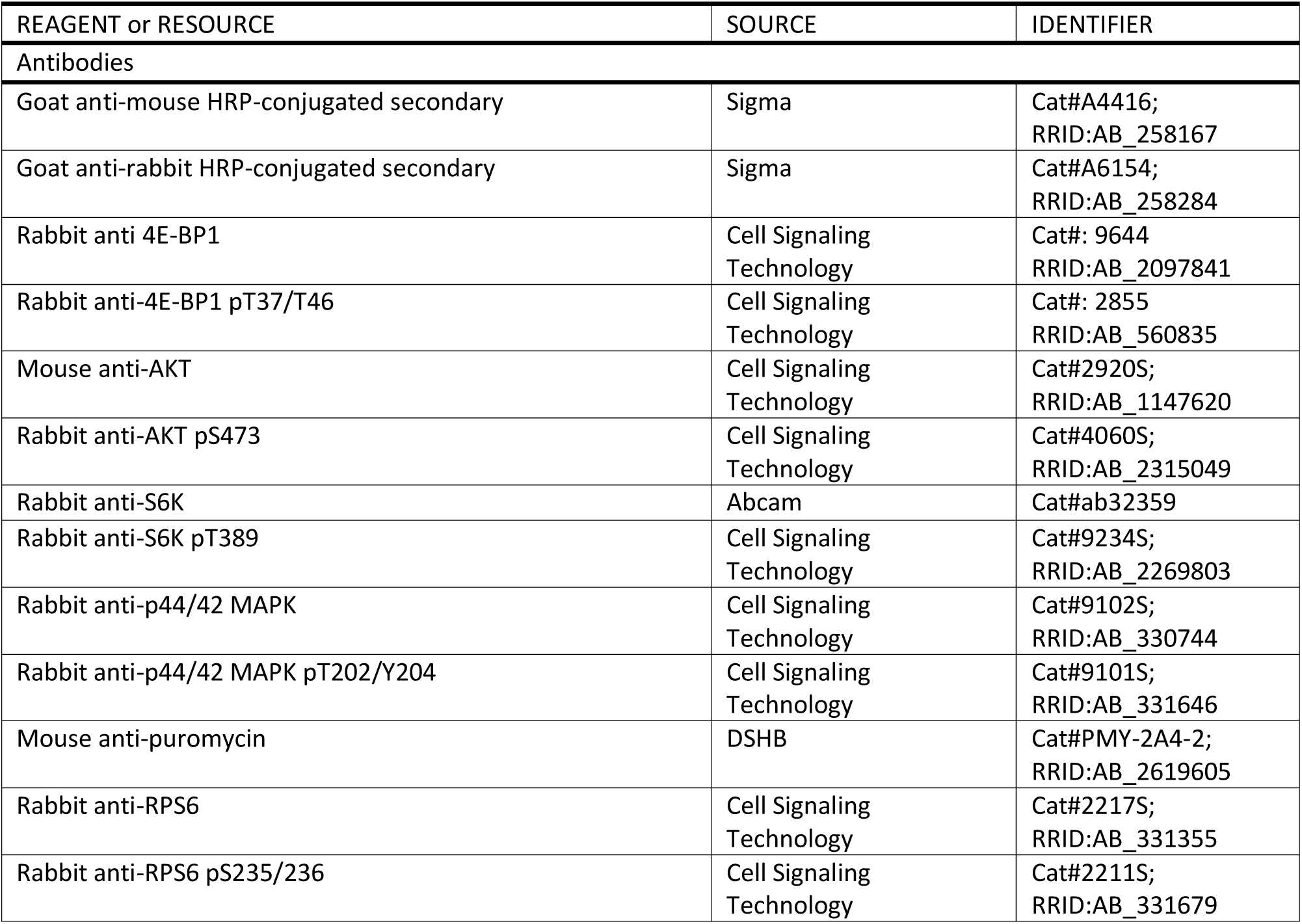
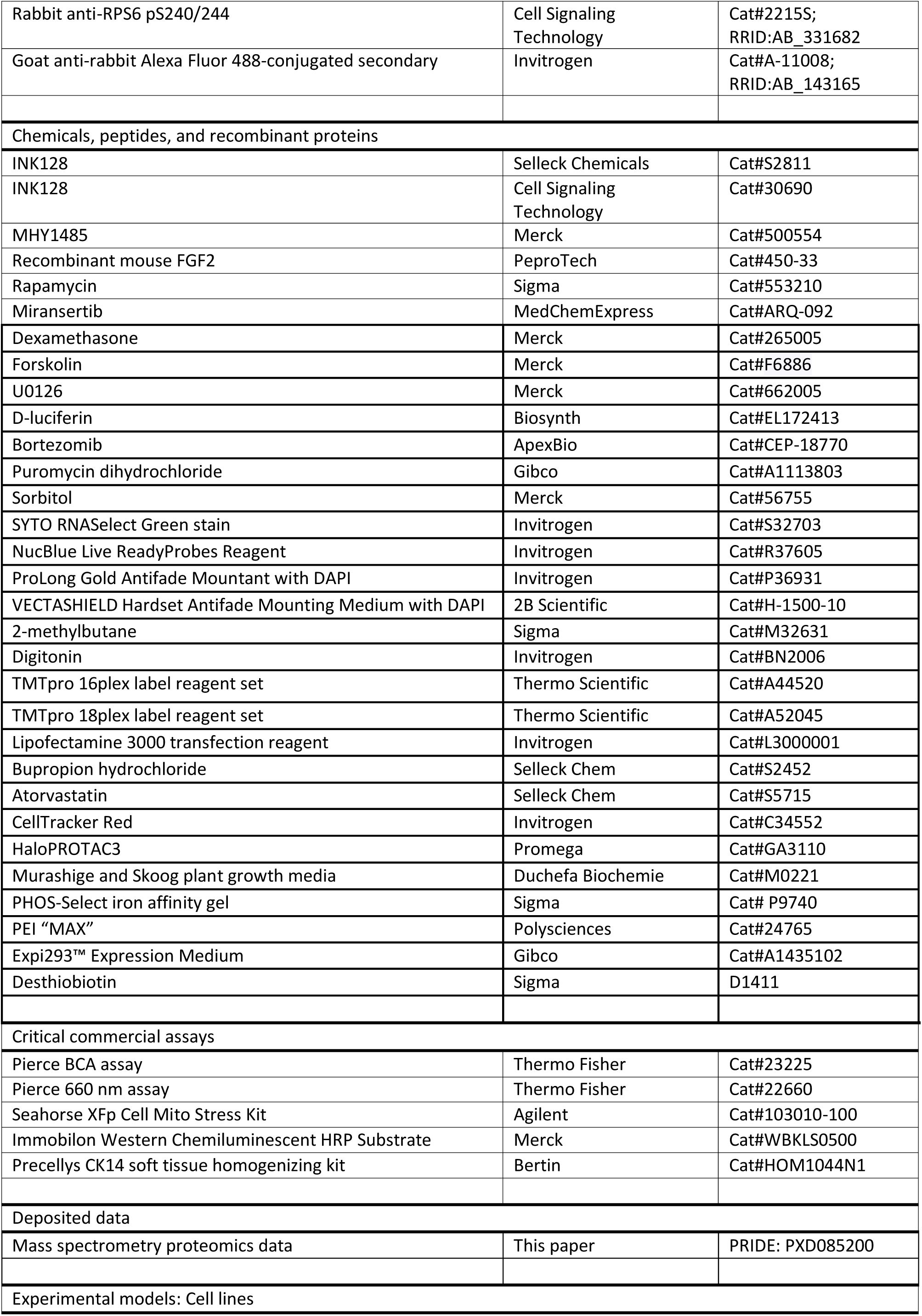
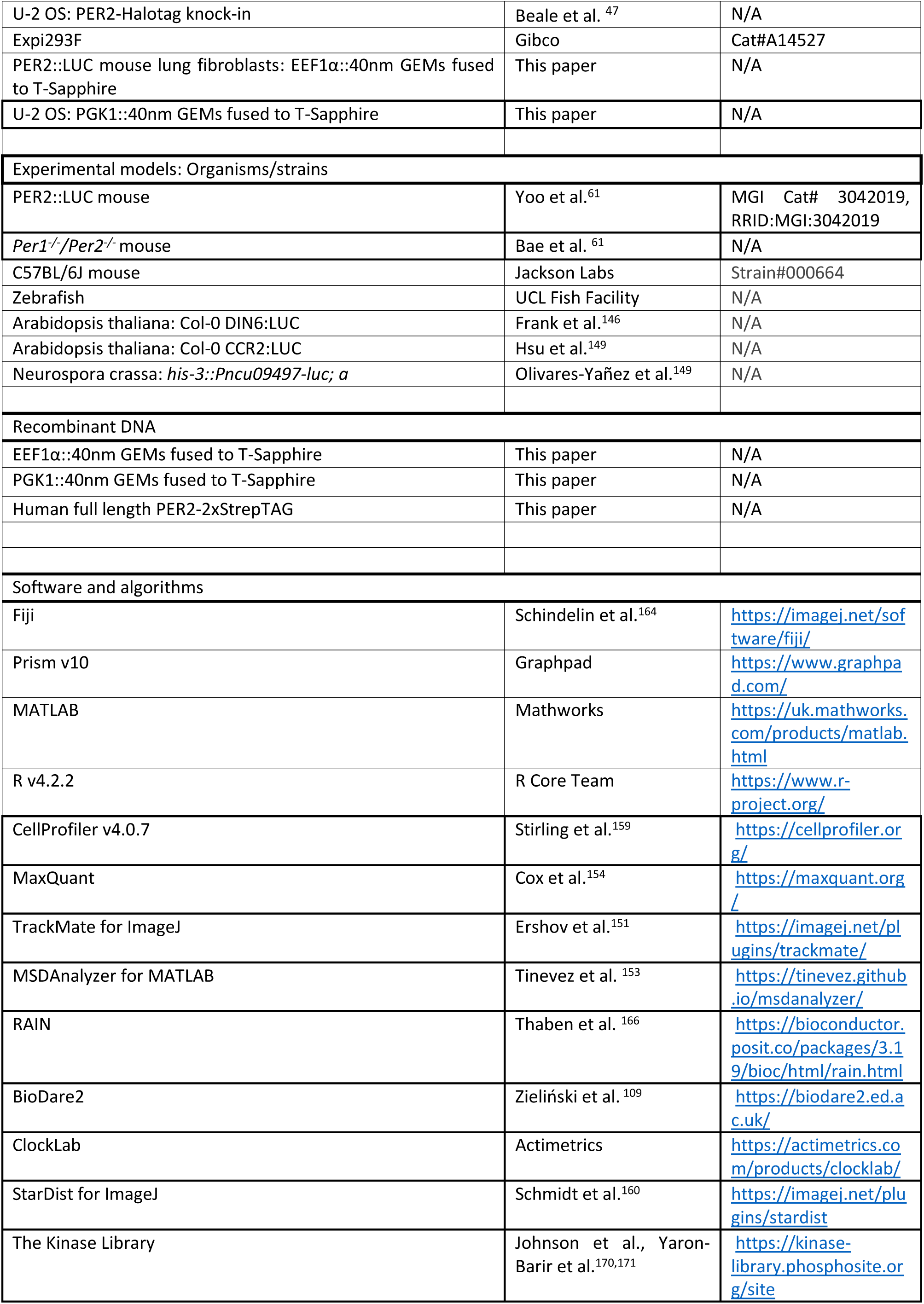
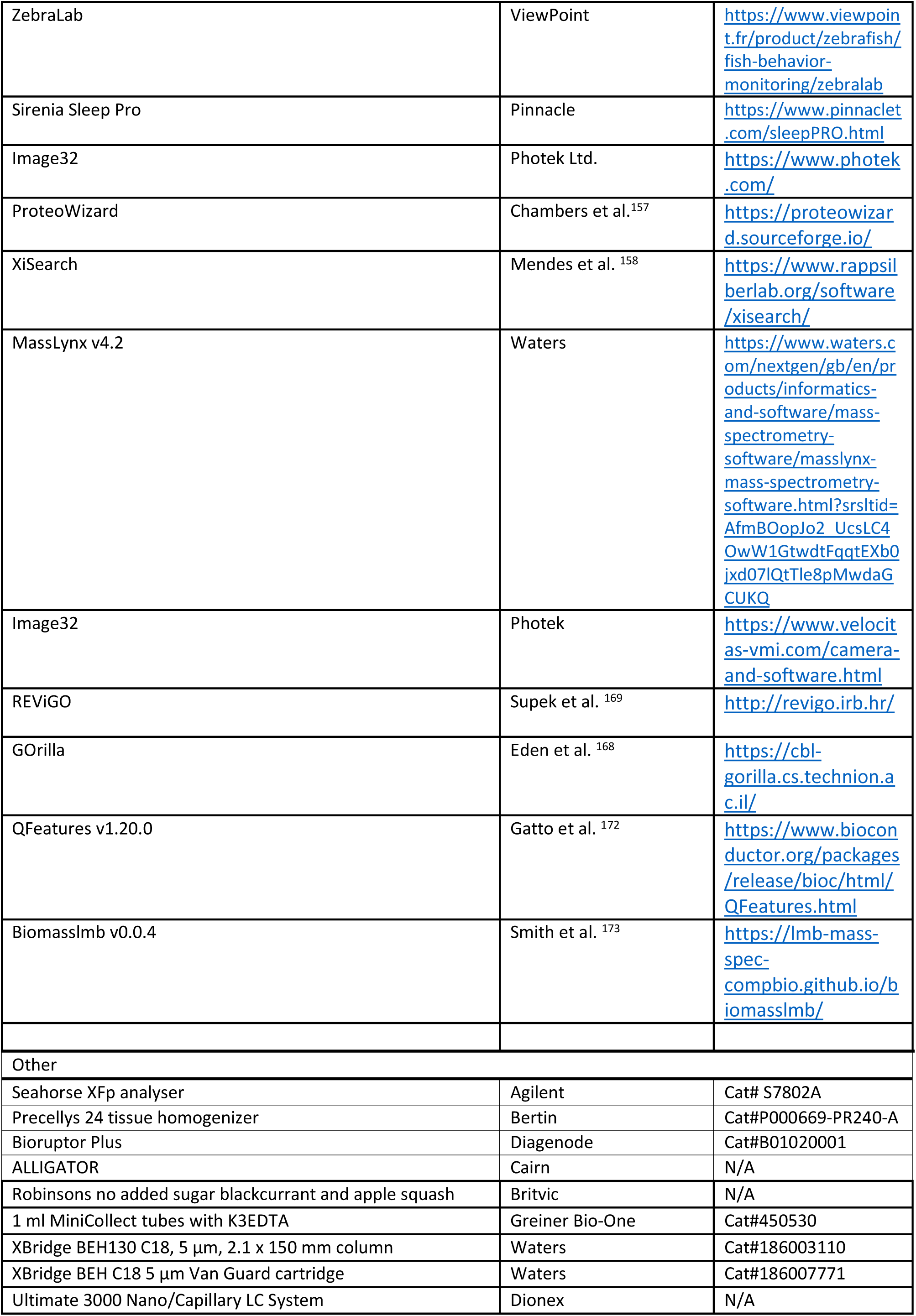
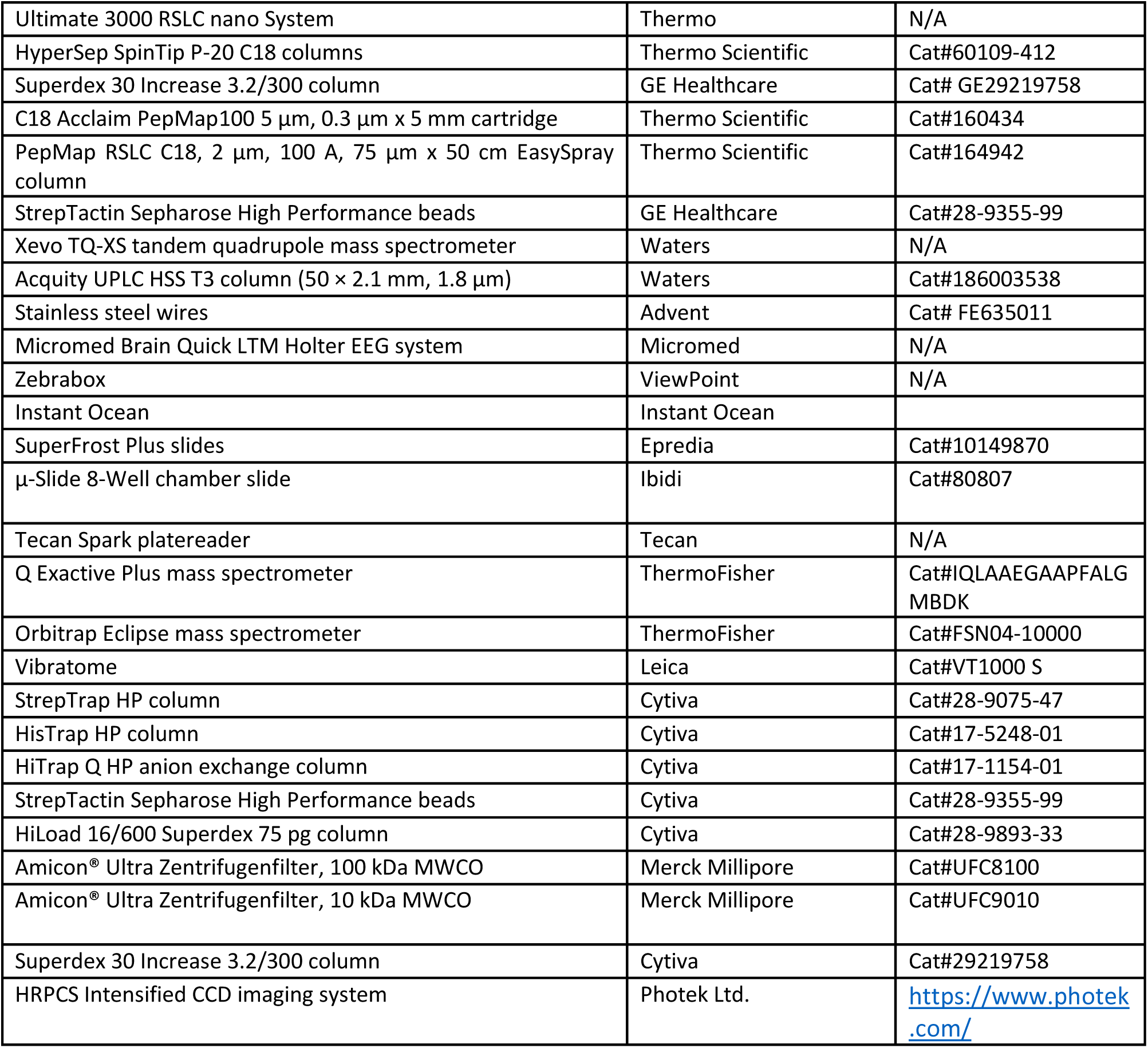

## Notes

### Summary of Updates

Minor edits to main text including abstract, introduction, discussion. Change of manuscript title.

