## Supplementary Figures for "mTOR signalling, not clock gene rhythms, organises daily physiology"

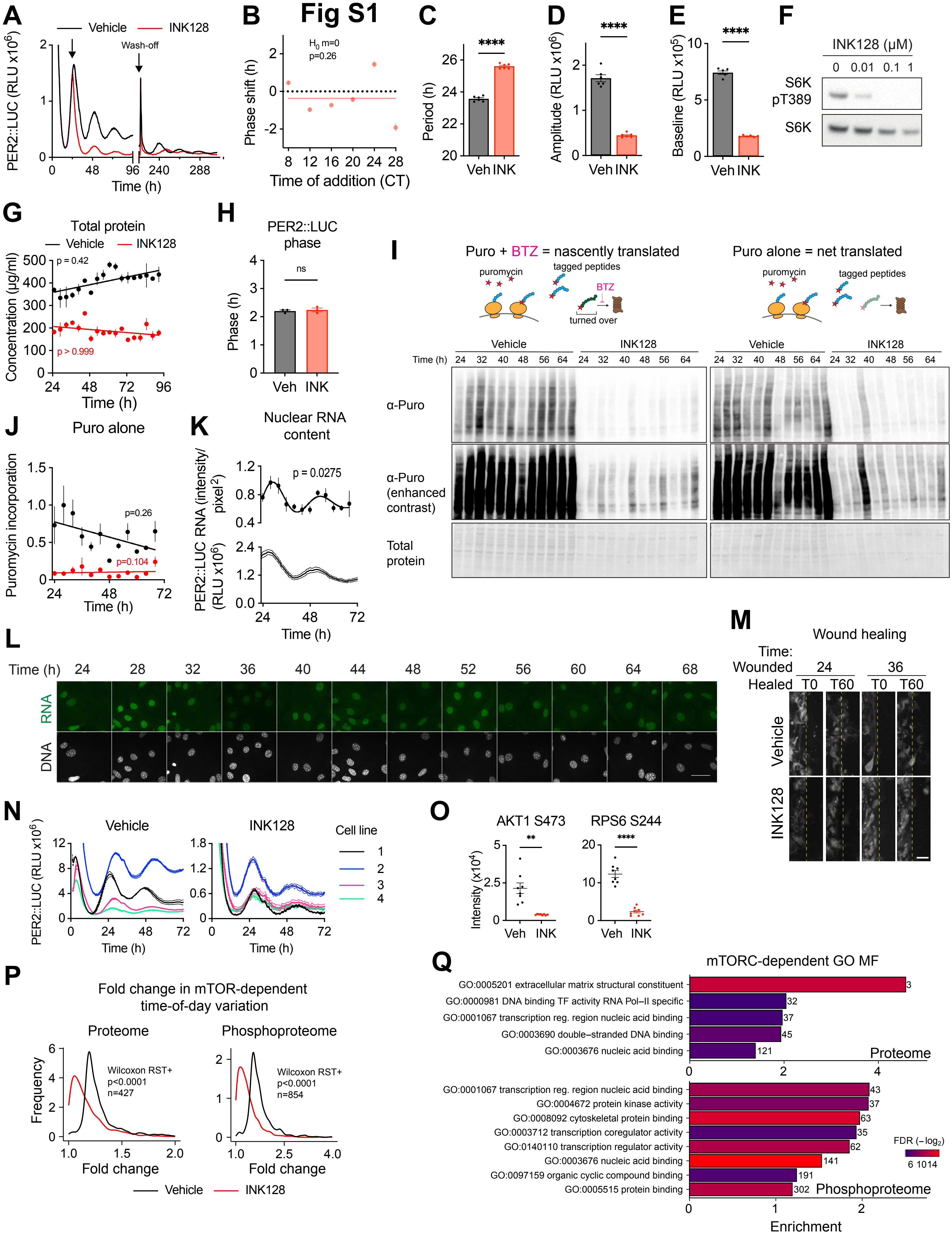

Fig S1
A
Vehicle
INK128
Wash-off
PER2::LUC (RLU x10^6)
Time (h)
B
H0 m=0
p=0.26
Phase shift (h)
Time of addition (CT)
C
Period (h)
Veh INK
D
Amplitude (RLU x10^6)
E
Baseline (RLU x10^5)
F
INK128 (µM)
0 0.01 0.1 1
S6K pT389
S6K
G
Total protein
Vehicle
INK128
p = 0.42
p > 0.999
Concentration (µg/ml)
Time (h)
H
PER2::LUC phase
ns
Phase (h)
Veh INK
I
Puro + BTZ = nascently translated
Puro alone = net translated
puromycin
tagged peptides
BTZ
turned over
Vehicle
INK128
Time (h)
α-Puro
α-Puro (enhanced contrast)
Total protein
J
Puro alone
p=0.26
p=0.104
Puromycin incorporation
Time (h)
K
Nuclear RNA content
p = 0.0275
PER2::LUC RNA (intensity/pixel^2)
(RLU x10^6)
Time (h)
L
Time (h)
24 28 32 36 40 44 48 52 56 60 64 68
RNA
DNA
M
Wound healing
Time:
Wounded 24 36
Healed T0 T60 T0 T60
Vehicle
INK128
N
Vehicle
INK128
Cell line
1
2
3
4
PER2::LUC (RLU x10^6)
Time (h)
O
AKT1 S473
RPS6 S244
Intensity (x10^4)
Veh INK
P
Fold change in mTOR-dependent time-of-day variation
Proteome
Phosphoproteome
Wilcoxon RST+
p<0.0001
n=427
Wilcoxon RST+
p<0.0001
n=854
Frequency
Fold change
Vehicle
INK128
Q
mTORC-dependent GO MF
GO:0005201 extracellular matrix structural constituent
GO:0000981 DNA binding TF activity RNA Pol–II specific
GO:0001067 transcription reg. region nucleic acid binding
GO:0003690 double–stranded DNA binding
GO:0003676 nucleic acid binding
Proteome
GO:0001067 transcription reg. region nucleic acid binding
GO:0004672 protein kinase activity
GO:0008092 cytoskeletal protein binding
GO:0003712 transcription coregulator activity
GO:0140110 transcription regulator activity
GO:0003676 nucleic acid binding
GO:0097159 organic cyclic compound binding
GO:0005515 protein binding
Phosphoproteome
FDR (−log2)
Enrichment

### Fig S2

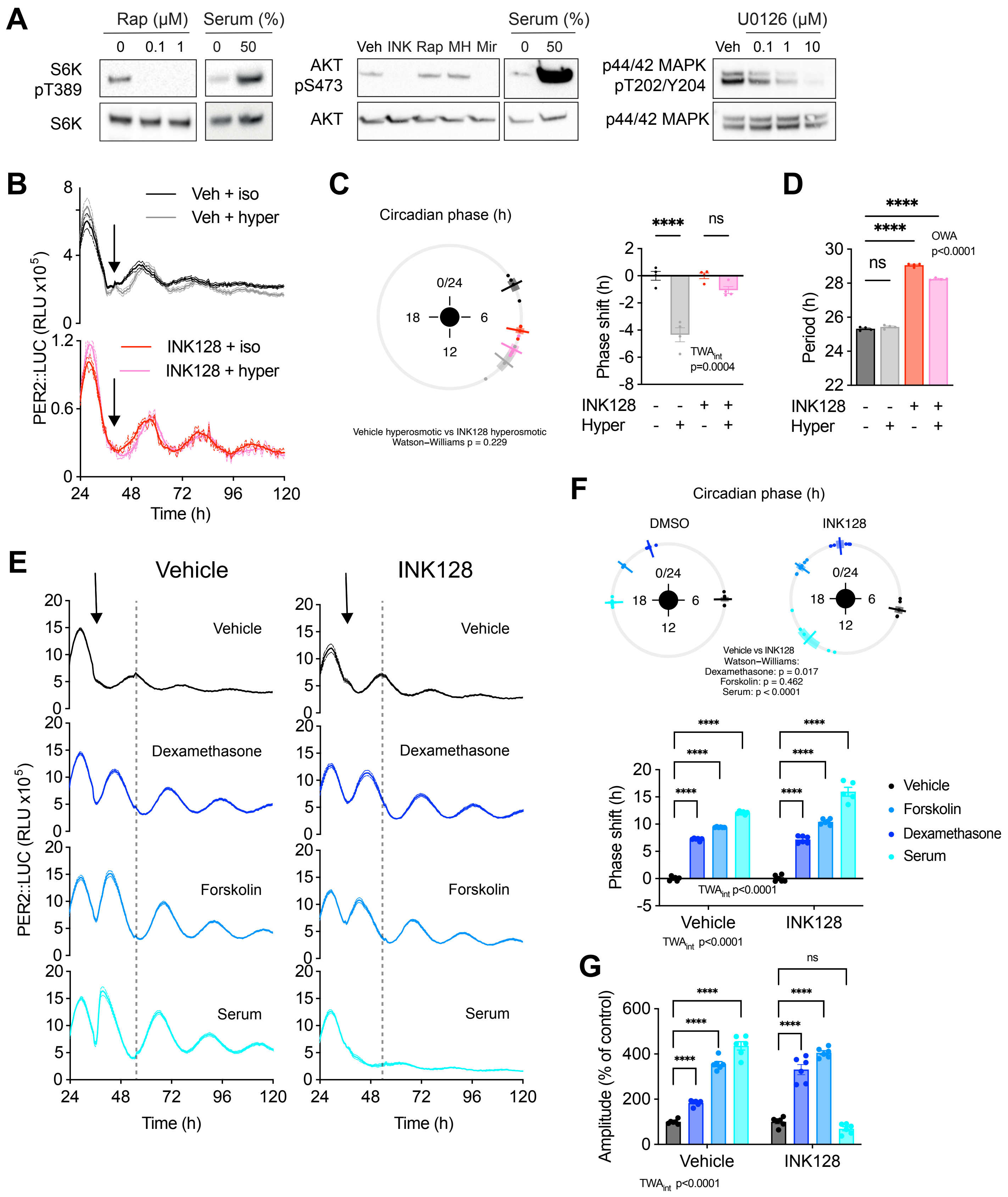
A
Rap (μM)
0 0.1 1
Serum (%)
0 50
S6K pT389
S6K
Veh INK Rap MH Mir
AKT pS473
AKT
U0126 (μM)
Veh 0.1 1 10
p44/42 MAPK pT202/Y204
p44/42 MAPK
B
Veh + iso
Veh + hyper
INK128 + iso
INK128 + hyper
PER2::LUC (RLU x10^5)
Time (h)
C
Circadian phase (h)
0/24
18
6
12
Vehicle hyperosmotic vs INK128 hyperosmotic
Watson–Williams p = 0.229
Phase shift (h)
TWAint p=0.0004
INK128
Hyper
D
Period (h)
OWA p<0.0001
ns
E
Vehicle
INK128
Dexamethasone
Forskolin
Serum
F
DMSO
Vehicle vs INK128
Watson–Williams:
Dexamethasone: p = 0.017
Forskolin: p = 0.462
Serum: p < 0.0001
TWAint p<0.0001
G
Amplitude (% of control)

### Fig S3

A

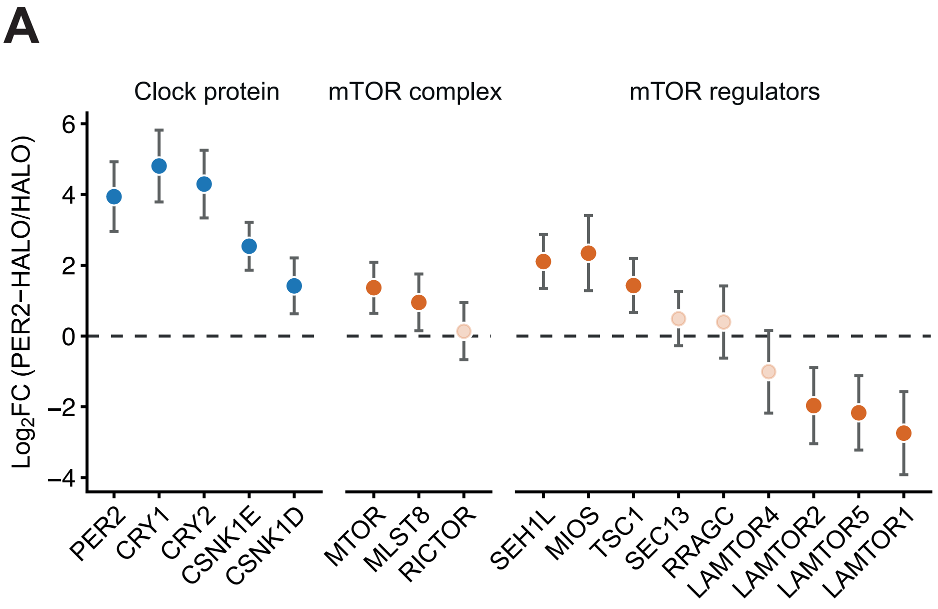


B

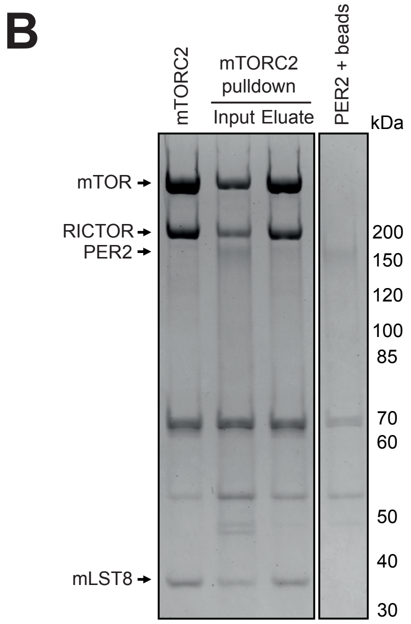


C

| Residue on PER2 | Crosslinked residue on PER peptide | Residue on mTOR | Crosslinked residue on mTOR peptide |
|---|---|---|---|
| 278 | K**S**HENEIR | 1316 | DLFNAAFVSCcmWSELN**E**DQQDELIR |
| 384 | **K**ILQSGGQPFDYSPIR | 1219 | GYTLA**D**EEEDPLIYQHR |
| 384 | **K**ILQSGGQPFDYSPIR | 1239 | SGQG**D**ALASGPVETGPMK |
| 384 | **K**ILQSGGQPFDYSPIR | 1515 | MAAAAAWGLGQWDSME**E**YTCcmMIPR |
| 436 | VGPLNEDVFAAHPCcmTE**E**K | 588 | TLGSFEFEGH**S**LTQFVR |
| 623 | **K**ATVSPGPHAGEAEPPSR | 1316 | DLFNAAFVSCcmWSELNEDQQ**D**ELIR |
| 623 | **K**ATVSPGPHAGEAEPPSR | 1515 | MAAAAAWGLGQWDSME**E**YTCcmMIPR |
| 722 | **K**LGLTK | 1316 | DLFNAAFVSCcmWSELNEDQQ**D**ELIR |

| Residue on PER2 | Crosslinked residue on PER peptide | Residue on mLST8 | Crosslinked residue on mLST8 peptide |
|---|---|---|---|
| 623 | KA**T**VSPGPHAGEAEPPSR | 87 | NIASVGFH**E**DGR |
| 623 | **K**ATVSPGPHAGEAEPPSR | 314 | AVVCcmLAFNDSVL**G** |
| 723 | LGLT**K**EVLAAHTQKEEQSFLQK | 314 | AVVCcmLAFNDSVL**G** |

| Residue on PER2 | Crosslinked residue on PER peptide | Residue on RAPTOR | Crosslinked residue on RAPTOR peptide |
|---|---|---|---|
| 593 | YLESCcmNEAATL**K**R | 1243 | MP**E**SVNVLQIVK |

D

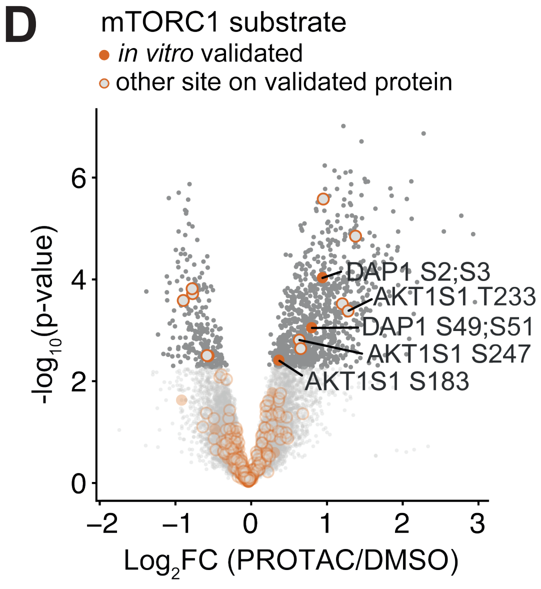

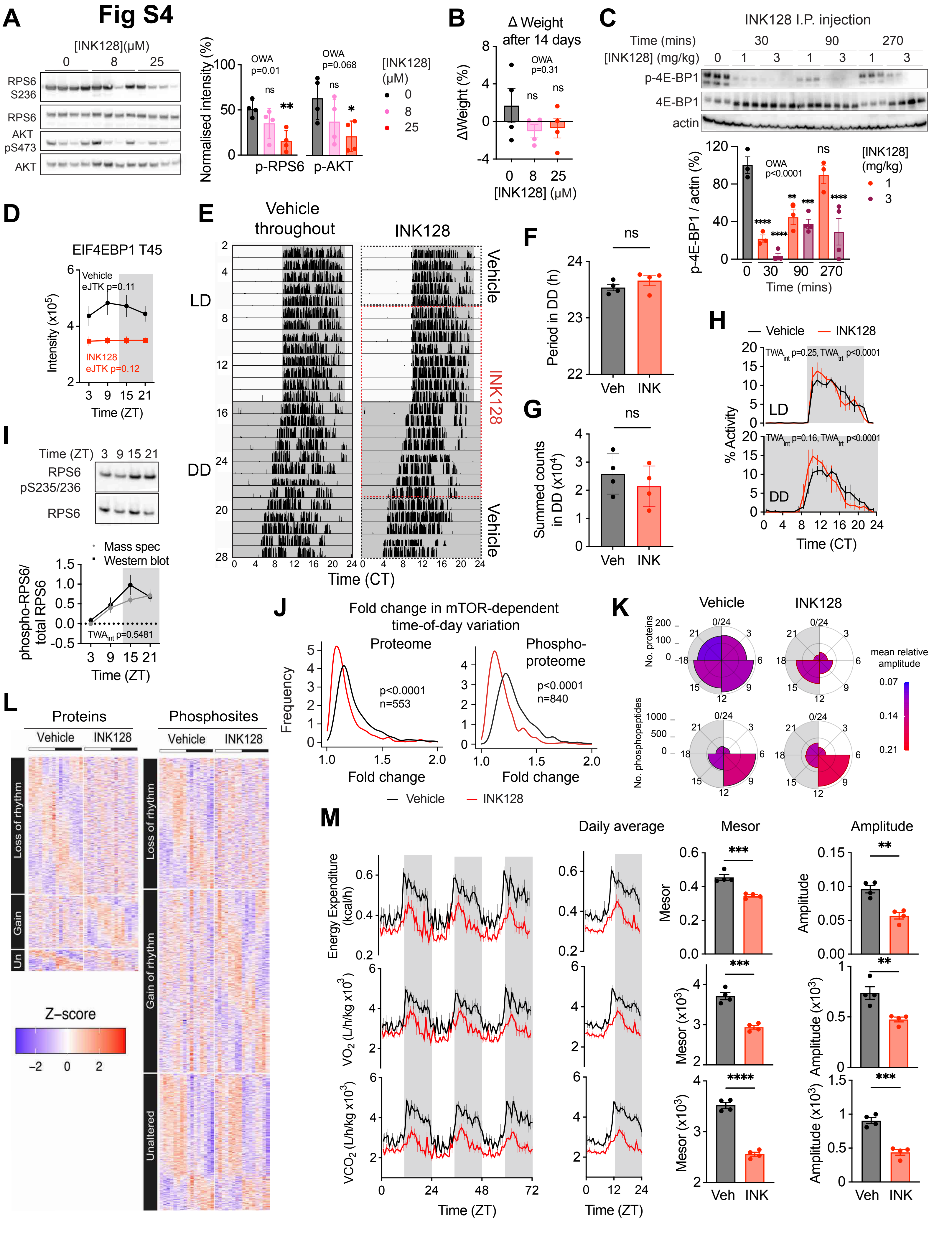
Fig S4
A
[INK128](μM)
0
8
25
RPS6 S236
RPS6
AKT pS473
AKT
Normalised intensity (%)
OWA p=0.01
OWA p=0.068
ns
**
*
p-RPS6
p-AKT
[INK128] (μM)
B
Δ Weight after 14 days
ΔWeight (%)
OWA p=0.31
[INK128] (μM)
C
INK128 I.P. injection
Time (mins)
[INK128] (mg/kg)
p-4E-BP1
4E-BP1
actin
p-4E-BP1 / actin (%)
OWA p<0.0001
[INK128] (mg/kg)
Time (mins)
D
EIF4EBP1 T45
Vehicle eJTK p=0.11
INK128 eJTK p=0.12
Intensity (x10^5)
Time (ZT)
E
Vehicle throughout
INK128
LD
DD
Vehicle
Time (CT)
F
Period in DD (h)
ns
Veh
INK
G
Summed counts in DD (x10^4)
H
Vehicle
INK128
TWA int p=0.25, TWA trt p<0.0001
TWA int p=0.16, TWA trt p<0.0001
% Activity
Time (CT)
I
Time (ZT)
RPS6 pS235/236
RPS6
Mass spec
Western blot
phospho-RPS6/ total RPS6
TWA int p=0.5481
J
Fold change in mTOR-dependent time-of-day variation
Proteome
Phospho-proteome
p<0.0001 n=553
p<0.0001 n=840
Frequency
Fold change
K
No. proteins
No. phosphopeptides
mean relative amplitude
L
Proteins
Phosphosites
Loss of rhythm
Gain
Un
Gain of rhythm
Unaltered
Z−score
M
Daily average
Mesor
Amplitude
Energy Expenditure (kcal/h)
VO2 (L/h/kg x10^3)
VCO2 (L/h/kg x10^3)
Mesor (x10^3)
Amplitude (x10^3)
Time (ZT)

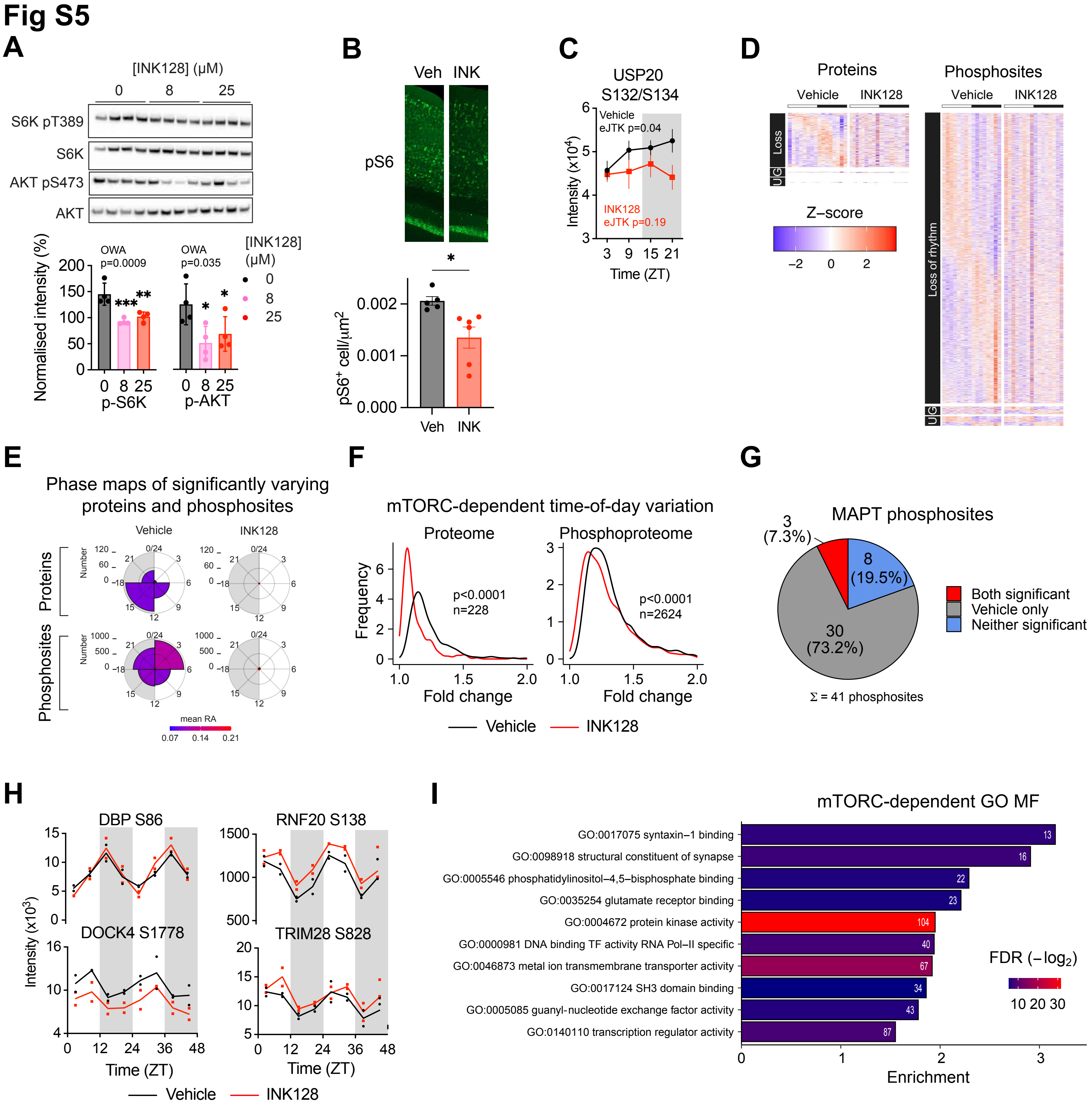

Fig S5
A
[INK128] (μM)
0
8
25
S6K pT389
S6K
AKT pS473
AKT
Normalised intensity (%)
OWA
p=0.0009
OWA
p=0.035
[INK128] (μM)
0
8
25
p-S6K
p-AKT
B
Veh
INK
pS6
pS6+ cell/μm2
C
USP20 S132/S134
Vehicle
eJTK p=0.04
INK128
eJTK p=0.19
Intensity (x10⁴)
Time (ZT)
D
Proteins
Phosphosites
Vehicle
INK128
Loss
Loss of rhythm
Z−score
E
Phase maps of significantly varying proteins and phosphosites
Vehicle
INK128
Proteins
Phosphosites
Number
mean RA
0.07
0.14
0.21
F
mTORC-dependent time-of-day variation
Proteome
Phosphoproteome
Frequency
p<0.0001
n=228
p<0.0001
n=2624
Fold change
Vehicle
INK128
G
MAPT phosphosites
3 (7.3%)
8 (19.5%)
30 (73.2%)
Both significant
Vehicle only
Neither significant
Σ = 41 phosphosites
H
DBP S86
RNF20 S138
DOCK4 S1778
TRIM28 S828
Intensity (x10³)
Time (ZT)
Vehicle
INK128
I
mTORC-dependent GO MF
GO:0017075 syntaxin−1 binding
GO:0098918 structural constituent of synapse
GO:0005546 phosphatidylinositol−4,5−bisphosphate binding
GO:0035254 glutamate receptor binding
GO:0004672 protein kinase activity
GO:0000981 DNA binding TF activity RNA Pol−II specific
GO:0046873 metal ion transmembrane transporter activity
GO:0017124 SH3 domain binding
GO:0005085 guanyl-nucleotide exchange factor activity
GO:0140110 transcription regulator activity
FDR (−log2)
10 20 30
Enrichment

### Fig S6

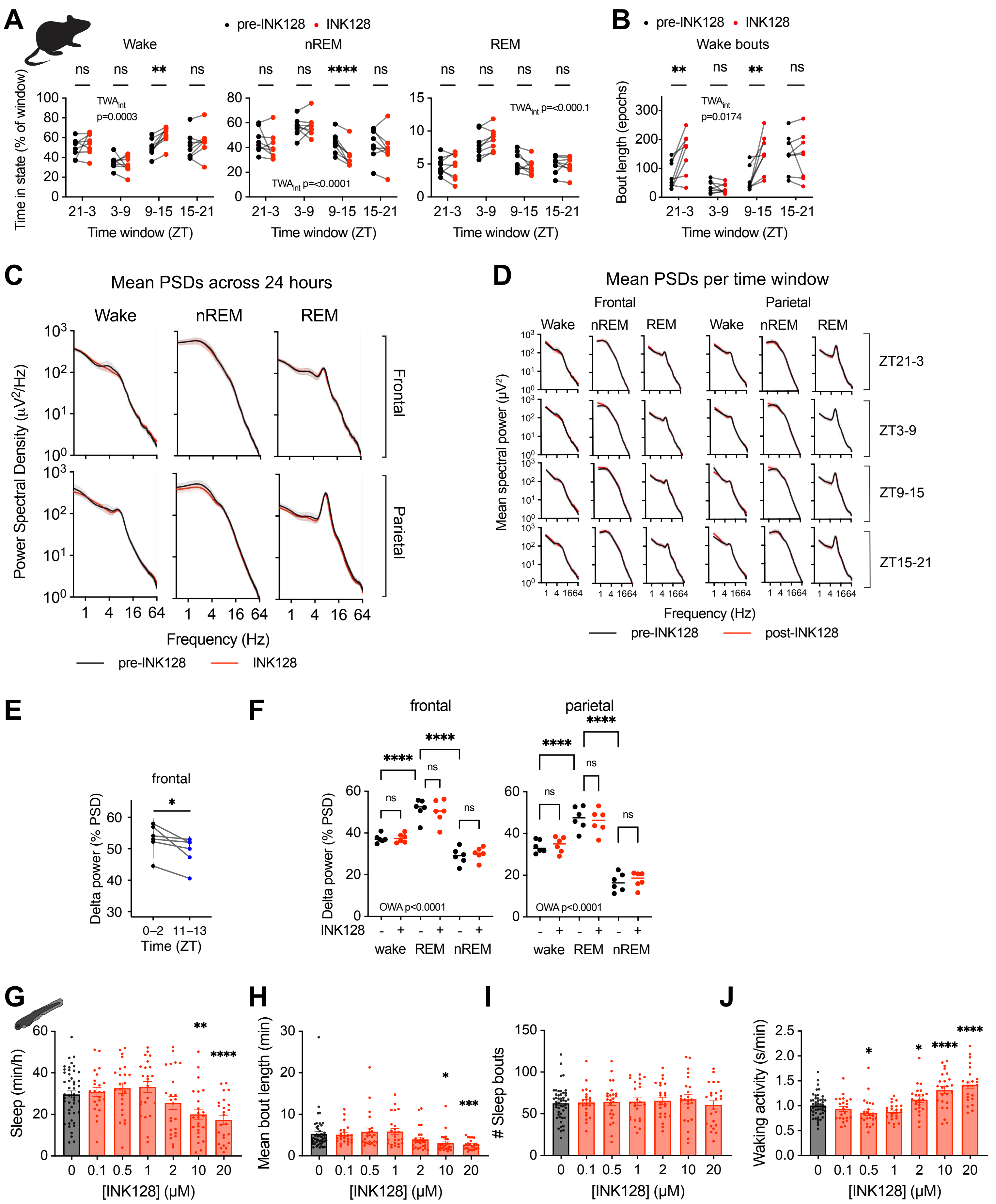

A
pre-INK128 • INK128
Wake
nREM
REM
ns ns ** ns
ns ns **** ns
ns ns ns ns
TWAint p=0.0003
TWAint p=<0.0001
TWAint p=<0.000.1
Time in state (% of window)
21-3 3-9 9-15 15-21
Time window (ZT)
B
pre-INK128 • INK128
Wake bouts
** ns ** ns
TWAint p=0.0174
Bout length (epochs)
21-3 3-9 9-15 15-21
Time window (ZT)
C
Mean PSDs across 24 hours
Wake
nREM
REM
Frontal
Parietal
Power Spectral Density (μV²/Hz)
1 4 16 64
Frequency (Hz)
pre-INK128
INK128
D
Mean PSDs per time window
Frontal
Parietal
Wake nREM REM Wake nREM REM
ZT21-3
ZT3-9
ZT9-15
ZT15-21
Mean spectral power (μV²)
1 4 1664
Frequency (Hz)
pre-INK128
post-INK128
E
frontal
*
Delta power (% PSD)
0–2 11–13
Time (ZT)
F
frontal
parietal
****
****
ns
ns
ns
OWA p<0.0001
Delta power (% PSD)
INK128 - + - + - +
wake REM nREM
G
** ****
Sleep (min/h)
0 0.1 0.5 1 2 10 20
[INK128] (μM)
H
* ***
Mean bout length (min)
0 0.1 0.5 1 2 10 20
[INK128] (μM)
I
### Sleep bouts
0 0.1 0.5 1 2 10 20
[INK128] (μM)
J
* * **** ****
Waking activity (s/min)
0 0.1 0.5 1 2 10 20
[INK128] (μM)

### Fig S7

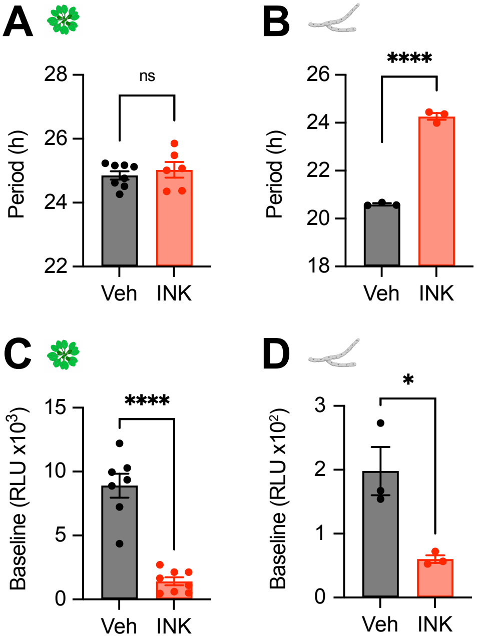

A
Period (h)
28
26
24
22
ns
Veh
INK
B
Period (h)
26
24
22
20
18
****
Veh
INK
C
Baseline (RLU x10³)
15
10
5
0
****
Veh
INK
D
Baseline (RLU x10²)
3
2
1
0
*
Veh
INK
